# Unblocking genome editing of major animal mycoplasmas using CRISPR/Cas9 base editor systems

**DOI:** 10.1101/2022.03.09.483585

**Authors:** Thomas Ipoutcha, Fabien Rideau, Geraldine Gourgues, Yonathan Arfi, Carole Lartigue, Alain Blanchard, Pascal Sirand-Pugnet

**Author notes:** **Corresponding author:** Pascal Sirand-Pugnet, Address: Univ. Bordeaux, INRAE, UMR BFP, 71 avenue Edouard Bourlaux, 33140 Villenave d’Ornon, France. The authors declare no conflict of interest.

## Abstract

Mycoplasmas are minimal bacteria that infect humans, wildlife, and most economically important livestock species. Mycoplasma infections cause a large range of chronic inflammatory diseases, eventually leading to death in some animals. Due to the lack of efficient recombination and genome engineering tools, the production of mutant strains for the identification of virulence factors and the development of improved vaccine strains is still a bottleneck for many mycoplasma species. Here, we demonstrate the efficacy of a CRISPR-derived genetic tool to introduce targeted mutations in three major pathogenic species that span the phylogenetic diversity of these bacteria: the avian pathogen *Mycoplasma gallisepticum* and the two most important bovine mycoplasmas, *Mycoplasma bovis* and *Mycoplasma mycoides* subsp. *mycoides*. As a proof of concept, we successfully used an inducible dCas9-cytidine deaminase system to disrupt several major virulence factors in these pathogens. Various induction times and inducer concentrations were evaluated to optimize editing efficiency. The optimized system was sufficiently powerful to disrupt 54 of 55 insertion sequence transposases in a single step. Whole genome sequencing showed that off-target mutations were limited and suggest that most variations detected in the edited genomes are Cas9-independent. This effective, rapid, and easy-to-use genetic tool opens a new avenue for the study of these important animal pathogens and, most likely, the entire class *Mollicutes*.

**Significance:** Mycoplasmas are minimal wall-less pathogenic bacteria that infect a wide range of hosts, including humans, livestock, and wild animals. Major pathogenic species cause acute to chronic infections involving still poorly characterized virulence factors. The lack of precise genome editing tools has hampered functional studies for many species, leaving multiple questions about the molecular basis of their pathogenicity unanswered. We developed a CRISPR-derived base editor for three major pathogenic species, *Mycoplasma gallisepticum*, *Mycoplasma bovis*, and *Mycoplasma mycoides* subsp*. mycoides*. Several virulence factors were successfully targeted and we were able to edit up to 54 target sites in a single step. The availability of this efficient and easy-to-use genetic tool will greatly facilitate functional studies in these economically important bacteria.

## Introduction

Mycoplasmas are minimal pathogens that belong to the class *Mollicutes* (1). They are characterized by a streamlining evolution from a Gram-positive ancestor, marked by drastic genome reduction (2–4). This evolution has led to the loss of diverse cellular functions including cell-wall production, various metabolic pathways and efficient recombination machinery (5). These minimal bacteria are found in a wide range of animals, including humans and livestock. Many mycoplasmas are etiological agents of diseases that considerably reduce animal production yields and inflate veterinary healthcare costs (3, 6, 7). Currently, three of the most prevalent and economically relevant species worldwide are *Mycoplasma gallisepticum* (*Mgal*), *Mycoplasma bovis* (*Mbov*), and *Mycoplasma mycoides* subsp *mycoides* (*Mmm*). *Mgal* is an animal pathogen listed by the World Organization for Animal Health (OIE) and responsible for respiratory disease in poultry farms worldwide (8–10). *Mbov* is responsible for a bovine respiratory disease (BRD), as well as mastitis and reproductive disease (11, 12). *Mmm* is a member of the so-called mycoides cluster, which groups five significant ruminant pathogens. It is the causative agent of contagious bovine pleuropneumonia (CBPP), an OIE-listed disease that can take chronic, acute, or hyperacute forms. In the hyperacute form, CBPP symptoms include pericardial effusion, fever, and death (13). The control strategies for mycoplasmoses vary geographically and include prophylaxis and the use of antibiotics, vaccines, and ultimately, culling. In the context of increasing antibiotic resistance (14) and the relative lack of effective vaccines, better knowledge of the molecular basis of virulence and host-pathogen interactions is required to improve curative protocols and design more efficient vaccines (13, 15). However, the lack of efficient genome engineering tools for many mycoplasmas, including the three species listed above, limits functional genomics approaches and hinders efforts toward the production of rationally designed vaccines. During the past decade, cutting-edge genome engineering methods have been developed that rely on the cloning of the mycoplasma genome in yeast before editing with various genetic tools and back transplantation into a recipient cell (16, 17). However, although offering unmatched possibilities to investigate and redesign complete genomes, these in-yeast approaches can be difficult to adapt to new species. They are still restricted to a small number of mycoplasmas and have not been developed for any of the three pathogens considered here. Indeed, efforts to adapt these synthetic biology approaches to *Mmm* have been thus far unsuccessful, whereas they are now available for all other members of the mycoides cluster (18). Only replicative (*oriC*) plasmids and transposon-based mutagenesis are currently available for *Mbov*, and *Mmm* (19, 20). For *Mgal*, a first tool for targeted homologous recombination of short genomic regions has recently been reported (21).

Since 2012, CRISPR-based genetic tools have revolutionized the field of genome engineering in eukaryotes and prokaryotes. The typical CRISPR/Cas9 tool is based on the Cas9 nuclease, guided to specific loci by single guide RNAs (sgRNAs) (22). Following DNA cleavage, repair occurs through distinct cellular mechanisms, including non-homologous end joining (NHEJ) and homology-directed recombination (HDR). However, many bacteria, including mycoplasmas, lack efficient NHEJ and HDR repair machineries. Therefore, the CRISPR/Cas9 tools can only be used as a counter-selection method (23). Base-editor systems (BEs) have offered a means to overcome this problem. Indeed, BEs combine an inactivated form of Cas9 (dCas9) fused with a cytosine deaminase (CBE) or an adenosine deaminase (ABE) and an uracil glycosylase inhibitor (UGI). At the molecular level, the mechanism of action of a CBE occurs as follows (Figure S1): (1) An R-loop is opened in the target site by the dCas9-sgRNA complex, (2) the single strand DNA is accessible to the fused deaminase within a specific editing window, and (3) the CBE catalyzes the deamination of cytosine into uracil, which is recognized as thymine after replication. ABE converts adenine into inosil, which is recognized as guanine after replication (24–26). Base excision repair on edited nucleotides is prevented by UGI, improving the global editing efficiency (27). Therefore, CBE and ABE induce C:G to T:A and A:T to G:C transitions, respectively (24, 28).

Here, we demonstrate the functionality of a base editor system in *Mgal* using a transposon to introduce the CBE and sgRNA into the bacterial cells. After optimization, three virulence genes located at various positions on the *Mgal* chromosome were successfully disrupted and individual mutants were isolated. Next, a replicative plasmid and another transposon were used as vectors to evaluate this CBE in *Mbov* and *Mmm,* demonstrating the high efficiency of the system for single-gene targeting in ruminant pathogens. Finally, multi-target mutagenesis using a single sgRNA was demonstrated by successfully disrupting 54 insertion sequences in *Mmm*.

## Results

### Design and construction of a base-editor genetic tool for mycoplasma

Two different cytosine deaminases commonly used for base editing, rAPOBEC1 from *Rattus norvegicus* (24) and pmcDA1 from *Petromyzan marinus* (24, 25) (Figure 1A, SI-2), were evaluated as editing tools in mycoplasmas. The coding sequences of these two proteins were codon optimized, chemically synthesized, and fused to an inactive version (dead) of Cas9 (dCas9) from *Streptococcus pyogenes* (SpdCas9). rAPOBEC1 was fused with the N-terminus of SpdCas9 using the a 16 amino acid-linker XTEN (29). By contrast, pmcDA1 was fused to the C-terminus of SpdCas9 with a 36 amino acid-linker (30). The uracil-glycosylase inhibitor from *Bacillus subtilis* phage (29, 31) was also fused to the C-terminus of the resulting hybrid protein (Figure S1) and used to block potential base excision repair activity of the bacterial cell. The tetracycline-inducible promoter P*xyl/tetO2* (32) was used to control the expression of these two hybrid proteins. The PS promoter, which is considered to be constitutive in many mollicutes species, was used to control the expression of the sgRNA (Figure S2A). All genetic elements were assembled within a Tn*4001*-derived transposon (33), resulting in plasmids pTI4.0_SpdCas9_pmcDA1 (Figure S2B) and pTI4.0_SpdCas9_rAPOBEC1 (Figure S3A).

**Figure 1.**
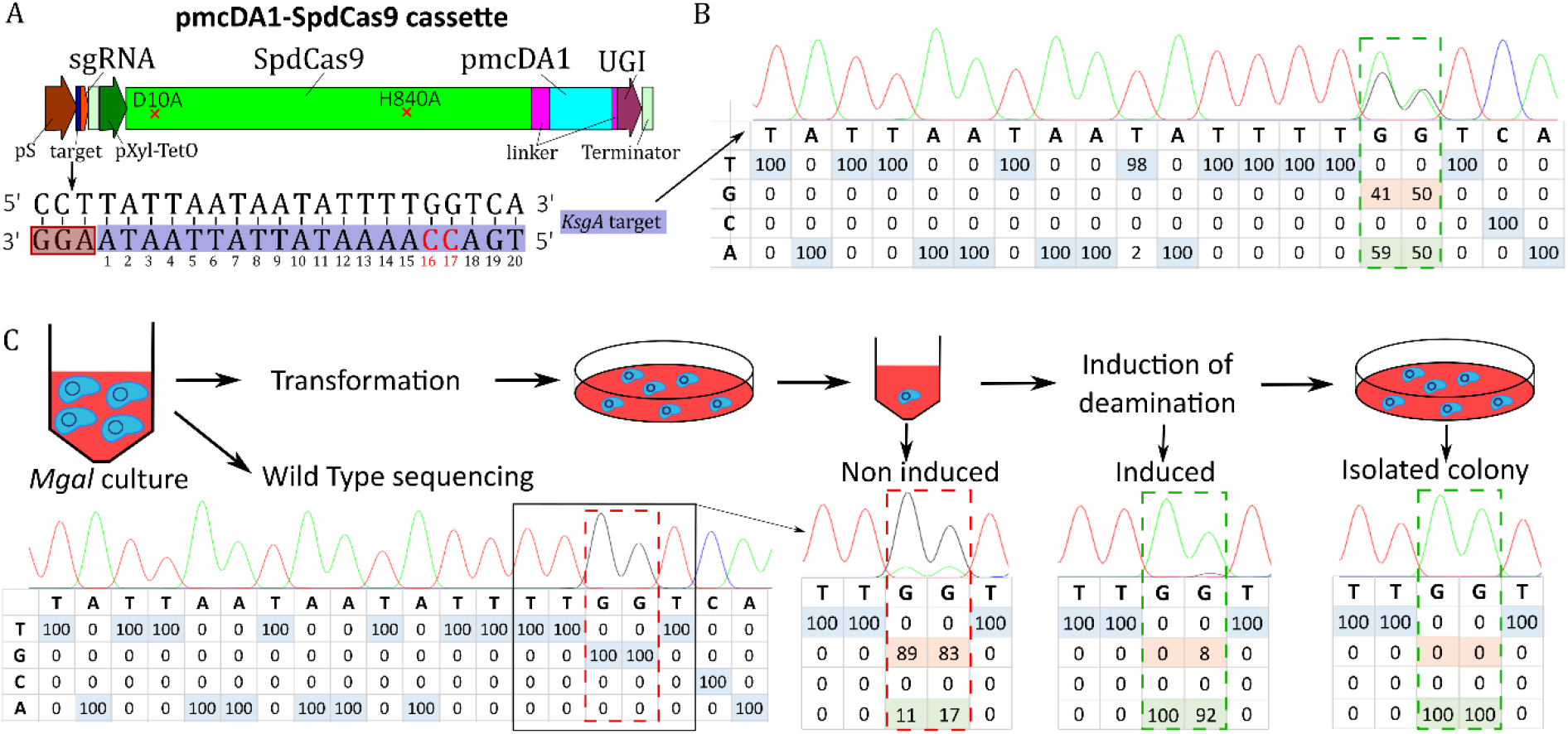
Targeting of the *ksgA* gene in *Mgal* using the pmcDA1 deaminase-encoding plasmid. **A-** Scheme of the CBE expression cassette. The synthetic cassette is composed of (1) a sgRNA (red), including the 20-nucleotide target spacer (dark blue) under the control of the spiralin promoter (brown), and (2) the codon-optimized hybrid protein composed of *S. pyogenes* dead Cas9 (SpdCas9, green), linkers (purple), the pmcDA1 deaminase protein (light blue), and UGI (uracil glycosylase inhibitor, dark purple), under the control of the P*xyl/tetO2* inducible promoter (dark green). Fibril terminators from *S. citri* (light green) were added downstream of the sgRNA and hybrid protein expression cassettes. The sequence of the *ksgA* target is represented here. Cytosine residues that are susceptible to deamination are colored in red. The positions of bases in the target are indicated below each base, with numbering starting at the PAM sequence. **B-**. The percentage of bases found in the population of transformants were determined by Sanger sequencing of complementary strand. The chromatograms were analyzed using EditR software and are represented in the table for each nucleotide position in the target sequence. Positions 16 and 17 are framed in a dotted rectangle. **C-** Schematic representation of the base-editing experiment and indication of various checkpoints (Sanger results and EditR analysis) for screening until obtaining isolated mutants.

We selected the *ksgA* gene of *Mgal* as a first target to evaluate the ability of these recombinant CBEs to produce mutations in mycoplasma. *ksgA* is a non-essential gene that encodes an RNA methyltransferase that renders the bacteria sensitive to kasugamycin (34). A 20-nucleotide sequence (5‘-TGA**<u>CC</u>**AAAATATTATTAATA-3’), upstream of the SpdCas9 compatible PAM sequence AGG (Figure 1A–B), was chosen as a target for *ksgA* inactivation. On the basis of previous reports, the two cytosines at positions 16 and 17 upstream of the PAM sequence should be in the theoretical editing window of the two CBE systems (35). Deamination of C_17_ is expected to change a Glutamine CAA codon into a TAA stop codon. After transformation of *Mgal* with either pTI4.0_SpdCas9_pmcDA1 or pTI4.0_SpdCas9_rAPOBEC1 and induction with 0.5 μg.mL^−1^ anhydrotetracycline (aTC) in growth medium (23), we analyzed the impact of the CBEs on cytosine deamination within the target site by PCR and Sanger sequencing on the global population of transformants (Figure 1A–B, Figure S3A). We observed only limited base editing for the two cytosines at positions 16 and 17 (2%) for the pTI4.0_SpdCas9_rAPOBEC1 transformants (Figure S3), whereas base editing was highly prevalent (59% of C to T conversion at cytosine 16, 50% at cytosine 17) for the pTI4.0_SpdCas9_pmcDA1 transformants (Figure 1). There was no conversion in the control without the sgRNA. Thus, the deaminase pmcDA1 can be used to design a CBE that is functional in *Mgal.*

### Optimization of the CBE system and isolation of *ksgA* mutants in *Mgal*

Given our initial results, the CBE system based on the pmcDA1 cytosine deaminase (pTI4.0_spdCas9_pmcDA1) was selected for further optimization. First, we determined the optimal inducer concentration by assessing the deamination level of targeted cytosines using the same experimental strategy and *ksgA* as a target after overnight induction with 0.1, 0.25, 0.5, 1, 2.5, or 5 μg.mL^−1^ aTC (Figure S4A). There was notable leakage of expression in the absence of the inducer, with up to 25% conversion at position C_16_. In the presence of the inducer, the optimal concentration for base-editing was in the range of 0.1 to 0.5 μg.mL^−1^ aTC (> 50% efficiency) (Figure S4A). There was no detectable growth defect of *Mgal* with up to 1 μg.mL^−1^ aTC. However, the addition of 5 μg.mL^−1^ aTC resulted in a marked reduction in growth, as shown by the almost complete absence of a pH shift of the broth medium after aTC induction (pH = 6.6 at induction versus 6.54, 5.41, and 5.3 after overnight induction with 5 μg.mL^−1^, 0.5 μg.mL^−1^, and no inducer, respectively). This result suggests that aTC is toxic for *Mgal* at high concentrations. The use of freshly prepared (< 24 h) inducer was also necessary for maximum efficiency of base editing (data not shown).

Then, we studied the time of induction required for maximum efficiency of the CBE system. Inducer concentrations used for this experiment were 0.25 and 0.5 μg.mL^−1^ and base conversion was determined at two time points: after a 2 h or overnight induction (Figure S4B). After 2 h, significant base conversion was already evident relative to the non-induced condition, with up to 30% of base conversion at the two cytosine positions. However, overnight induction resulted in the highest efficiency, with two-fold higher conversion than after 2 h of induction (approximately 60%). Thus, the best induction conditions for maximum efficiency of the mycoplasma CBE in *Mgal* were 0.1 to 0.5 μg.mL^−1^ aTC and overnight incubation.

Finally, we performed a third experiment using the newly defined conditions to obtain isolated *Mgal* mutants using the pTI4.0_SpdCas9_pmcDA1 construct (Figure 1C). Targeted *ksgA* sites were analyzed at four time points: before transformation (wildtype), after transformation, after induction, and on isolated colonies selected on puromycin selective plates. In non-induced condition, 11% of C_16_ and 17% at C_17_ were edited (Figure 1C), indicating leakage of the inducible promoter, as previously shown. After induction, deamination in the edited population was nearly complete, with 100% conversion of C_16_ and 92% conversion of C_17_ to thymine. Further screening of five isolated colonies confirmed that four carried mutations at the two targeted cytosines, whereas the last one showed an incomplete deamination profile suggesting deamination occurred during clonal expansion. Resistance to kasugamycin was confirmed by plating an isolated mutant on a Hayflick plate supplemented, or not, with the antibiotic (Figure S5). Thus, the pmcDA1-based CBE tool promotes base conversion and gene inactivation in *Mgal* with high efficiency.

### Determination of the editing window for maximum CBE efficiency in *Mgal*

We investigated the base-editing window of the CBE system by targeting three other *Mgal* genes encoding virulence factors: *crmA* (GCW_RS01080), *gapA* (GCW_RS01075), and *cysP* (GCW_RS01695) (Figure 2). The *crmA* and *gapA* genes encode two primary cytoadherence proteins (36, 37). The cysteine protease encoded by *cysP* has been shown to cleave chicken immunoglobulin G (38). In the *crmA* target (Figure 2A) (5’-ATT**<u>C</u>**_17_AATAT**<u>C</u>**_11_**<u>C</u>**_10_TT**<u>C</u>**_7_GTTGTT/TGG-3’), cytosine residues C_7_, C_10_, C_11_, and C_17_ were potential deamination sites. After induction, C_17_ was the only base to be deaminated: 57% at the population level before the second plating (Figure 1C). Screening of isolated colonies resulted in 5 of 10 clones showing a C_17_ to T_17_ transition (Figure 2A, *crmA* edited). In the *gapA* target (Figure 2B) (5’-TGAA**<u>C</u>**_16_A**<u>C</u>**_14_AAGGTT**<u>C</u>**_7_TG**<u>C</u>**_4_TAA/CGG-3’), C_4_, C_7_, C_14_, and C_16_ were potential deamination sites. After induction, C_16_ was the sole deaminated position: 54% conversion estimated in the transformant population. In this population 4 of 10 isolated clones showed a C_16_ to T_16_ mutation (Figure 2B, *gapA* edited). Finally, in the *cysP* target (Figure 2C) (5’-**<u>C</u>**_20_AG**<u>C</u>**_17_AATGAGGAGGAATTTT/CGG-3’), C_17_ and C_20_ were potential deamination sites. After induction and population analysis, the conversion levels were 76% and 77% for these two cytosines, respectively. All 10 out of 10 isolated clones showed mutations at the two positions (Figure 2C, *cysP* edited). Thus, the editing window of the mycoplasma CBE system ranged from positions 16 to 20, upstream of the PAM sequence (Figure 2D).

**Figure 2.**
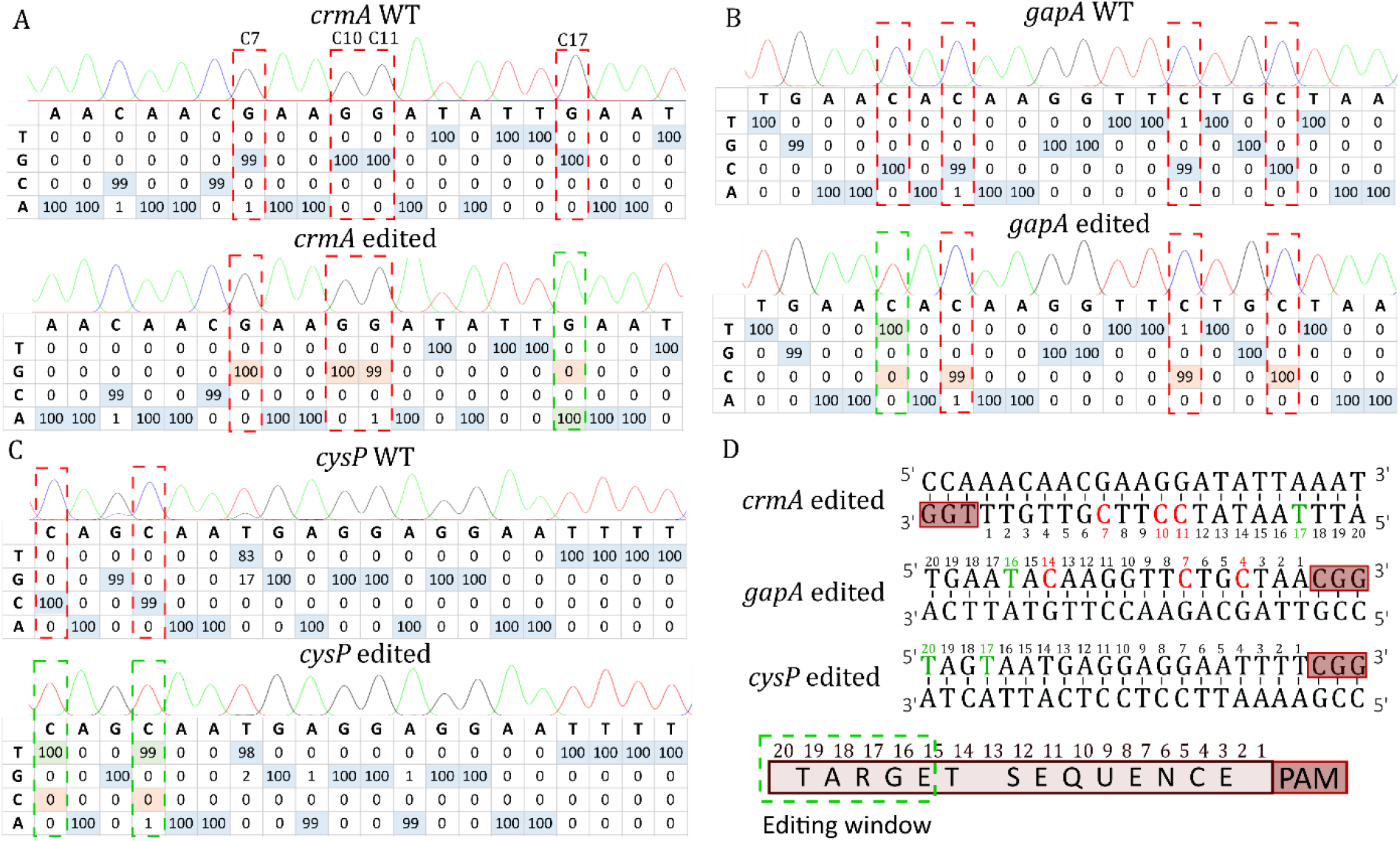
Targeting of three virulence factors of *Mgal* to explore the editing window of the CBE system in *Mgal*. The 20-nucleotide target sites were sequenced and are represented here for the *crmA* (**A**), *gapA* (**B**), and c*ysP* (**C**) genes before and after the base-editing experiments. Cytosines susceptible to deamination (or guanine in the reverse strand) are framed in red and those that were deaminated in green. The percentage of each base is shown in the tables, as determined using the Sanger sequencing.ab file and EditR software. Edited bases are highlighted in green. For the *crmA* target, the complementary strand was sequenced. **D-** The 20-nucleotides target sites are shown for the three targets. The position of each base in the target is indicated below each nucleotide and corresponds to the nucleotide position in the target upstream of the PAM sequence. Red letters represent un-deaminated cytosines and green letters converted thymines. A scheme based on the three experiments highlighting the editing window is shown at the bottom. As shown in the scheme, cytosines located at positions 16 to 20 can be CBE-targeted.

### Development of the CBE system for *Mbov*

We initially evaluated pTI4.0_spdCas9_pmcDA1, successfully used in *Mgal,* in *Mbov*. However, we did not obtain *Mbov* transformants using this CBE-carrying plasmid. This may be linked to use of the *pac* (puromycin) resistance marker, which has not been described in the literature for *Mbov*. We therefore inserted the inducible CBE system in another transposon-based plasmid carrying a gentamycin resistance marker to generate pMT85_SpdCas9_pmcDA1 (Figure S2). The resulting construct was evaluated in *Mbov* using the *mnuA* gene (MBOVPG45_0215) as a target (Figure 3). MnuA is a major membrane nuclease that has been suggested to play a key role in *Mbov* virulence by degrading the neutrophil extracellular traps (NETs) produced by the host in response to the pathogen (39). In the target spacer (5’-AA**<u>C</u>**_18_**<u>C</u>**_17_AAAAATATGACTTAGT/AGG-3’), C_17_ and C_18_ stand as two potential deamination sites. Conversion of C_17_ would change a Glutamine CAA codon into a TAA stop codon and disrupt the *mnuA* gene. *Mbov* cells were transformed with pMT85_SpdCas9_pmcDA1 targeting the *mnuA* gene and the transformants grown in gentamycin selective medium (liquid) for three passages. Expression of the CBE system was induced overnight with 0.5 μg.mL^−1^ aTC and *Mbov* transformants were plated on selective medium to isolate colonies. Target sites were analyzed in the global population before transformation, after each passage following transformation (P1, P2, and P3) and after induction (for the cell suspension and isolated colonies) (Figure 3). As previously found in *Mgal*, there was considerable leakage of the inducible promoter in the *Mbov* population immediately after transformation, with ~50% of the cytosines converted to thymines at positions C_17_ and C_18_ at P1, P2, and P3. Overnight induction with aTC increased the conversion level to 90%. Screening of three independent colonies showed both cytosines (C_17_ and C_18_) in the *Mbov_mnuA* mutants to have been converted to thymine. We then carried out a nuclease phenotypic assay to demonstrate that the mutations introduced into the *mnuA* gene led to its inactivation (40). Indeed, the tested *Mbov_mnuA* mutant was unable to hydrolyze DNA, whereas the WT strain degraded it all (Figure S6). These results demonstrate the portability of this CBE-based method to produce targeted mutations in *Mbov*.

**Figure 3.**
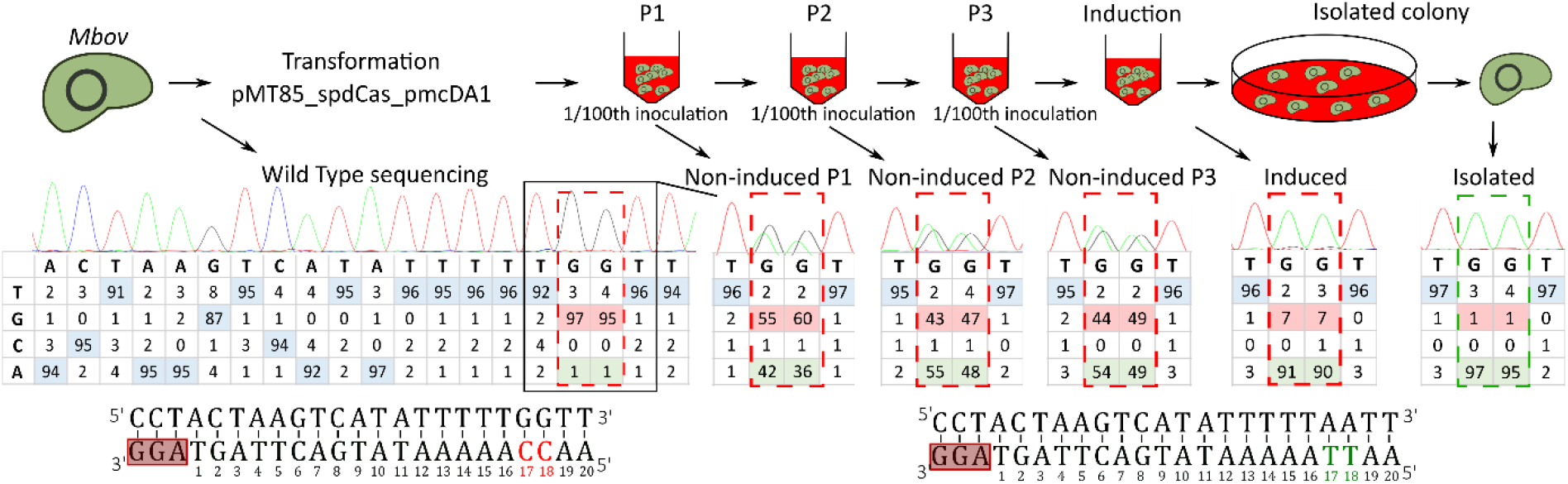
Disruption of the MnuA nuclease-encoding gene by base editing in *Mbov*. After transformation of *Mbov* with the plasmid pMT85_SpdCas9_pmcDA1, cells were propagated in liquid media supplemented with gentamycin. After three passages in liquid medium (P1 to P3), the inducer was added to the cell suspension. Aliquots were collected at each passage and after induction to monitor the target site sequence by Sanger sequencing and EditR analysis. Sequencing chromatograms of the complementary strand are presented, as well as the percentage of the base for each position of the target sequence. Cytosines susceptible to deamination are framed in red and the edited bases in green.

### Development of the CBE system for editing *Mycoplasma mycoides* subsp. *mycoides* genome

We next introduced the sgRNA and CBE expression cassettes into the replicative *oriC* plasmid pMYCO1 to demonstrate the flexibility of the mycoplasma CBE system. This plasmid is routinely used in several species of the mycoides cluster (20), including *Mmm*. The resulting pMYCO1_SpdCas9_pmcDA1 plasmid (Figure S2) was used to target the non-essential gene *glpO* (5’-**<u>C</u>**_20_AA**<u>C</u>**_17_AA**<u>C</u>**_14_AATA**<u>C</u>**_9_GATAA**<u>C</u>**_3_AT/TGG-3’), which encodes the metabolic enzyme L-α-glycerophosphate oxidase putatively involved in mycoplasma virulence (41). Indeed, GlpO catalyzes the oxidation of glycerol-3-phosphate, leading to the release of hydrogen peroxide (H_2_O_2_), a product known to contribute to cytopathic effects in host tissue. C_3_, C_9_, C_14_, C_17_, and C_20_ were potential deamination sites, with conversion of the last three positions leading to three TAA stop codons. After transformation of *Mmm*, we tested various inducer concentrations and induction times and monitored conversion of the cytosines in the cell population (Figure S7). Before the induction of CBE expression, conversion of cytosines was observed as in *Mgal* and *Mbov*, reaching up to 30% at position C_17_. The best results were obtained after overnight induction with 0.5 μg.mL^−1^ aTC. We observed conversion for each cytosine within the target region, as well as for the one located 22 nt upstream of the PAM sequence, suggesting an extended editing window between positions C_14_ and C_22_ in *Mmm*. We observed low conversion frequencies (20-30%) at the editing window extremities, whereas higher efficiencies were observed between C_17_ and C_20_ (60-70%). After subcloning on agar plates, the mutants showed diverse editing profiles at the four cytosine positions (mixed population and different fully deaminated cytosine combinations). Nevertheless, 2 of 10 isolated clones showed the expected four mutations (C_14_, C_17_, C_20_, and C_22_). A phenotypic assay to evaluate H_2_O_2_ production confirmed inactivation of the *glpO* gene in these clones (Figure S7E). Finally, after three passages in liquid medium without selective pressure and one passage in solid medium for isolation, we recovered plasmid-free *Mmm* mutants edited at *glpO*. These are the first reported site-specific mutants generated in *Mmm*.

### Inducing multiple mutations in *Mmm* with a single sgRNA

We tested the limits of the mycoplasma CBE system by targeting transposase-encoding genes associated with two families of insertion sequence, IS*1634* and IS*3*, present 58 and 26 times in the *Mmm* T1/44 genome, respectively (Figures 4 and S8). As transposase gene sequences are highly conserved within each family, we attempted to target a maximum number of copies using a single sgRNA per family. For the IS*1634* transposase, a 20-nucleotides spacer was designed to perform the inactivation of 55 of 58 copies (5’-AGA**<u>C</u>**_17_**<u>C</u>**_16_AGATTGTTATAGGTA/TGG-3’). Deamination of C_16_ on the reverse strand would change a Glutamine CAG codon into a TAG stop codon at position 204 of the protein (Figure 4A). For the IS*3* transposase, two 20-nucleotide spacers were designed for the sgRNAs, the first targeting the start codon of all 26 copies of the transposase (sgRNA1, 5’-**<u>C</u>**_20_ATATAAAAACCCCATTTCC/TGG-3’) and the second allowing the introduction of a stop codon in 22 of the 26 copies (sgRNA2, 5’-A**<u>C</u>**_19_AAGTGGAATACTATAAGT/TGG-3’) (Figure S8). After transformation and induction, all target sites were assessed by PCR and Sanger sequencing (Figures 4B and S7B). We observed low base-editing efficiency for the *IS1634* target after the first induction, with only 5% conversion for C_16_ and 4% for C_17_. Two additional induction steps were carried out, resulting in an increase in base conversion of C_16_ and C_17_, with 37% and 40% after the second induction and 72% and 75% after the third induction, respectively. After plating and PCR/sequencing screening of 20 isolated clones, two showed sequencing profiles suggesting that all IS*1634* sites had been mutated (Figure 4C). Finally, full genome sequencing showed that 52 and 54 transposases from among 55 were inactivated in these two clones. For IS*3* inactivation, 66% and 77% conversion were observed in population analyses after three inductions using sgRNAs with spacers one and two, respectively (Figure S8B). After plating and subsequent PCR screening of 10 clones, one fully deaminated clone was rescued for each of the two sgRNAs (Figure S8C). Full genome sequencing confirmed mutations at 22/26 and 22/22 target sites with sgRNA1 and sgRNA2, respectively (Table S3). These results demonstrate the ability of the mycoplasma CBS system to edit numerous target sites in a single step.

**Figure 4.**
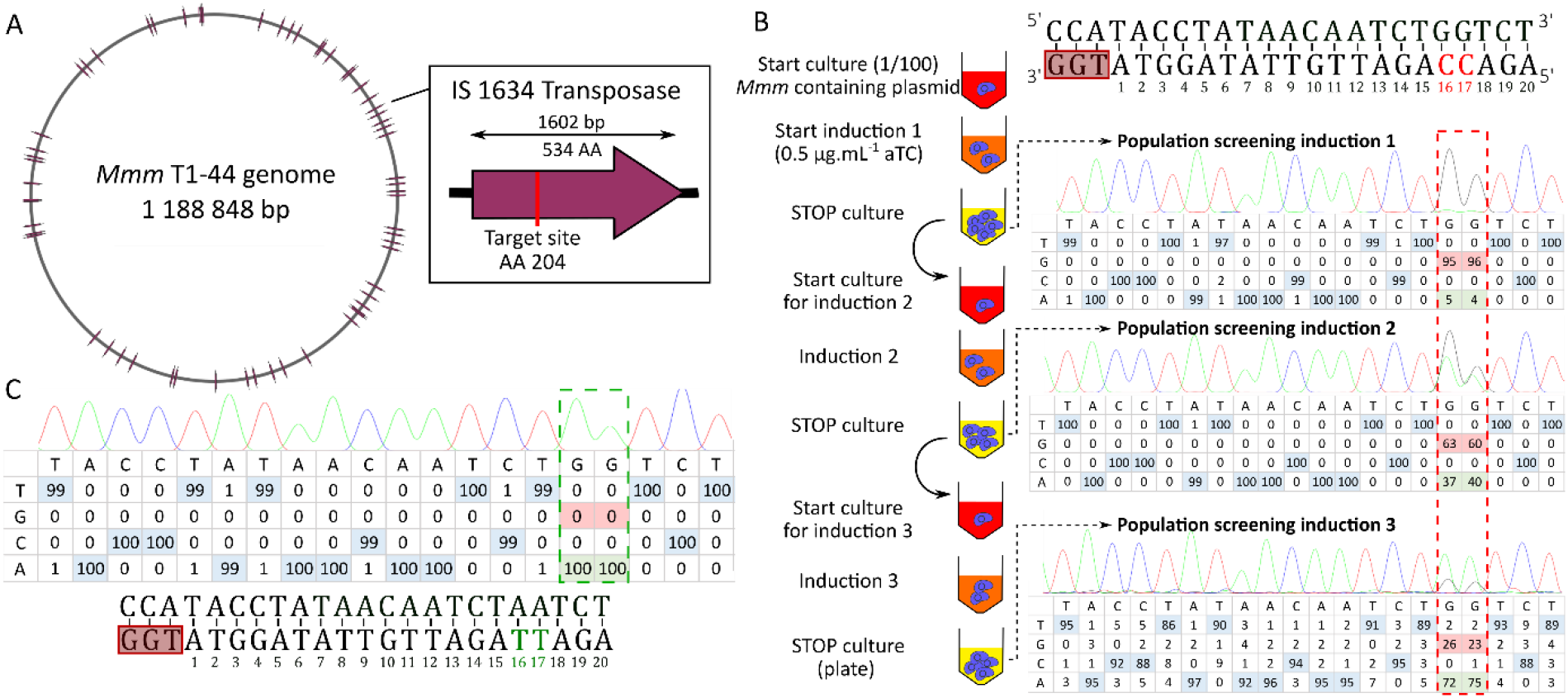
Multitargeting of IS *1634* copies in the *Mmm* genome. **A-** Schematic representation of the *Mmm* T1/44 genome, with 58 complete or truncated copies of the IS*1634* transposases represented as solid lines. In the right panel, the targeted site within the IS*1634* transposase gene is represented as a red bar. **B-** Schematic representation of the three induction steps of the mycoplasma CBE system for targeting 55 IS*1634* sites using a single gRNA. The nucleotide sequence of the target is represented with the PAM sequence framed by a red rectangle; cytosines susceptible to deamination are shown in red. Fresh cultures of *Mmm* cells harboring the pMYCO1_SpdCas_pmcDA1 plasmid are represented as red-colored tubes. Cultures in exponential and stationary growth phases are indicated by orange- and yellow-colored tubes, respectively. Three consecutive culture steps were performed (including the starting culture) by a 1/100 dilution in fresh medium. At each step, induction was performed by adding aTC and incubating the cells until stationary phase was reached. Deamination of target cytosines was evaluated by PCR and Sanger sequencing. The percentage of each base at each position was estimated from chromatograms using EditR software. The two positions of interest are highlighted in a dashed red rectangle. **C-** Sanger sequencing results and base distribution in a selected isolated clone. Fully mutated positions are framed in green in the table and shown in green in the target sequence.

### Evaluation of off-target mutations by whole genome sequencing

We investigated potential undesired mutations and evaluated the off-target activity of the CBE system in mycoplasmas by carrying out whole genome sequencing on seven edited (mutant) clones using both short-read (Illumina) and long-read (Oxford Nanopore Technologies) sequencing platforms: *Mgal_ksgA*, *Mbov_0215*, *Mmm_glpO*, *Mmm_IS1634_cl3.1.6*, *Mmm_IS1634_cl3.1.11*, *Mmm_IS3_cl4.1.2*, and *Mmm_IS3_cl5.1.18* (Table S3). We detected 5 to 36 undesired mutations in the sequenced genomes. Mutations were further analyzed and classified into three categories (Figure S9A, Table S3): 1) sgRNA-guided off-target mutations possibly generated by the CBE system, if the targeted sequence was similar to the sgRNA spacer sequence and adjacent to a NGG PAM sequence, 2) potentially unguided spurious deamination by pmcDA1 (28, 42) for all C-T or G-A mutations in genome regions with no similarity to the sgRNA spacer sequence, and 3) other mutations that could not have been induced by the CBE system and that potentially occurred during passaging. Based on these criteria, 0 to 5 sgRNA-guided off-target mutations were predicted, representing 0 to 17% of the total mutations, with the exception of *Mmm_glpO*, for which 2 of 5 mutations were classified in this category. Two examples of off-target interactions of the *ksgA* sgRNA targeting DNA in *Mgal* (mutation at position 1,313 in the genome) and the *glpO* sgRNA targeting DNA in *Mmm* (mutation at position 220,532 in the genome) are presented (Figures S9B, S9C, Table S3). Spurious deamination represented 20 to 84% of mutations, whereas other variations represented 11 to 50%. Interestingly, the number and percentage of spurious deamination that occurred when using the mycoplasma CBE system in *Mmm* were much higher when a three-induction protocol was used to maximize the efficiency of IS*1634* and IS*3* targeting (11-24 mutations) than the single induction protocol used to target *glpO* (1 mutation). Thus, sgRNA-guided off-target mutations were relatively rare, whereas spurious deaminations accumulated with extended CBE activity.

## Discussion

The functional genomics of mollicutes has long been hampered by the absence of efficient genetic tools to generate targeted mutations. Here, we developed a new genetic tool based on CRISPR for direct mutagenesis in mollicutes and evaluated its efficiency in three economically relevant animal pathogens.

CBE systems have proven to be highly efficient in diverse eukaryotic, including plant and human cells (43, 44) but also in prokaryotic organisms including *Klebsiella* (45), *Pseudomonas* (46), *Clostridium* (47), *Streptomyces* (29), and *Agrobacterium* (48). Here, we designed and constructed two mycoplasma CBE systems, but our preliminary results suggested that only the pmcDA1-based CBE was active in *Mgal* (Figures 1, S3). Both cytosine deaminases have been successfully used in bacteria (29, 30) and further studies would be needed to understand why the rAPOBEC1-based CBE was not efficient in *Mgal*. By contrast, the pmcDA1-based CBE was highly efficient in generating point mutations in *Mgal*, *Mbov*, and *Mmm* (Figures 1, 3, S7). Although belonging to different phylogenetic groups of mollicutes, the same promoters were used to drive the expression of the sgRNA (PS) and CBE system (P*xyl/tetO2*), suggesting that both expression cassettes could be used without modification in various species of mycoplasmas and possibly other mollicutes. Three different plasmid backbones were used here, including two Tn*4001*-derived transposons and one replicative *oriC* plasmid (Figure S2). Transposons and *oriC* plasmids are currently the most widely used genetic tools available in mollicutes. Transposon mutagenesis with Tn*4001* derivatives has been used in 15 species and *ori*C plasmids are available for 14. The mycoplasma CBE system could thus be easily evaluated in many species, either directly or after cloning the sgRNA and CBE expression cassettes into a compatible vector. During this study, we also found that induction could be performed immediately following plasmid transformation (*i.e.,* immediately after the 2-h cell recovery in Hayflick medium) instead of being carried out after colony recovery and regrowth. This improvement allowed us to reduce the time for experiments by 14 days and obtain the expected mutants in less than three weeks.

The cell toxicity of the sgRNA/SpdCas9-pmcDA1-UGI complex was low as expected and no particular difficulty was encountered when expressing the CBE system in the three mycoplasmas tested here. Taking advantage of such tool, we succeeded in targeting up to 54 targets in the *Mmm* chromosome with a single sgRNA (Figure 4). Multiple targeting of repeated sequences has been previously reported in eukaryotic and prokaryotic systems (30, 31, 49), but this is the first time that so many target sites have been modified in bacteria. This application of CBE opens new possibilities for targeting multigene families with a limited number of sgRNA molecules. In mycoplasmas, such families are often predicted to encode surface proteins suspected to be involved in host-pathogen interactions but current mutagenesis methods, including in-yeast genome engineering, are unable to disrupt all genes of a family in a reasonable amount of time.

CBE used in other organisms have been shown to have some off-target effects. Here, we selected seven mutants for whole genome sequencing to evaluate potential off-target mutations induced by the mycoplasma CBE system (Figure S9, Table 3). Analysis of the detected mutations showed that only a few could have resulted from sgRNA-guided off-target deamination events. Spurious deamination of pmcDA1 which preferentially deaminates TC motifs (28), appeared to increase with extended induction periods, suggesting that the control of CBE expression could be crucial to reducing the frequency of undesired mutations. Such spurious deamination induced by CBE systems has already been reported in other bacteria, including *Corynobacterium glutamicum* and *Bacillus subtilis*, in which 9 and 19 SNVs, respectively, could be attributed to deaminase activity (50, 51). In the CBE system developed here, expression of the SpdCas9-pmcDA1-UGI hybrid protein is driven by the promoter P*xyl/tetO2*, which can be induced by aTC. Although we observed a clear increase in the frequency of deaminated bases in the three studied mycoplasmas after induction, the conversion process was already observed before induction (Figures 1, 3 and S7). This indicates a certain level of promoter leakage in the three species, in accordance with another report on *M. pneumoniae* (52). Several strategies can be envisaged to limit the background of undesired mutations, including limiting the induction time and using an improved inducible promoter, such as that recently developed for *M. pneumoniae* (52). In addition, the use of high-fidelity Cas9 variants or CBE variants (35) may also improve the specificity of the genetic tool. Finally, elimination of the CBE system after mutagenesis is also crucial for avoiding the accumulation of undesired mutations over time. This can be achieved for the CBE systems based on the pMT85_2Res and pMYCO1 backbones. Indeed, all genetic elements flanked by the *res* sequences in the pMT85_2 Res can be removed from the chromosome using dedicated resolvase activity (53) and the *oriC* plasmids such as the pMYSO1 are generally lost after a few passages in non-selective medium (54). Alternatively, CBE constructs can be enhanced with the CRE-Lox system, which is functional in some mycoplasma species (21, 52, 55).

In conclusion, the mycoplasma CBE system is an easy-to-use genetic tool that we have shown to be highly efficient for the targeting of individual or multiple genes in three mycoplasma species for which no targeted mutagenesis methods were previously available. This tool is independent from cell repair mechanisms and makes it possible to obtain an isolated mutant of *Mgal, Mbov*, or *Mmm* in less than a month. During this study, we already generated individual mutants of potential virulence genes for each species, demonstrating the efficacy and versatility of this new genetic tool.

## Materials and Methods

### Oligonucleotides and plasmids

All oligonucleotides used in this study were supplied by Eurogentec and are described in Table S1. All plasmids constructed and used in this study are listed in Table S2. Detailed protocols for plasmid construction and other methods are provided as **SI Materials and Methods**.

### Bacterial strains and culture

*Mgal* strain S6 (Tax ID: 1006581) was cultivated at 37°C in Hayflick modified medium (56) in a 5% CO_2_ atmosphere and 10 μg.mL^−1^ puromycin and 400 μg.mL^−1^ kasugamycin were used for selection. *Mbov* PG45 (Tax ID: 289397) was cultivated in SP4 medium (56) and 100 μg.mL^−1^ gentamycin was used for selection. *Mmm* T1/44 (Tax ID: 2103) was cultivated in SP5 medium (18) and 8 μg.mL^−1^ puromycin was used for selection (57). Phenol red was used as a pH indicator in the mycoplasma media. *Escherichia coli* NEB-5α (NEB, C2987H) was used for plasmid propagation and was cultivated in LB Broth (ThermoFisher: 12795027), with the addition of 100 μg.mL^−1^ ampicillin or 50 μg.mL^−1^ kanamycin for selection.

### Transformation of *Mycoplasma* species

Mycoplasma transformation was performed using a polyethylene glycol mediated protocol (58, 59). Late-log phase mycoplasma cultures were transformed with plasmid DNA (20 μg). After transformation, cells were resuspended in 1 mL of the appropriate medium, incubated 2 h at 37°C, and plated onto selective solid medium. After incubation at 37°C for 3 to 10 days, single colonies containing the deaminase constructs were obtained. Expression of the CBE system was induced in early logarithmic growth-phase cultures overnight and the cultures plated on selective medium to isolate colonies.

## Supporting information

**Supplementary Figures**. Figure S1, Schematic representation of the CBE system; Figure S2, Design of the three plasmids for genome editing in mycoplasma; Figure S3, Targeting of the *ksgA* gene in *Mgal* using the rAPOBEC1deaminase-encoding plasmid; Figure S4, Optimization of CBE induction conditions for base editing in *Mgal*; Figure S5, Kasugamycin resistance phenotypic assay on a *Mgal_ksgA* mutant; Figure S6, *Mbov*_*mnuA* DNase phenotypic assay; Figure S7, Targeting the *glpO* gene in *Mmm* using the mycoplasma CBE system; Figure S8, Multitargeting of IS*3* copies in the *Mmm* genome; Figure S9, Analysis of undesired mutations from whole genome sequences.

**Supplementary tables.** Table S1. List of oligonucleotides used in this work; Table S2, list of plasmids used in this work; Table S3. Mutations detected in edited genomes after WGS.

**Supplementary text. SI-1. Materials and Methods. SI-2. Genes and plasmids sequences.**

## Acknowledgements and funding sources

This work was partially funded by ANR, as part of the project RAMbo-V (ANR-21-CE35-0008). Genome sequencing was performed by the Genome Transcriptome Facility of Bordeaux (https://pgtb.cgfb.u-bordeaux.fr, Grants from Investissements d’avenir, Convention attributive d’aide EquipEx Xyloforest ANR-10-EQPX-16-01).

## Author contribution

Conceptualization, TI, PS-P; Formal Analysis, TI, FR; Funding acquisition, YA, CL, AB, PS-P; Investigation, TI, FR, GG; Methodology, TI, PS-P; Supervision, CL, PS-P; Validation, YA, CL, AB, PS-P; Visualization, TI, PS-P; Writing – original draft, TI, AB, PS-P; Writing – review & editing, TI, FR, YA, CL, AB, PS-P

## References

1. M. May, M. F. Balish, A. Blanchard, The Order Mycoplasmatales (2014) https:/doi.org/https://doi.org/10.1007/978-3-642-30120-9_289.

2. P. Sirand-Pugnet, C. Citti, A. Barré, A. Blanchard, Evolution of mollicutes: down a bumpy road with twists and turns. Res. Microbiol. 158, 754–766 (2007).

3. S. Razin, D. Yogev, Y. Naot, Molecular biology and pathogenicity of Mycoplasmas. Microbiol. Mol. Biol. Rev. 165, 199–221 (1998).

4. C. Citti, E. Dordet-Frisoni, L. X. Nouvel, C. H. Kuo, E. Baranowski, Horizontal gene transfers in mycoplasmas (Mollicutes). Curr. Issues Mol. Biol. 29, 3–22 (2018).

5. H. Grosjean, et al., Predicting the Minimal Translation Apparatus: Lessons from the Reductive Evolution of Mollicutes. PLoS Genet. 10 (2014).

6. C. Citti, A. Blanchard, Mycoplasmas and their host: Emerging and re-emerging minimal pathogens. Trends Microbiol. 21, 196–203 (2013).

7. R. Rosengarten, et al., The changing image of mycoplasmas: from innocent bystanders to emerging and reemerging pathogens in human and animal diseases. Contrib. Microbiol. 8, 166–185 (2001).

8. S. L. Hennigan, et al., Detection and differentiation of avian mycoplasmas by surface-enhanced raman spectroscopy based on a silver nanorod array. Appl. Environ. Microbiol. 78, 1930–1935 (2012).

9. E. D. Peebles, S. L. Branton, *Mycoplasma gallisepticum* in the commercial egg-laying hen: A historical perspective considering the effects of pathogen strain, age of the bird at inoculation, and diet on performance and physiology. J. Appl. Poult. Res. 21, 897–914 (2012).

10. CABI, Avian mycoplasmosis (2019).

11. K. Dudek, E. Szacawa, *Mycoplasma bovis* infections: Occurrence, pathogenesis, diagnosis and control, including prevention and therapy. Pathogens 9, 1–3 (2020).

12. A. Jordan, et al., *Mycoplasma bovis* outbreak in New Zealand cattle: An assessment of transmission trends using surveillance data. Transbound. Emerg. Dis. 68, 3381–3395 (2021).

13. G. Di Teodoro, et al., Contagious Bovine Pleuropneumonia: A Comprehensive Overview. Vet. Pathol. 57, 476–489 (2020).

14. A. V. Gautier-Bouchardon, Antimicrobial Resistance in Mycoplasma spp. Microbiol. Spectr. 6 (2018).

15. M. Ishfaq, et al., Current status of vaccine research, development, and challenges of vaccines for *Mycoplasma gallisepticum*. Poult. Sci. 99, 4195–4202 (2020).

16. E. Ruiz, et al., CReasPy-Cloning: A Method for Simultaneous Cloning and Engineering of Megabase-Sized Genomes in Yeast Using the CRISPR-Cas9 System. ACS Synth. Biol. (2019) https:/doi.org/10.1021/acssynbio.9b00224.

17. C. Lartigue, et al., Creating bacterial strains from genomes that have been cloned and engineered in yeast. Science (80-.). 325, 1693–1696 (2009).

18. F. Labroussaa, et al., Impact of donor-recipient phylogenetic distance on bacterial genome transplantation. Nucleic Acids Res. 44, 8501–8511 (2016).

19. R. Chopra-Dewasthaly, M. Zimmermann, R. Rosengarten, C. Citti, First steps towards the genetic manipulation of *Mycoplasma agalactiae* and *Mycoplasma bovis* using the transposon Tn4001mod. Int. J. Med. Microbiol. 294, 447–453 (2005).

20. C. Janis, et al., Versatile Use of oriC Plasmids for Functional Genomics of *Mycoplasma capricolum* subsp. *capricolum*†. Society 71, 2888–2893 (2005).

21. T. Ipoutcha, G. Gourgues, C. Lartigue, A. Blanchard, P. Sirand-Pugnet, Genome Engineering in Mycoplasma gallisepticum Using Exogenous Recombination Systems. ACS Synth. Biol., acssynbio.1c00541 (2022).

22. M. Jinek, et al., A programmable dual-RNA-guided DNA endonuclease in adaptive bacterial immunity. Science (80-.). 337, 816–821 (2012).

23. C. Piñero-Lambea, et al., *Mycoplasma pneumoniae* Genome Editing Based on Oligo Recombineering and Cas9-Mediated Counterselection. ACS Synth. Biol. 9, 1693–1704 (2020).

24. A. C. Komor, Y. B. Kim, M. S. Packer, J. A. Zuris, D. R. Liu, Programmable editing of a target base in genomic DNA without double-stranded DNA cleavage. Nature 533, 420–424 (2016).

25. K. Nishida, et al., Targeted nucleotide editing using hybrid prokaryotic and vertebrate adaptive immune systems. Science (80-.). 353 (2016).

26. N. M. Gaudelli, et al., Programmable base editing of T to G C in genomic DNA without DNA cleavage. Nature 551, 464–471 (2017).

27. A. C. Komor, et al., Improved base excision repair inhibition and bacteriophage Mu Gam protein yields C:G-to-T:A base editors with higher efficiency and product purity. Sci. Adv. 3, 1–10 (2017).

28. A. V. Anzalone, L. W. Koblan, D. R. Liu, Genome editing with CRISPR–Cas nucleases, base editors, transposases and prime editors. Nat. Biotechnol. 38, 824–844 (2020).

29. Y. Tong, et al., Highly efficient DSB-free base editing for streptomycetes with CRISPR-BEST. Proc. Natl. Acad. Sci. U. S. A. 116, 20366–20375 (2019).

30. S. Banno, K. Nishida, T. Arazoe, H. Mitsunobu, A. Kondo, Deaminase-mediated multiplex genome editing in *Escherichia coli*. Nat. Microbiol. 3, 423–429 (2018).

31. S. J. Bae, B. G. Park, B. G. Kim, J. S. Hahn, Multiplex Gene Disruption by Targeted Base Editing of Yarrowia lipolytica Genome Using Cytidine Deaminase Combined with the CRISPR/Cas9 System. Biotechnol. J. 15, 1–46 (2019).

32. M. Breton, et al., First report of a tetracycline-inducible gene expression system for mollicutes. Microbiology 156, 198–205 (2010).

33. G. G. Mahairas, F. C. Minion, Random Insertion of the Gentamicin Resistance Transposon Tn4001 in *Mycoplasma pulmonis*. Plasmid (1989).

34. A. M. Mariscal, et al., Tuning Gene Activity by Inducible and Targeted Regulation of Gene Expression in Minimal Bacterial Cells. ACS Synth. Biol. 7, 1538–1552 (2018).

35. T. P. Huang, G. A. Newby, D. R. Liu, Precision genome editing using cytosine and adenine base editors in mammalian cells. Nat. Protoc. 16, 1089–1128 (2021).

36. L. Papazisi, et al., GapA and CrmA coexpression is essential for *Mycoplasma gallisepticum* cytadherence and virulence. Infect. Immun. 70, 6839–6845 (2002).

37. S. P. Mugunthan, M. C. Harish, Multi-epitope-Based Vaccine Designed by Targeting Cytoadherence Proteins of *Mycoplasma gallisepticum*. ACS Omega 6, 13742–13755 (2021).

38. I. Cizelj, et al., *Mycoplasma gallisepticum* and *Mycoplasma synoviae* express a cysteine protease CysP, which can cleave chicken IgG into Fab and Fc. Microbiology 157, 362–372 (2011).

39. F. Mitiku, et al., The major membrane nuclease MnuA degrades neutrophil extracellular traps induced by *Mycoplasma bovis*. Vet. Microbiol. 218, 13–19 (2018).

40. S. Sharma, K. A. Tivendale, P. F. Markham, G. F. Browning, Disruption of the membrane nuclease gene (MBOVPG45_0215) of *Mycoplasma bovis* greatly reduces cellular nuclease activity. J. Bacteriol. 197, 1549–1558 (2015).

41. P. Pilo, et al., A metabolic enzyme as a primary virulence factor of Mycoplasma *Mycoplasma mycoides* subspecies *mycoides* small colony. J. Bacteriol. (2005) https:/doi.org/10.1128/JB.187.19.6824-6831.2005.

42. Y. Yu, et al., Cytosine base editors with minimized unguided DNA and RNA off-target events and high on-target activity. Nat. Commun. 11, 1–10 (2020).

43. Y. Wu, et al., Increasing cytosine base editing scope and efficiency with engineered Cas9-PMCDA1 fusions and the modified sgRNA in rice. Front. Genet. 10, 1–10 (2019).

44. H. S. Kim, Y. K. Jeong, J. K. Hur, J. S. Kim, S. Bae, Adenine base editors catalyze cytosine conversions in human cells. Nat. Biotechnol. 37, 1145–1148 (2019).

45. Y. Wang, et al., CRISPR-Cas9 and CRISPR-Assisted Cytidine Deaminase Enable Precise and Efficient Genome Editing in Klebsiella pneumoniae. Appl. Environ. Microbiol. 84, 1–15 (2018).

46. J. Sun, L. B. Lu, T. X. Liang, L. R. Yang, J. P. Wu, CRISPR-Assisted Multiplex Base Editing System in *Pseudomonas putida* KT2440. Front. Bioeng. Biotechnol. 8, 1–13 (2020).

47. Q. Li, et al., CRISPR–Cas9 D10A nickase-assisted base editing in the solvent producer Clostridium beijerinckii. Biotechnol. Bioeng. 116, 1475–1483 (2019).

48. S. D. Rodrigues, et al., Efficient CRISPR-mediated base editing in agrobacterium spp. Proc. Natl. Acad. Sci. U. S. A. 118, 1–8 (2021).

49. Z. Zhong, et al., Base editing in Streptomyces with Cas9-deaminase fusions. BioRxiv, 630137 (2019).

50. Y. Wang, et al., MACBETH: Multiplex automated *Corynebacterium glutamicum* base editing method. Metab. Eng. 47, 200–210 (2018).

51. S. Yu, et al., CRISPR-dCas9 mediated cytosine deaminase base editing in *Bacillus subtilis*. ACS Synth. Biol. (2020) https:/doi.org/10.1021/acssynbio.0c00151.

52. A. M. Mariscal, L. González-González, E. Querol, J. Piñol, All-in-one construct for genome engineering using Cre-lox technology. DNA Res. 23, 263–270 (2016).

53. C. Janis, et al., Unmarked insertional mutagenesis in the bovine pathogen *Mycoplasma mycoides* subsp. *mycoides* SC. Microbiology 154, 2427–2436 (2008).

54. C. Lartigue, A. Blanchard, J. Renaudin, F. Thiaucourt, P. Sirand-Pugnet, Host specificity of mollicutes oriC plasmids: Functional analysis of replication origin. Nucleic Acids Res. 31, 6610–6618 (2003).

55. L. Garcia-Morales, et al., A RAGE Based Strategy for the Genome Engineering of the Human Respiratory Pathogen *Mycoplasma pneumoniae*. ACS Synth. Biol. 9, 2737–2748 (2020).

56. E. A. Freundt, “CULTURE MEDIA FOR CLASSIC MYCOPLASMAS” in Methods in Mycoplasmology, (1983) https:/doi.org/10.1016/b978-0-12-583801-6.50029-9.

57. M. A. Algire, et al., New selectable marker for manipulating the simple genomes of Mycoplasma species. Antimicrob. Agents Chemother. 53, 4429–4432 (2009).

58. K. W. King, K. Dybvig, Transformation of *Mycoplasma capricolum* and examination of DNA restriction modification *Mycoplasma capricolum* and *Mycoplasma mycoides subsp. mycoides*. Plasmid 31, 308–311 (1994).

59. X. Zhu, et al., Mbov_0503 encodes a novel cytoadhesin that facilitates *Mycoplasma bovis* interaction with tight junctions. Microorganisms 8 (2020).

