## Supplementary Information for "Unblocking genome editing of major animal mycoplasmas using CRISPR/Cas9 base editor systems"

#### Supplementary Figures

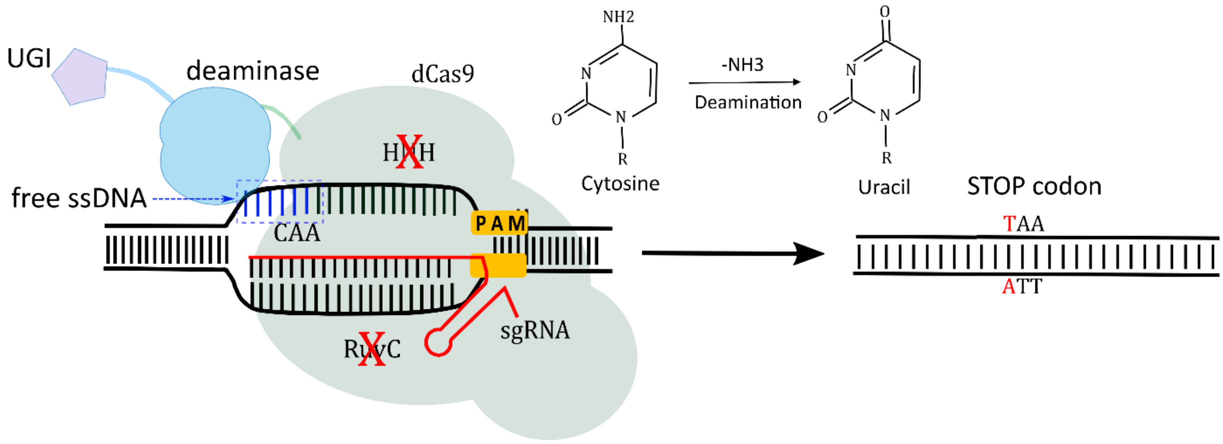


**Figure S1. Schematic representation of the CBE system.** An R-loop mediated by the dCas9 (Green) – sgRNA (red) complex is represented. UGI (uracil glycosylase inhibitor, purple) and the deaminase protein (light blue) are fused to dCas9 by an amino-acid linker. The editing window is represented as free ssDNA (dark blue) and corresponds to the only nucleotide positions where deaminase can act. The cytosine deamination reaction is represented.

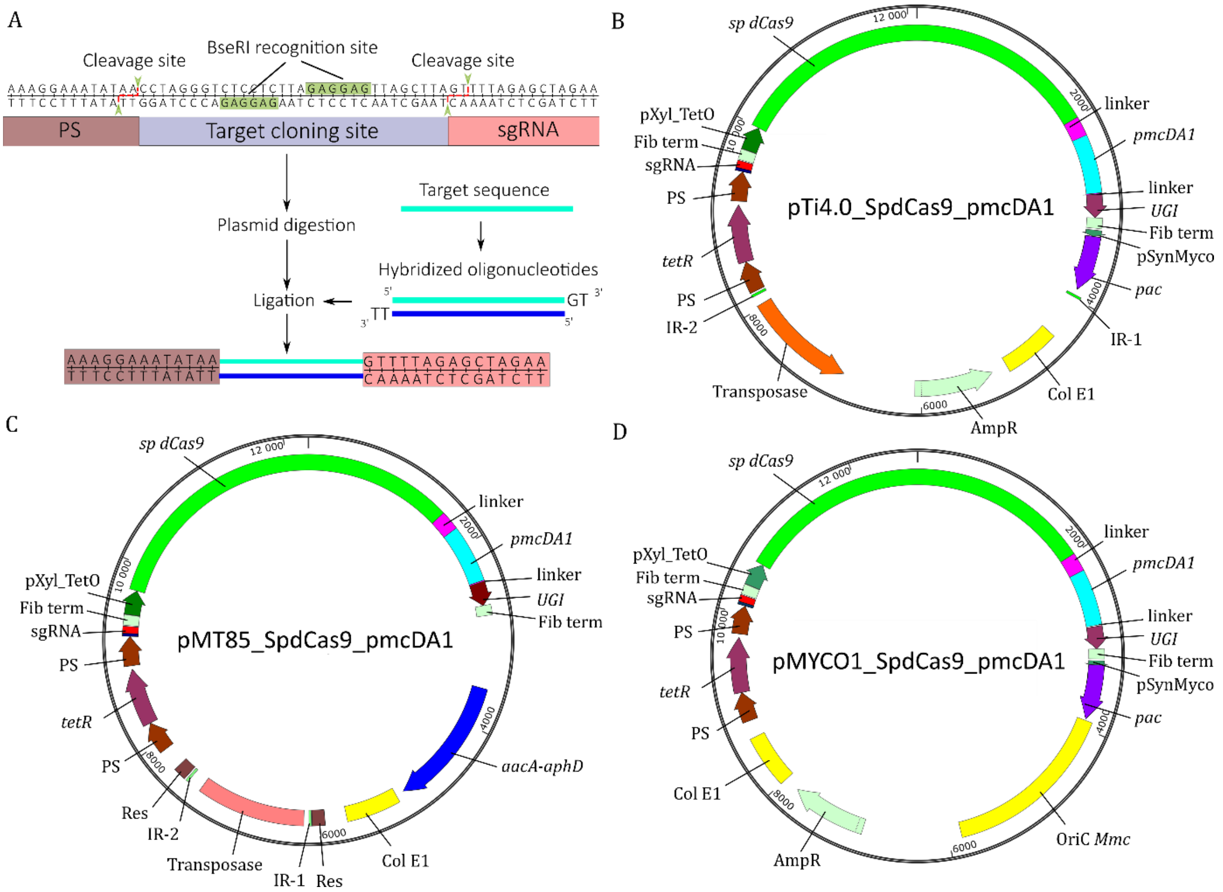


**Figure S2. Design of the three plasmids for genome editing in mycoplasma. A-** Scheme of the method used to add a 20-nucleotide spacer defining the target site of the sgRNA using double BseRI sites. **B-** Representation of the three plasmid backbones used for base editing experiments in mycoplasma. The antibiotic resistance markers *pac* (puromycin) and *aacA-aphD* (gentamicin) are represented in dark purple and blue, respectively. The pTi4.0_SpCas9_pmcDA1 plasmid (for *Mgal* experiments) contains inverted repeats (IR-1 and IR-2, green) flanking the CBE cassette and allows its integration into the genome using the dedicated transposase (orange). The pMT85_SpdCas9_pmcDA1 plasmid (for *Mbov* experiments) has inverted repeat sequences and contains two resolvase sequences (Res). These Res sequences allow elimination of the deaminase cassette and the antibiotic resistance marker using an *oriC* plasmid that encodes resolvase activity (6). The pMYCO1_SpdCas9_pmcDA1 (for *Mmm* experiments) is a replicative plasmid containing an *ori*C region from *Mycoplasma mycoides* subsp. *capri* (*ori*C *Mmc,* yellow).

A

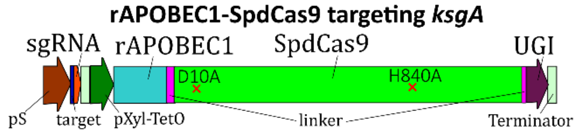


B

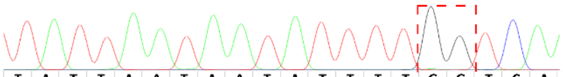

| | T | A | T | T | A | A | T | A | A | T | A | T | T | T | T | G | G | T | C | A |
|---|---|---|---|---|---|---|---|---|---|---|---|---|---|---|---|---|---|---|---|---|
| **T** | 100 | 1 | 100 | 100 | 0 | 0 | 97 | 0 | 0 | 100 | 0 | 100 | 100 | 100 | 100 | 0 | 0 | 100 | 0 | 0 |
| **G** | 0 | 0 | 0 | 0 | 0 | 0 | 0 | 0 | 0 | 0 | 0 | 0 | 0 | 0 | 0 | 98 | 98 | 0 | 0 | 0 |
| **C** | 0 | 0 | 0 | 0 | 0 | 0 | 0 | 0 | 0 | 0 | 0 | 0 | 0 | 0 | 0 | 0 | 0 | 0 | 100 | 0 |
| **A** | 0 | 99 | 0 | 0 | 100 | 100 | 3 | 100 | 100 | 0 | 100 | 0 | 0 | 0 | 0 | 2 | 2 | 0 | 0 | 100 |

**Figure S3. Targeting of the *ksgA* gene in *Mgal* using the rAPOBEC1deaminase-encoding plasmid. A-** Diagram of the CBE based on the rAPOBEC1 deaminase. The sgRNA (red) expression cassette includes a 20-nucleotide target spacer (dark blue) under the control of the spiralin promoter (in brown). The P*xyl/tetO2* inducible promoter (dark green) drives the expression of a codon-optimized hybrid protein fusing *S. pyogenes* dead Cas9 (SpdCas9, green) (inactivated by 2 mutations at positions 10 and 840) and linkers (purple) with the rAPOBEC1 deaminase protein (light blue) and UGI (uracil glycosylase inhibitor (dark purple)). Fibril terminators from *S. citri* (light green) are present downstream of the sgRNA and CBE-encoding gene. **B-** The percentage of bases found in the population were determined from Sanger sequencing chromatograms using EditR software and are represented in the table for each nucleotide position in the target sequence.

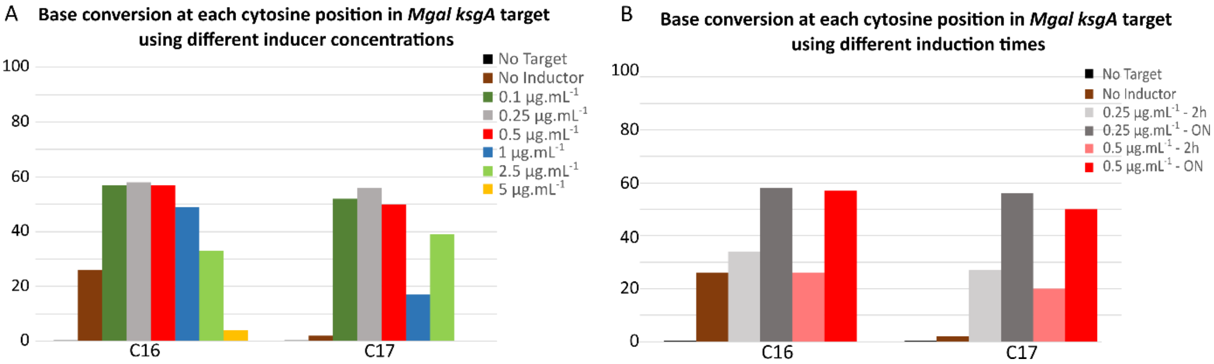


**Figure S4. Optimization of CBE induction conditions for base editing in *Mgal*. A-** Identification of optimal inducer concentrations for the induction of base editing. The percentage of C to T base conversion at cytosine positions 16 and 17 in the target were measured after overnight induction with 0, 0.1, 0.25, 0.5, 1, 2.5, or 5 µg.mL$^{-1}$ aTC. **B-** Identification of the optimal induction times for maximum efficiency. The percentage of C to T base conversion at cytosine positions 16 and 17 in the target is shown after a 2-h or overnight induction.

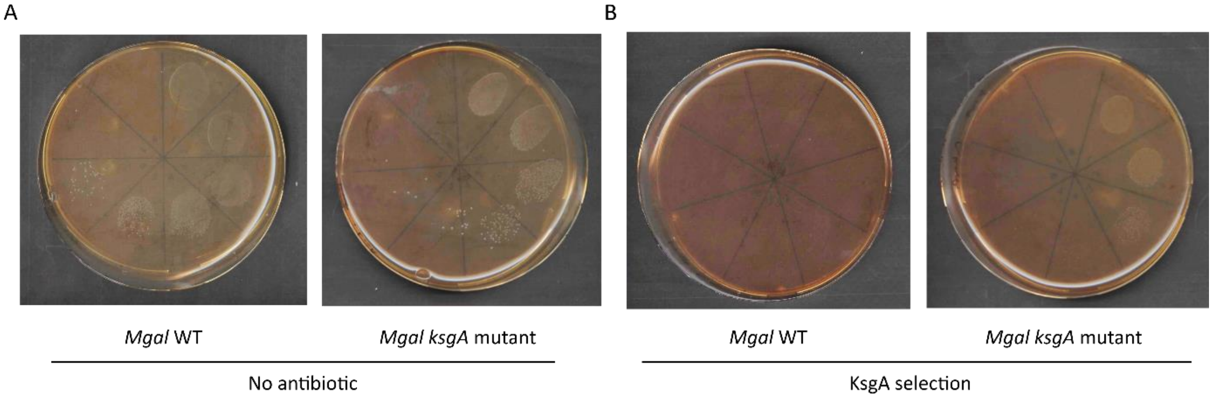


**Figure S5. Kasugamycin resistance phenotypic assay on a *Mgal_ksgA* mutant. A-** Growth of WT or mutant *Mgal_ksgA* was evaluated in the absence (A) or presence of kasugamycin (B) using diluted cell suspensions ($10^{-1}$ to $10^{-7}$).

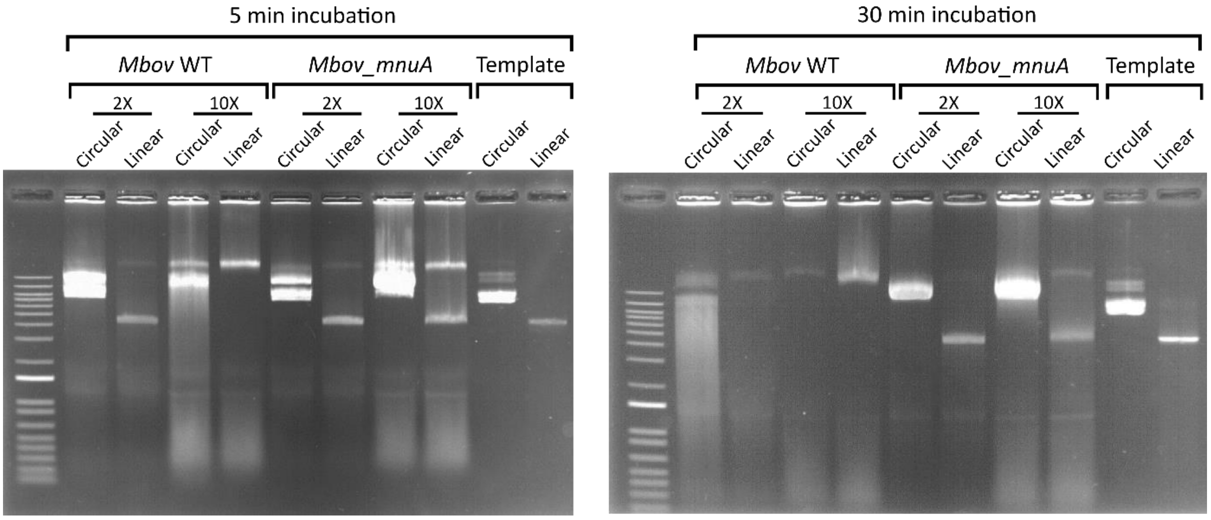


**Figure S6. *Mbov_mnuA* DNase phenotypic assay.** *Mbov* Wt and the *Mbov_mnuA* mutant were incubated with 2 µg linear or circular DNA. After 5 (left) or 30 min (right) at 37°C, aliquots were removed and migrated on 1% agarose gels. DNA digestion was visualized after staining the gels with ethidium bromide. Two concentrations of *Mbov* cells (2X and 10X) were used, as described in Materials and Methods.

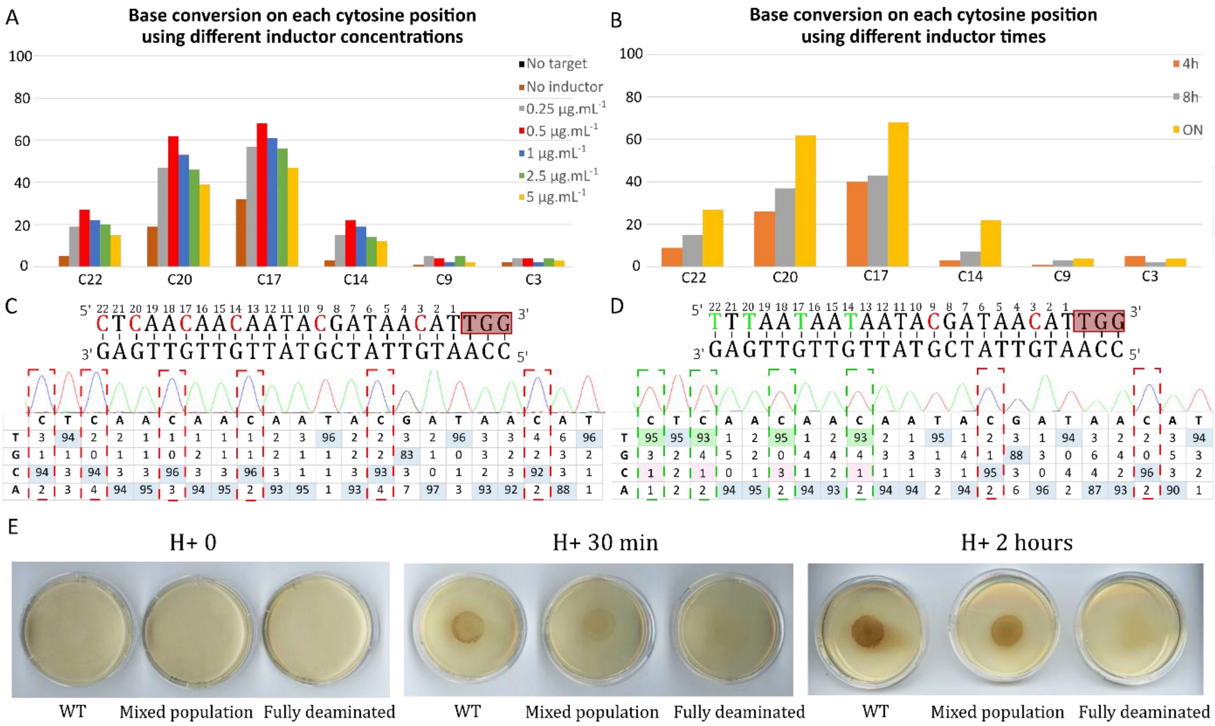


**Figure S7. Targeting the *glpO* gene in *Mmm* using the mycoplasma CBE system. A-** Identification of optimal inducer concentration for the induction of base editing. The percentage of C to T base conversion was calculated at each cytosine position in the target sequence following overnight induction with 0, 0.25, 0.5, 1, 2.5, or 5 µg.mL$^{-1}$ aTC. **B-** Efficiency of cytosine deamination after induction for 4 h, 8 h, or overnight using 0.5 µg/mL aTC. **C, D-** Results of *glpO* editing. The target sequences used are represented; cytosines susceptible to deamination are shown in red and the position of the nucleotide before PAM sequencing is indicated above each nucleotide. These results represent Sanger sequencing chromatograms and EditR analyses before induction (**C**) and after clone isolation (**D**). **E-** $H_2O_2$ detection using a DAB (3,3′-diaminobenzidine) assay on three *Mmm* clones, wildtype (WT) cells, a mixed population of *Mmm* cells with incomplete deamination, and a clone fully deaminated at positions $C_{14}$, $C_{17}$, $C_{20}$, and $C_{22}$. The results were observed at various times after adding the reagent: 0 min, 30 min, and 2 h of incubation.

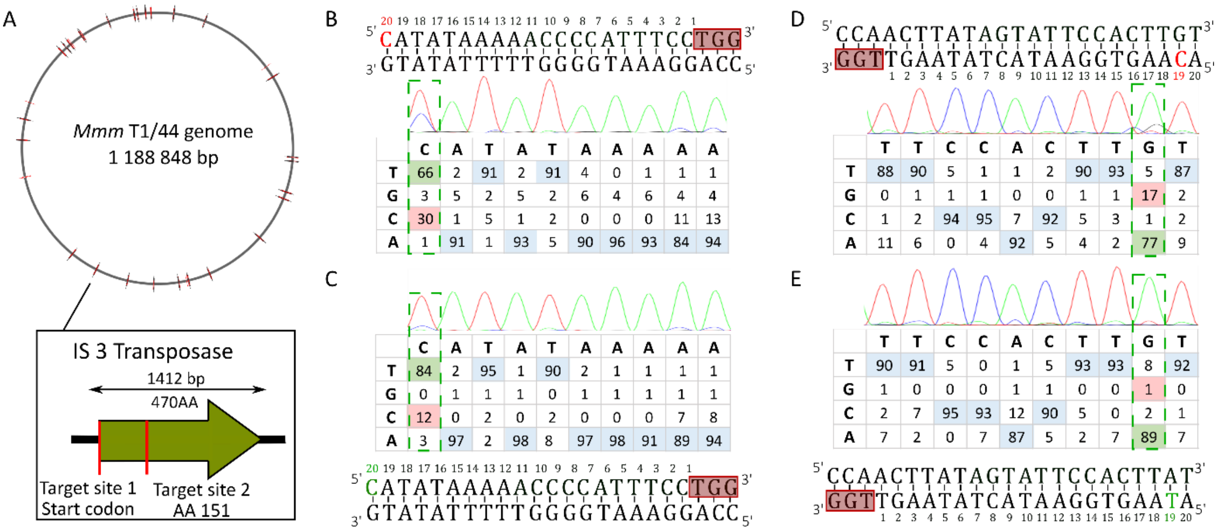


**Figure S8. Multitargeting of IS*3* copies in the *Mmm* genome. A-** Schematic representation of the *Mmm* T1/44 genome, with 30 complete or truncated copies of the IS*3* transposases represented as solid lines. In the right panel, the targeted site within the IS*3* transposase gene is represented as a red bar. **B, D-** Population screening was performed by PCR and Sanger sequencing after three steps of induction using sgRNA1 (**B**) or sgRNA2 (**D**). Targeted cytosines are indicated in red. The percentage of each base at each position is shown in the tables. **C, E-** Sequence profiles of fully mutated clones on sites targeted with sgRNA1 (**C**) and sgRNA2 (**E**). Mutated positions are framed in green in the tables and are shown in green in the target sequences.

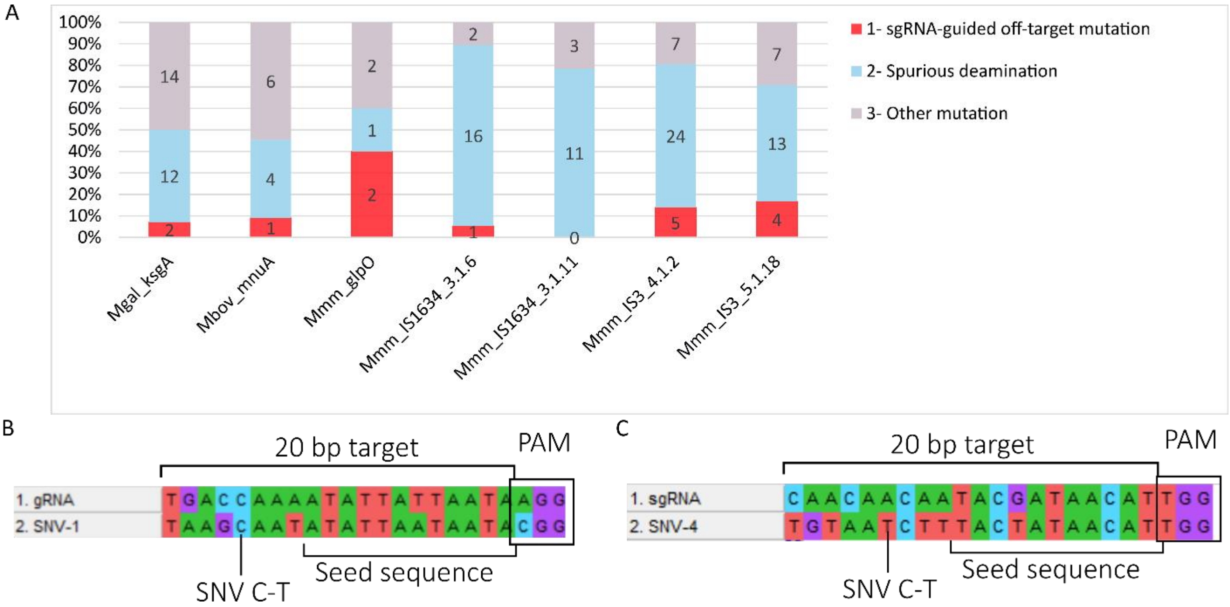


**Figure S9. Analysis of undesired mutations from whole genome sequences. A-** Histogram summarizing the percentage of different categories of mutations predicted from analyses of whole genome sequencing data on seven mycoplasma mutants. The number of mutations is indicated within the bars: blue, sgRNA-guided off-target mutations; orange, spurious deaminations; grey, other mutations. **B-** Example of an off-target interaction of the *ksgA* sgRNA targeting position 1,313 of the *Mgal* genome. **C-** Example of an off-target interaction of the *glpO* sgRNA targeting position 220,532 in the *Mmm* genome.

**Supplementary text**

**SI-1. Materials and Methods**

***In silico* design of codon-optimized base-editor systems.** DNA sequences encoding dead SpCas9, pmcDA1, rAPOBEC1, and UGI, as well as the puromycin resistance marker (linked to SynMYCO promoter), were codon optimized with the Optimizer online tool (http://genomes.urv.es/OPTIMIZER/) using the codon usage of *Mgal* from the Kazusa database (https://www.kazusa.or.jp/codon/). The sgRNA-encoding gene was cloned between the promoter of the *spiralin* gene and the terminator of the *fibril* gene from *Spiroplasma citri* (Figure S1). Sp dCas9 was fused to the rAPOBEC1 sequence by a 48-bp linker used in the BE4 system (1). Sp dCas9 was fused to the pmcDA1 sequence using the same 207-bp linker as that used in the hAID system (2). UGI was fused to the N-terminal sequence using a 12-bp linker (1). The inducible promoter P*xyl/tetO2* (3) and the terminator of the fibril gene were added upstream and downstream, respectively, of the CBE sequence. Manual modifications to avoid regions with AT-rich tracks or unwanted BseRI restriction sites were performed before synthesis of the DNA fragments by Twist Bioscience.

**Plasmid construction.** The Mini-Tn4001 (4) transposon vector derived from Tn*4001* (5) and the synthesized fragments provided by Twist Biosciences were used as PCR templates. The Q5 High-Fidelity DNA Polymerase kit (NEB, M0491) was used for all PCR reactions. Primers A1/A2 (SI) were used to amplify the plasmid backbone encoding the transposon and the tetracycline repressor, primers A3/A4 the synthetized fragment encoding the sgRNA and the first 665 amino-acids of the hybrid protein dCas9-deaminase-UGI, and primers A5/A6 the synthesized fragment encoding the last 1067 amino-acids of the dCas9-deaminase-UGI hybrid protein and the puromycin resistance marker. PCR products were incubated with the DpnI restriction enzyme (NEB, R0176S) following the manufacturer's recommendations. Both products were purified using the GFX™ PCR DNA or Gel Band Purification Kit (GE-healthcare). The NEBuilder HiFi DNA Assembly Cloning Kit (NEB, E5520S) was used to assemble the DNA fragments and create the circular plasmids pTi4.0_spCas9_rAPOBEC1 and pTi4.0_ SpCas9_pmcDA1. *E. coli* NEB 5α (NEB, C2987H) was transformed with 2 µL of the assembled constructs. Transformants were screened after DNA extraction with the NucleoSpin Plasmid kit (Macherey-nagel, 740588.50) and enzymatic digestion. Sanger sequencing of the targeted locus was performed (Genewiz) for final verification.

For the plasmid pMT85_spCas9-pmcDA1, the pMT85-2Res-Genta backbone (6) was used instead of the Mini-Tn4001 transposon backbone. Primers G38/G39 were used to amplify a fragment containing the sgRNA, deaminase complex, and tetracycline repressor and primers G36/G37 were used to amplify the pMT85-2Res-Genta backbone. Cloning was performed using the same procedure as for pTi4.0 (see above).

For the plasmid pMYCO1_spCas9-pmcDA1, the pMYCO1 backbone (*oriC* plasmid (7)) was used instead of the transposon backbone. Primers D27/D28 were used to amplify a fragment containing the sgRNA, deaminase complex, tetracycline repressor, and puromycin resistance marker and primers D29/D30 were used to amplify the pMYCO1 backbone. Cloning was performed using the same procedure as for pTi4.0 (see above).

For all CBE constructs, 20-nucleotide targets were added using the same process. The targets were designed as oligonucleotides R and F. Ten microliters of each primer (100 µM) was heated with 2 µL Adv2 polymerase buffer (TAKARA, 639232) at 95°C for 5 min and slowly cooled (-0.1°C/sec) to room temperature. Plasmids were digested with BseRI (NEB, R0581S) for 2 h at 37°C. After purification, target sequences and plasmids were ligated using T4 DNA ligase (Promega) overnight at 4°C. NEB 5α competent *E. coli* (NEB, C2987H) was transformed with 2 µL of the ligation mix. Colonies were screened by digestion of the plasmids with BseRI and KpnI restriction enzymes and the target sequences verified by Sanger sequencing (Genewiz).

**Induction of CBE.** For each mycoplasma species, transformants were picked and grown in selective liquid media for three passages (one passage is equivalent to a 1:100 dilution). At passage 3, fresh aTC (Abcam, ab145350) (in ETOH 50%) was added to the culture at early logarithmic growth phase (~10 h for *Mbov* and *Mmm*, ~24 h for *Mgal*) and the cells grown until the stationary phase was reached (12 h for *Mbov*, 18 h for *Mmm*, 24 h for *Mgal*). A concentration of 0.5 $\mu g.mL^{-1}$ aTC was used in the three mycoplasma species. After induction, cells were plated on selective solid medium. Isolated clones were obtained after incubation at 37°C for 3 to 10 days. Alternatively, induction was performed immediately after transformation. In this case, after the 2 h incubation step at 37°C (recovery of the cells), antibiotics (puromycin or gentamicin) were added to the media and the cultures incubated for 2 h. Then, the base-editor system was induced using fresh aTC (0.5 $\mu g.mL^{-1}$) for 12 h to 15 h (overnight). Induced cultures were plated on selective media and incubated at 37°C with 5% $CO_2$.

**PCR screening of transformants.** PCR screening was performed on all transformants before induction, after induction, or on isolated clones using the Advantage HF 2 PCR Kit (Takara, 639123). A list of all primers used to screen sites of deamination is available in Table S1.

**EditR analysis.** EdiTR 1.0.10 software (https://moriaritylab.shinyapps.io/editr_v10/) was used to analyze and quantify base editing at each position of the 20-nucleotide target (8). The analysis was performed using the .ab files available after Sanger sequencing.

**Whole genome sequencing of mycoplasmas.** Genomic DNA of *Mgal* was extracted from a 10-mL culture using the Qiagen Genomic-Tips 100/G kit. Genome sequencing was performed by the Genome Transcriptome Facility of Bordeaux. Long reads were produced using a GridION device (Oxford Nanopore) and short reads a MiSeq device (Illumina). For *Mmm_glpO* mutant cl18_4, ONT sequencing generated 24,400 reads (Mean read length: 29,377 bp) and Illumina 1,044,350 read pairs. Analyses were performed using Galaxy (https://usegalaxy.eu/). Mutations were detected after mapping onto the *Mmm T1/44* genome (CP014346.1). Illumina reads were trimmed using Trimmomatic (V 0.38.1; Sliding Window 10, 20; Drop read below minimal length of 250), mapped using BWA-MEM (V 0.7.17.1), Samtools sort (V 2.0.3), and MPileup ( V 2.1.1), and variants detected using VarScan mpileup (V 2.4.3.1; Minimum coverage 30, Minimum supporting read 20, Minimum Base quality 30, Minimum variant allele frequency 0.8, Minimum homozygous variants 0.75). Mutations are shown in Table S3. Genome assembly was performed using the following steps: ONT reads were filtered using Filter FASTQ (V 1.1.5, Minimum size 45,000 bp), assembled using Flye Assembly (V 2.6), and polished using four rounds of Pilon (1.20.1) combined with Illumina short reads. The assembled genome was compared to the *Mmm* T1/44 (CP014346.1) reference genome using MAUVE software (9). For the *Mgal_ksgA* mutant, the same procedure was used with 45,406 ONT long-reads (Mean read length: 23,295 bp) and 851,806 Illumina short-read pairs using the *Mgal*_S6 reference genome (CP006916.3). For the detection of SNVs in *Mgal*, the results were obtained by comparison with a previously sequenced clone (accession number PRJNA769398) to eliminate mutations that are present in the laboratory clone. For *Mbov*_WT, the same procedure was used with 28,729 ONT long reads (Mean read length: 27,460 bp) and 881,705 Illumina short-read pairs. Mutations of our laboratory strain were detected after mapping the *Mbov*_PG45 reference genome (CP002188.1). For the *Mbov_mnuA* mutant cl3, the same procedure was used with 42,506 ONT long reads (Mean read length: 24,459 bp) and 910,742 Illumina short-read pairs. Mutations were detected after mapping using the *Mbov*_PG45 reference genome (CP002188.1). For the detection of SNVs in *Mbov* linked to our CBE, the genome sequence of *Mbov*_WT was used to eliminate mutations that are present in the laboratory strain. For the four *Mmm*_IS mutants, the same procedure was used. Sequencing of *Mmm*_IS3 mutant cl 5.1.18 generated 54,523 ONT long reads (Mean read length: 22,424 bp) and 941,944 Illumina short-read pairs. Sequencing of *Mmm*_IS3 mutant cl 4.1.2 generated 57,466 ONT long-reads (Mean read length: 20,074 bp) and 778,172 Illumina short-read pairs. Sequencing of *Mmm*_IS1634 mutant cl 3.1.11 generated 67,316 ONT long-reads (Mean read length: 22,647 bp) and 724,029 Illumina short-read pairs. Finally, sequencing of *Mmm*_IS1634 mutant cl 3.1.6 generated 31,875 ONT long-reads (Mean read length: 24,551 bp) and 1,199,725 Illumina short-read pairs. Mutations were detected in these four clones after mapping using the *Mmm* T1/44 reference genome (CP014346.1).

SNPs found in all sequenced clones were classified to identify their potential origin. A summary can be found in Table S3. For sgRNA-mediated SNPs, a Blast of the target sequence against the DNA region around the SNP was performed. Two conditions were established to consider an SNV as an sgRNA-dependent off-target mutation: (1) the presence of the PAM sequence and (2) at least 70% sequence similarity for the first 12 nucleotides (considered as the Seed-sequence). For other mutations, C to T or G to A SNPs were considered to result from spurious deamination and the nucleotide before was observed to be a TC motif, which are preferred by the deaminase protein.

**Phenotypic assay for *Mgal*: kasugamycin resistance.** *Mgal* S6 and the mutant *Mgal_ ksgA* were cultured for 36 h. Serial dilutions were performed down to $10^{-7}$. Each dilution (20 µL) was plated onto Hayflick medium, with or without kasugamycin, and incubated for 10 days at 37°C (5% $CO_2$).

**Phenotypic assay for *Mbov*: nuclease activity.** Nuclease activity assays on WT *M. bovis* PG45 and the mutant *Mbov_mnuA* were performed as previously described in Sharma *et al*. 2015 (10) with several modifications. Briefly, for each strain, 2 mL of culture was prepared in SP4 medium. At late log phase, cultures were divided in half and centrifuged for 10 min at 7,000 x *g* at 10°C. One 1-mL sample was mixed with 500 µL nuclease buffer (25 mM Tris-HCl, pH 8.8, 10 mM $CaCl_2$, 10 mM $MgCl_2$) and the other 1-mL sample was mixed with 100 µL nuclease buffer to concentrate the mycoplasma cells (2 and 10 times, respectively). Then, 2 µg circular plasmid DNA (pMT85_SpCas9_pmcDA1) or 500 ng double stranded linear DNA (PCR fragment) was incubated with 50 µL of each sample at 37°C for 5 or 60 min. At each time point, 10-µL aliquots were removed and the reaction was stopped by the addition of EDTA to a final concentration of 20 mM. Aliquots were mixed with 6X loading buffer (Promega) and immediately loaded onto 1% agarose gels made in 1X TAE buffer.

**Phenotypic assay for *Mmm*: $H_2O_2$ production.** *Mmm* T1/44 WT and *glpO* mutants were grown until reaching a pH of 7. A 20-µL drop of each culture was then plated on PPLO media and the plates incubated for 48 h. GlpO activity was evaluated using a qualitative "on-the-plate" $H_2O_2$ test that allows the detection of $H_2O_2$ production in response to the addition of a glycerol-containing reaction mix (11). Briefly, 40 µL of glycerol reaction mix ($H_2O$ qsp 1.6 mL + DAB [3,3'-Diaminobenzidine, 9.6 mg] + 64 µL HCl [1N] + horseradish peroxidase [1.6 mg] + 64 µL PDS stock [$CaCl_2$ (1.5 $mg.mL^{-1}$), KCl (1.5 $mg.mL^{-1}$), NaHCO3 (0.625 $mg.mL^{-1}$), NaCl (56.25 $mg.mL^{-1}$)]) was carefully spread onto bacterial lawns and the colorimetric reaction observed after 30 min or 2 h of incubation at 37°C. *Mycoplasma* colonies producing $H_2O_2$ adopt a red-brown color.

### Supplementary References

1. A. C. Komor, *et al.*, Improved base excision repair inhibition and bacteriophage Mu Gam protein yields C:G-to-T:A base editors with higher efficiency and product purity. *Sci. Adv.* **3**, 1–10 (2017).

2. K. Nishida, *et al.*, Targeted nucleotide editing using hybrid prokaryotic and vertebrate adaptive immune systems. *Science (80-. ).* **353** (2016).
3. M. Breton, *et al.*, First report of a tetracycline-inducible gene expression system for mollicutes. *Microbiology* **156**, 198–205 (2010).
4. I. Pour-El, C. Adams, F. C. Minion, Construction of mini-Tn4001tet and its use in *Mycoplasma gallisepticum*. *Plasmid* **47**, 129–137 (2002).
5. Clyde A. Hutchison, *et al.*, Global Transposon Mutagenesis and a Minimal Mycoplasma Genome. *Science (80-. ).* **286** (1999).
6. C. Janis, *et al.*, Unmarked insertional mutagenesis in the bovine pathogen *Mycoplasma mycoides* subsp. *mycoides* SC. *Microbiology* **154**, 2427–2436 (2008).
7. C. Lartigue, A. Blanchard, J. Renaudin, F. Thiaucourt, P. Sirand-Pugnet, Host specificity of mollicutes oriC plasmids: Functional analysis of replication origin. *Nucleic Acids Res.* **31**, 6610–6618 (2003).
8. M. G. Kluesner, *et al.*, EditR: A Method to Quantify Base Editing from Sanger Sequencing. *Cris. J.* **1**, 239–250 (2018).
9. A. C. E. Darling, B. Mau, F. R. Blattner, N. T. Perna, Mauve: Multiple alignment of conserved genomic sequence with rearrangements. *Genome Res.* **14**, 1394–1403 (2004).
10. S. Sharma, K. A. Tivendale, P. F. Markham, G. F. Browning, Disruption of the membrane nuclease gene (MBOVPG45_0215) of *Mycoplasma bovis* greatly reduces cellular nuclease activity. *J. Bacteriol.* **197**, 1549–1558 (2015).
11. P. Rice, B. M. Houshaymi, E. A. M. Abu-Groun, R. A. J. Nicholas, R. J. Miles, Rapid screening of H2O2 production by Mycoplasma mycoides and differentiation of European subsp. mycoides SC (small colony) isolates. *Vet. Microbiol.* **78**, 343–351 (2001).

### SI-2. Genes and plasmids sequences

#### Nucleic sequence of original SpdCas9 (KJ796484.1)

```
ATGGACAAGAAGTACTCCATTGGGCTCGCTATCGGCACAAACAGCGTCGGTTGGGCCGTC
ATTACGGACGAGTACAAGGTGCCGAGCAAAAAATTCAAAGTTCTGGGCAATACCGATCGC
CACAGCATAAAGAAGAACCTCATTGGCGCCCTCCTGTTCGACTCCGGGGAGACGGCCGAA
GCCACGCGGCTCAAAAGAACAGCACGGCGCAGATATACCCGCAGAAAGAATCGGATCTGC
TACCTGCAGGAGATCTTTAGTAATGAGATGGCTAAGGTGGATGACTCTTTCTTCCATAGG
CTGGAGGAGTCCTTTTTGGTGGAGGAGGATAAAAAGCACGAGCGCCACCCAATCTTTGGC
AATATCGTGGACGAGGTGGCGTACCATGAAAAGTACCCAACCATATATCATCTGAGGAAG
AAGCTTGTAGACAGTACTGATAAGGCTGACTTGCGGTTGATCTATCTCGCGCTGGCGCAT
ATGATCAAATTTCGGGGACACTTCCTCATCGAGGGGGACCTGAACCCAGACAACAGCGAT
GTCGACAAACTCTTTATCCAACTGGTTCAGACTTACAATCAGCTTTTCGAAGAGAACCCG
ATCAACGCATCCGGAGTTGACGCCAAAGCAATCCTGAGCGCTAGGCTGTCCAAATCCCGG
CGGCTCGAAAACCTCATCGCACAGCTCCCTGGGGAGAAGAAGAACGGCCTGTTTGGTAAT
CTTATCGCCCTGTCACTCGGGCTGACCCCCAACTTTAAATCTAACTTCGACCTGGCCGAA
GATGCCAAGCTTCAACTGAGCAAAGACACCTACGATGATGATCTCGACAATCTGCTGGCC
CAGATCGGCGACCAGTACGCAGACCTTTTTTTGGCGGCAAAGAACCTGTCAGACGCCATT
CTGCTGAGTGATATTCTGCGAGTGAACACGGAGATCACCAAAGCTCCGCTGAGCGCTAGT
ATGATCAAGCGCTATGATGAGCACCACCAAGACTTGACTTTGCTGAAGGCCCTTGTCAGA
CAGCAACTGCCTGAGAAGTACAAGGAAATTTTCTTCGATCAGTCTAAAAATGGCTACGCC
GGATACATTGACGGCGGAGCAAGCCAGGAGGAATTTTACAAATTTATTAAGCCCATCTTG
GAAAAAATGGACGGCACCGAGGAGCTGCTGGTAAAGCTTAACAGAGAAGATCTGTTGCGC
AAACAGCGCACTTTCGACAATGGAAGCATCCCCCACCAGATTCACCTGGGCGAACTGCAC
GCTATCCTCAGGCGGCAAGAGGATTTCTACCCCTTTTTGAAAGATAACAGGGAAAAGATT
GAGAAAATCCTCACATTTCGGATACCCTACTATGTAGGCCCCCTCGCCCGGGGAAATTCC
AGATTCGCGTGGATGACTCGCAAATCAGAAGAGACCATCACTCCCTGGAACTTCGAGGAA
GTCGTGGATAAGGGGGCCTCTGCCCAGTCCTTCATCGAAAGGATGACTAACTTTGATAAA
AATCTGCCTAACGAAAAGGTGCTTCCTAAACACTCTCTGCTGTACGAGTACTTCACAGTT
TATAACGAGCTCACCAAGGTCAAATACGTCACAGAAGGGATGAGAAAGCCAGCATTCCTG
TCTGGAGAGCAGAAGAAAGCTATCGTGGACCTCCTCTTCAAGACGAACCGGAAAGTTACC
GTGAAACAGCTCAAAGAAGACTATTTCAAAAAGATTGAATGTTTCGACTCTGTTGAAATC
AGCGGAGTGGAGGATCGCTTCAACGCATCCCTGGGAACGTATCACGATCTCCTGAAAATC
ATTAAAGACAAGGACTTCCTGGACAATGAGGAGAACGAGGACATTCTTGAGGACATTGTC
CTCACCCTTACGTTGTTTGAAGATAGGGAGATGATTGAAGAACGCTTGAAAACTTACGCT
CATCTCTTCGACGACAAAGTCATGAAACAGCTCAAGAGGCGCCGATATACAGGATGGGGG
CGGCTGTCAAGAAAACTGATCAATGGGATCCGAGACAAGCAGAGTGGAAAGACAATCCTG
GATTTTCTTAAGTCCGATGGATTTGCCAACCGGAACTTCATGCAGTTGATCCATGATGAC
TCTCTCACCTTTAAGGAGGACATCCAGAAAGCACAAGTTTCTGGCCAGGGGGACAGTCTT
CACGAGCACATCGCTAATCTTGCAGGTAGCCCAGCTATCAAAAAGGGAATACTGCAGACC
GTTAAGGTCGTGGATGAACTCGTCAAAGTAATGGGAAGGCATAAGCCCGAGAATATCGTT
ATCGAGATGGCCCGAGAGAACCAAACTACCCAGAAGGGACAGAAGAACAGTAGGGAAAGG
ATGAAGAGGATTGAAGAGGGTATAAAAGAACTGGGGTCCCAAATCCTTAAGGAACACCCA
GTTGAAAACACCCAGCTTCAGAATGAGAAGCTCTACCTGTACTACCTGCAGAACGGCAGG
GACATGTACGTGGATCAGGAACTGGACATCAATCGGCTCTCCGACTACGACGTGGATGCT
ATCGTGCCCCAGTCTTTTCTCAAAGATGATTCTATTGATAATAAAGTGTTGACAAGATCC
GATAAAAATAGAGGGAAGAGTGATAACGTCCCCTCAGAAGAAGTTGTCAAGAAAATGAAA
AATTATTGGCGGCAGCTGCTGAACGCCAAACTGATCACACAACGGAAGTTCGATAATCTG
```

ACTAAGGCTGAACGAGGTGGCCTGTCTGAGTTGGATAAAGCCGGCTTCATCAAAAGGCAG
CTTGTTGAGACACGCCAGATCACCAAGCACGTGGCCCAAATTCTCGATTCACGCATGAAC
ACCAAGTACGATGAAAATGACAAACTGATTCGAGAGGTGAAAGTTATTACTCTGAAGTCT
AAGCTGGTCTCAGATTTCAGAAAGGACTTTCAGTTTTATAAGGTGAGAGAGATCAACAAT
TACCACCATGCGCATGATGCCTACCTGAATGCAGTGGTAGGCACTGCACTTATCAAAAAA
TATCCCAAGCTTGAATCTGAATTTGTTTACGGAGACTATAAAGTGTACGATGTTAGGAAA
ATGATCGCAAAGTCTGAGCAGGAAATAGGCAAGGCCACCGCTAAGTACTTCTTTTACAGC
AATATTATGAATTTTTTCAAGACCGAGATTACACTGGCCAATGGAGAGATTCGGAAGCGA
CCACTTATCGAAACAAACGGAGAAACAGGAGAAATCGTGTGGGACAAGGGTAGGGATTTC
GCGACAGTCCGGAAGGTCCTGTCCATGCCGCAGGTGAACATCGTTAAAAAGACCGAAGTA
CAGACCGGAGGCTTCTCCAAGGAAAGTATCCTCCCGAAAAGGAACAGCGACAAGCTGATC
GCACGCAAAAAAGATTGGGACCCCAAGAAATACGGCGGATTCGATTCTCCTACAGTCGCT
TACAGTGTACTGGTTGTGGCCAAAGTGGAGAAAGGGAAGTCTAAAAAACTCAAAAGCGTC
AAGGAACTGCTGGGCATCACAATCATGGAGCGATCAAGCTTCGAAAAAAACCCCATCGAC
TTTCTCGAGGCGAAAGGATATAAAGAGGTCAAAAAAGACCTCATCATTAAGCTTCCCAAG
TACTCTCTCTTTGAGCTTGAAAACGGCCGGAAACGAATGCTCGCTAGTGCGGGCGAGCTG
CAGAAAGGTAACGAGCTGGCACTGCCCTCTAAATACGTTAATTTCTTGTATCTGGCCAGC
CACTATGAAAAGCTCAAAGGGTCTCCCGAAGATAATGAGCAGAAGCAGCTGTTCGTGGAA
CAACACAAACACTACCTTGATGAGATCATCGAGCAAATAAGCGAATTCTCCAAAAGAGTG
ATCCTCGCCGACGCTAACCTCGATAAGGTGCTTTCTGCTTACAATAAGCACAGGGATAAG
CCCATCAGGGAGCAGGCAGAAAACATTATCCACTTGTTTACTCTGACCAACTTGGGCGCG
CCTGCAGCCTTCAAGTACTTCGACACCACCATAGACAGAAAGCGGTACACCTCTACAAAG
GAGGTCCTGGACGCCACACTGATTCATCAGTCAATTACGGGGCTCTATGAAACAAGAATC
GACCTCTCTCAGCTCGGTGGAGAC

### Nucleic sequence of original rAPOBEC1 (NM_012907.2)

ATGAGTTCCGAGACAGGCCCTGTAGCTGTTGATCCCACTCTGAGGAGAAGAATTGAGCCC
CACGAGTTTGAAGTCTTCTTTGACCCCCGGGAACTTCGGAAAGAGACCTGTCTGCTGTAT
GAGATCAACTGGGGAGGAAGGCACAGCATCTGGCGACACACGAGCCAAAACACCAACAAA
CACGTTGAAGTCAATTTCATAGAAAAATTTACTACAGAAAGATACTTTTGTCCAAACACC
AGATGCTCCATTACCTGGTTCCTGTCCTGGAGTCCCTGTGGGGAGTGCTCCAGGGCCATT
ACAGAATTTTTGAGCCGATACCCCCATGTAACTCTGTTTATTTATATAGCACGGCTTTAT
CACCACGCAGATCCTCGAAATCGGCAAGGACTCAGGGACCTTATTAGCAGCGGTGTTACT
ATCCAGATCATGACGGAGCAAGAGTCTGGCTACTGCTGGAGGAATTTTGTCAACTACTCC
CCTTCGAATGAAGCTCATTGGCCAAGGTACCCCCATCTGTGGGTGAGGCTGTACGTACTG
GAACTCTACTGCATCATTTTAGGACTTCCACCCTGTTTAAATATTTTAAGAAGAAAACAA
CCTCAACTCACGTTTTTCACGATTGCTCTTCAAAGCTGCCATTACCAAAGGCTACCACCC
CACATCCTGTGGGCCACAGGGTTGAAATAA

### Nucleic sequence of original pmcDA1 (ABO15149.1)

ATGACAGACGCCGAGTACGTGCGCATTCATGAGAAACTGGATATTTACACCTTCAAGAAG
CAGTTCTTCAACAACAAGAAATCTGTGTCACACCGCTGCTACGTGCTGTTTGAGTTGAAG
CGAAGGGGCGAAAGAAGGGCTTGCTTTTGGGGCTATGCCGTCAACAAGCCCCAAAGTGGC
ACCGAGAGAGGAATACACGCTGAGATATTCAGTATCCGAAAGGTGGAAGAGTATCTTCGG
GATAATCCTGGGCAGTTTACGATCAACTGGTATTCCAGCTGGAGTCCTTGCGCTGATTGT
GCCGAGAAAATTCTGGAATGGTATAATCAGGAACTTCGGGGAAACGGGCACACATTGAAA
ATCTGGGCCTGCAAGCTGTACTACGAGAAGAATGCCCGGAACCAGATAGGACTCTGGAAT
CTGAGGGACAATGGTGTAGGCCTGAACGTGATGGTTTCCGAGCACTATCAGTGTTGTCGG
AAGATTTTCATCCAAAGCTCTCATAACCAGCTCAATGAAAACCGCTGGTTGGAGAAAACA
CTGAAACGTGCGGAGAAGCGGAGATCCGAGCTGAGCATCATGATCCAGGTCAAGATTCTG
CATACCACTAAGTCTCCAGCCGTTTAA

### Nucleic sequence of optimized APOBEC1

ATGAGTAGTGAAACTGGACCAGTTGCTGTTGACCCCACTCTCAGAAGAAGAATAGAACCA
CATGAATTTGAAGTATTCTTCGATCCAAGAGAATTAAGAAAAGAAACTTGTTTATTATAT
GAAATTAATTGGGGAGGAAGACATAGTATTTGGAGACATACTAGTCAAAATACTAATAAA
CATGTTGAAGTTAATTTTATTGAGAAGTTTACTACTGAAAGATATTTCTGTCCAAATACT
AGATGTAGTATTACTTGGTTCTTAAGTTGGAGTCCATGTGGAGAATGTAGTAGAGCTATA
ACTGAATTCCTGAGTAGATATCCACATGTTACTTTATTTATTTATATTGCTAGATTATAT
CATCATGCTGATCCAAGAAATAGACAAGGATTAAGAGATTTAATTAGTAGTGGAGTTACT
ATTCAAATTATGACTGAACAAGAAAGTGGATATTGTTGGAGAAATTTTGTTAATTATAGT
CCAAGTAATGAAGCTCATTGGCCAAGATATCCACATTTATGGGTTAGATTATATGTTTTA
GAATTATATTGTATTATTTTAGGATTACCACCATGTTTAAATATTTTAAGAAGAAAACAA
CCACAATTAACATTCTTCACTATTGCTTTACAAAGTTGTCATTATCAAAGATTACCACCA
CATATTTTATGGGCTACTGGATTAAAATAA

### Nucleic sequence of optimized pmcDA1

ATGACTGATGCTGAATATGTTAGAATTCATGAGAAATTAGATATTTATACTTTTAAGAAA
CAATTCTTTAATAATAAGAAGAGTGTTAGTCATCGTTGCTATGTGCTCTTTGAATTAAAG
AGAAGAGGTGAAAGAAGAGCTTGCTTTTGAGGATATGCTGTTAATAAACCACAAAGTGGA
ACTGAAAGAGGAATTCATGCTGAAATATTTAGTATTAGAAAAGTTGAAGAATATTTAAGA
GATAATCCAGGACAATTTACTATTAATTGGTATAGTAGTTGGAGTCCATGTGCTGATTGT
GCTGAGAAGATTTTAGAATGGTATAATCAAGAATTAAGAGGAAATGGACATACTTTGAAA
ATTTGGGCTTGTAAATTATATTATGAGAAGAATGCTAGAAATCAAATTGGATTATGGAAT
TTAAGAGATAATGGAGTTGGATTAAATGTTATGGTTAGTGAACATTATCAATGTTGTAGA
AAGATATTTATTCAAAGTAGTCATAATCAATTAAATGAAAATAGATGGTTAGAGAAAACT
TTAAAGAGAGCTGAGAAAAGAAGAAGTGAATTAAGTATTATGATTCAAGTTAAAATTTTA

CATACTACTAAAAGTCCAGCTGTTTAA

### Nucleic sequence of optimized SpdCas9

ATGGACAAAAAATATTCTATTGGATTAGCTATTGGAACTAATTCAGTAGGTTGAGCTGTT
ATTACAGATGAATATAAGGTACCATCAAAGAAATTTAAGGTTTTAGGTAATACTGATAGA
CATTCAATTAAAAAAAATTTAATCGGAGCATTACTTTTTGATTCTGGAGAAACAGCAGAA
GCTACCAGATTAAAAAGAACGGCTCGTAGACGATATACTAGACGTAAAAACAGAATCTGT
TATCTTCAAGAAATTTTTAGTAATGAAATGGCTAAAGTTGATGACTCTTTTTTTCACAGA
TTGGAAGAGTCTTTCTTAGTAGAAGAAGATAAAAAGCATGAAAGACACCCCATCTTTGGT
AATATTGTAGATGAAGTCGCCTATCATGAAAAATATCCTACAATTTATCATTTAAGAAAA
AAATTAGTAGATAGCACAGATAAAGCTGATTTAAGATTAATTTATTTAGCACTAGCACAT
ATGATCAAATTTAGAGGTCACTTCTTAATTGAAGGTGATTTAAACCCTGATAATAGTGAC
GTTGATAAATTATTTATTCAATTAGTACAAACGTATAATCAACTTTTCGAAGAAAACCCA
ATTAATGCTAGTGGGGTTGATGCAAAAGCTATCCTTTCGGCTCGTCTTTCAAAATCTAGG
AGACTTGAAAACTTAATTGCACAATTACCGGGAGAGAAAAAAAACGGTTTATTTGGTAAC
TTAATCGCGTTATCTTTAGGTTTAACCCCGAATTTTAAGAGTAATTTCGATCTAGCTGAA
GATGCTAAACTACAATTATCTAAAGATACTTATGACGATGACTTAGATAATTTATTAGCT
CAGATTGGTGATCAATATGCAGACTTATTTTTAGCAGCAAAAAACTTAAGCGACGCAATC
TTATTGAGTGATATATTGAGAGTTAACACAGAAATCACTAAAGCACCATTAAGTGCAAGT
ATGATTAAACGTTATGATGAACACCACCAAGATTTAACACTATTAAAAGCATTAGTTAGA
CAACAATTACCTGAAAAGTATAAAGAAATTTTCTTCGATCAAAGCAAAAATGGTTATGCT
GGTTATATTGATGGTGGAGCTTCACAAGAAGAATTTTATAAGTTCATTAAGCCTATCCTA
GAAAAAATGGATGGAACAGAAGAACTATTAGTCAAGTTAAATCGTGAAGATTTACTACGC
AAACAAAGAACTTTTGATAATGGTAGCATTCCTCATCAAATTCACTTAGGAGAACTACAC
GCTATCCTAAGAAGACAAGAAGATTTTTATCCTTTTTTAAAAGATAATAGAGAAAAAATT
GAAAAAATCTTAACATTTAGAATCCCTTACTATGTAGGTCCGTTAGCTAGAGGAAATAGT
AGATTTGCATGAATGACTCGAAAATCAGAAGAGACTATCACACCATGAAATTTTGAGGAA
GTTGTGGATAAAGGTGCATCTGCGCAATCTTTTATTGAGCGAATGACTAATTTCGATAAG
AACTTACCTAATGAAAAAGTATTACCTAAGCACTCATTATTATATGAATACTTTACTGTT
TATAACGAACTTACTAAAGTAAAATATGTTACCGAAGGAATGAGAAAACCAGCGTTCCTA
AGTGGAGAACAAAAGAAGGCTATTGTTGATTTATTATTTAAGACAAATAGAAAAGTAACT
GTAAAACAACTAAAAGAAGATTATTTTAAAAAAATTGAATGTTTTGATTCAGTCGAAATT
TCTGGAGTTGAAGACCGTTTCAACGCAAGTTTAGGCACTTACCACGATCTACTAAAAATT
ATTAAAGATAAAGATTTTCTTGATAACGAAGAAAATGAAGACATTCTAGAAGATATTGTC
CTAACTTTAACTTTATTCGAAGACAGAGAAATGATTGAAGAAAGATTAAAAACTTACGCT
CACTTATTTGATGATAAAGTTATGAAGCAGTTGAAGCGCCGACGATATACTGGTTGAGGT
AGACTCTCAAGAAAGCTAATCAATGGTATTAGAGACAAACAATCAGGTAAAACAATTTTA
GATTTTTTAAAAAGCGACGGATTTGCTAATAGAAACTTCATGCAATTGATCCACGATGAT
TCATTAACTTTTAAAGAAGATATTCAAAAGGCACAAGTCTCAGGTCAAGGTGATAGCTTA
CACGAACATATCGCTAATCTAGCTGGTTCACCTGCAATCAAAAAGGGAATTTTACAGACA
GTGAAAGTTGTTGATGAACTAGTAAAGGTAATGGGACGCCACAAACCAGAGAACATCGTG
ATTGAAATGGCTAGAGAAAACCAAACAACACAAAAAGGCCAAAAAAACAGTAGAGAAAGA
ATGAAGAGAATCGAAGAAGGTATCAAAGAGTTAGGGTCTCAAATCTTAAAGGAACATCCT
GTTGAAAACACTCAATTACAAAATGAAAAATTATACCTATATTACTTACAAAATGGTAGA
GATATGTATGTTGATCAGGAATTAGATATTAACCGTTTATCAGATTACGACGTGGATGCT
ATTGTACCTCAATCATTTCTAAAAGATGATTCTATCGACAATAAGGTTTTAACTAGATCC
GATAAAAATCGAGGAAAATCTGACAATGTACCTAGTGAAGAAGTTGTAAAAAAGATGAAA
AATTACTGACGACAACTTTTAAACGCAAAATTAATTACACAAAGAAAATTTGATAACCTA
ACTAAAGCAGAGCGtGGAGGTCTGTCTGAACTTGATAAGGCTGGATTTATTAAACGACAA
CTAGTTGAAACACGTCAAATCACCAAACATGTTGCACAAATTTTAGATTCTCGTATGAAT
ACAAAGTACGACGAAAACGATAAATTAATCAGAGAAGTTAAAGTTATTACTCTAAAATCA
AAATTAGTGAGTGATTTTCGCAAAGATTTCCAATTTTACAAGGTTAGAGAGATCAATAAT
TATCATCACGCACATGATGCTTATTTAAATGCTGTGGTTGGGACTGCTTTAATCAAAAAG
TATCCTAAATTAGAAAGCGAATTTGTATACGGTGATTATAAGGTTTATGATGTTCGTAAA
ATGATTGCTAAAAGTGAACAAGAAATTGGAAAGGCTACTGCTAAATATTTTTTTTACTCA
AATATTATGAATTTTTTCAAAACTGAGATCACATTAGCAAATGGTGAAATCCGTAAAAGA
CCTTTAATTGAAACTAACGGGGAAACTGGGGAAATTGTGTGAGATAAAGGTCGTGATTTT
GCAACAGTAAGAAAAGTATTATCAATGCCACAAGTTAATATCGTTAAAAAAACAGAGGTG
CAAACTGGAGGTTTCTCTAAAGAATCGATCTTACCTAAAAGAAACAGTGATAAATTAATT
GCGAGAAAAAAAGATTGAGATCCAAAAAAATATGGTGGTTTTGATTCGCCGACTGTTGCA
TACTCTGTATTAGTGGTTGCTAAAGTTGAAAAAGGTAAGAGTAAAAAATTAAAATCTGTT
AAAGAATTATTAGGCATTACGATTATGGAGAGATCATCTTTTGAAAAAAATCCAATCGAT
TTTTTGGAAGCTAAAGGGTACAAAGAAGTAAAAAAAGATCTTATAATTAAATTACCTAAA
TATTCTTTATTTGAACTTGAAAACGGTCGTAAAAGAATGTTAGCTTCTGCAGGAGAACTA
CAAAAAGGTAATGAATTAGCTCTACCAAGTAAATATGTAAATTTTTTATATTTAGCGAGT
CACTATGAAAAATTAAAAGGATCACCTGAAGATAATGAACAAAAACAATTATTTGTTGAA
CAACACAAACATTATTTAGATGAAATTATTGAACAAATTTCAGAATTTAGCAAAAGAGTT
ATACTTGCCGATGCAAATTTAGATAAAGTGTTAAGTGCGTACAACAAGCACAGAGATAAA
CCAATCCGCGAACAAGCAGAAAATATTATCCACTTATTCACTCTTACTAACTTAGGTGCT
CCAGCAGCTTTTAAATATTTTGATACAACTATCGATAGAAAGAGATATACATCAACAAAG
GAAGTTTTAGACGCGACTTTAATTCATCAAAGTATTACTGGACTTTATGAGACTCGCATC
GATCTATCACAATTGGGTGGAGACTAG

### Nucleic sequence of Puromycin N-acetyltransferase

ATGACTGAATATAAACCTACTGTTAGATTAGCTACTAGAGATGATGTTCCTAGAGCTGTT
AGAACTTTAGCTGCTGCTTTTGCTGATTATCCTGCTACTAGACATACTGTTGATCCTGAT
AGACATATTGAAAGAGTTACTGAATTACAAGAATTATTTTTAACTAGAGTTGGTTTAGAT

ATTGGTAAAGTTTGGGTTGCTGATGATGGTGCTGCTGTTGCTGTTTGGACTACTCCTGAA
AGTGTTGAAGCTGGTGCTGTTTTTGCTGAAATTGGTCCTAGAATGGCTGAATTAAGTGGT
AGTAGATTAGCTGCTCAACAACAAATGGAAGGTTTATTAGCTCCACATAGACCTAAAGAA
CCTGCTTGGTTTTTAGCTACTGTTGGTGTTAGTCCTGATCATCAAGGTAAAGGTTTAGGT
AGTGCTGTTGTTTTACCTGGTGTTGAAGCTGCTGAAAGAGCTGGTGTTCCTGCTTTTTTA
GAAACTAGTGCTCCTAGAAATTTACCTTTTTATGAAAGATTAGGTTTTACTGTTACTGCT
GATGTTGAAGTTCCTGAAGGTCCTAGAACTTGGTGTATGACTAGAAAACCTGGTGCTTAA
TGTAATTTAAGTTGTTATATAAAGATCTGAACTGCAGGTCGACTCTAGAG

### Nucleic sequence of BseRI double sites – Sp sgRNA

ATCTATGTCTCCTCTTAGAGGAGTTAGCTTAgttttagagctagaaatagcaagttaaaa
taaggctagtccgttatcaacttgaaaaagtggcaccgagtcggtgcttttttttacg

### Nucleic sequence of Spiralin promoter

AGAATTAAAAGTTAGTGAACAAGAAAACAGTGAAGCACCAGTTTCTGAACCAAAAGAAGA
CGAAAAAACAAAAAAAGATTAAGCAATTTATTTGGAAAATCTTTTTTTGTTTTTTTAAGA
AATATTTATTGTTTTTTTTAAAAAATTATGTACAATTGCTACTATAAGGGAAAGAAAAAA
AGAAAGATATAAATTGTATAAAGTAGGGTTAGAAGCAATTAATAATTATTATTAATGTTA
TTTTTCTCTTATATATTCAATGTAATTTTAATTACATTTGCTTTTAATAAAAACACTACT
TAATAGAGAAAGGAAATATAA

### Nucleic sequence of Fibril terminator

GATCTAAATTAAAGTTGGTTCATTCAAAGTTACAATTCGTTTAAAAAGTGAATAATTAAT
TATTAATTAATTTATTAAAAAACAACCAAAAGGTTGTTTTTTATTTTTTTAAA

### Nucleic sequence of inducible prom pXyl-TetO

GTCCGTAATACGACTCACTTAAGGCCTTGACTAGAGGGTACCAATTCCAGAGTCTCCATA
TATGAATAATGGATTTCATTTTTTTCCAAGATCTAAGAAATGAAAAAACTTTTATCGATC
AAAAACTAAAAAAAATATTGACACTCTATCATTGATAGAGTATAATTAACGGGATCCTCT
ATCATTGATAGAGGGATCCCGCCAAGCTTGGGATCCCCAGCTTGTTGATACACTAATGCT
TTTATATAGGGAAAAGGTGGTGAACTACT

### pTi4.0_SpdCas9_rAPOBEC1

TACATTTCCAACATTGAAGCCATGAGGGCTTTTTCGTTTCGTTGATAACGTTCAAACACT
GTGGGTGAAAAATGGCCATCACGTGTTCTGGGTACTTTTAATTCTAGCGTGCCTACACGT
GTCGTAAAGCTGCGCTCATAATAGCCATTTCGTTGACTTTGTCGGTTTTCTGTTCGTTCA
TATTCTTTTGCTTGAATATATTCTGTTCGTTGATTTTCCATTAGTTGATTAAATACCGTT
GTTAAAATATTTTTAGAAACGTCATCCTTTACAGAATATTCAATAATGCTTTGAATCTCT
TCGCTTTTCAGTGTAAAATGTACTTGGGTCATGTAAAAGTCCTCCTGGGTATGTTTTTGT
CGTTAAAAACATTGTACCGTAAAAGGACTGTTATATGGCCTTTTTACTTTTACACGTCGA
CGGTATCGATAAGCTTGATAAAGTCCGTATAATTGTGTAAAAGAATTCGCTTGCATGCCT
GCAGAATTAAAAGTTAGTGAACAAGAAAACAGTGAAGCACCAGTTTCTGAACCAAAAGAA
GACGAAAAAACAAAAAAAGATTAAGCAATTTATTTGGAAAATCTTTTTTTGTTTTTTTAA
GAAATATTTATTGTTTTTTTTAAAAAATTATTGTACAGTTGCTACTATAAGGGAAAGAAA
AAAAGAAAGATATAAATTGTATAAAGTAGGGTTAGAAGCAATTAATAATTATTATTAATG
TTATTTTTCTCTTATATATTCAATGTAATTTTAATTACATTTGCTTTTAATAAAAACACT
ACTTAATAGAGAAAGGAAATATAAGATCTCATATGTCTAGATTAGATAAAAGTAAAGTGA
TTAACAGCGCATTAGAGCTGCTTAATGAGGTCGGAATCGAAGGTTTAACAACCCGTAAAC
TCGCCCAGAAGCTAGGTGTAGAGCAGCCTACATTGTATTGGCATGTAAAAAATAAGCGGG
CTTTGCTCGACGCCTTAGCCATTGAGATGTTAGATAGGCACCATACTCACTTTTGCCCTT
TAGAAGGGGAAAGCTGGCAAGATTTTTTACGTAATAACGCTAAAAGTTTTAGATGTGCTT
TACTAAGTCATCGCGATGGAGCAAAAGTACATTTAGGTACACGGCCTACAGAAAAACAGT
ATGAAACTCTCGAAAATCAATTAGCCTTTTTATGCCAACAAGGTTTTTCACTAGAGAATG
CATTATATGCACTCAGCGCTGTGGGGCATTTTACTTTAGGTTGCGTATTGGAAGATCAAG
AGCATCAAGTCGCTAAAGAAGAAAGGGAAACACCTACTACTGATAGTATGCCGCCATTAT
TACGACAAGCTATCGAATTATTTGATCACCAAGGTGCAGAGCCAGCCTTCTTATTCGGCC
TTGAATTGATCATATGCGGATTAGAAAAACAACTTAAATGTGAAAGTGGGTCTTAAGGAT
CTGAACTGCAGGTCGATggatccCCGCCTCCAGCAGAATTAAAAGTTAGTGAACAAGAAA
ACAGTGAAGCACCAGTTTCTGAACCAAAAGAAGACGAAAAAACAAAAAAAGATTAAGCAA
TTTATTTGGAAAATCTTTTTTTGTTTTTTTAAGAAATATTTATTGTTTTTTTTAAAAAAT
TATGTACAATTGCTACTATAAGGGAAAGAAAAAAAGAAAGATATAAATTGTATAAAGTAG
GGTTAGAAGCAATTAATAATTATTATTAATGTTATTTTTCTCTTATATATTCAATGTAAT
TTTAATTACATTTGCTTTTAATAAAAACACTACTTAATAGAGAAAGGAAATATAACCTAG
GGTCTCCTCTTAGAGGAGTTAGCTTAGTTTTAGAGCTAGAAATAGCAAGTTAAAATAAGG
CTAGTCCGTTATCAACTTGAAAAAGTGGCACCGAGTCGGTGCTTTTTTTACGGATCTAAA
TTAAAGTTGGTTCATTCAAAGTTACAATTCGTTTAAAAAGTGAATAATTAATTATTAATT
AATTTATTAAAAAACAACCAAAAGGTTGTTTTTTATTTTTTTAAAGGTCGGTCCGTAATAC
GACTCACTTAAGGCCTTGACTAGAGGGTACCAATTCCAGAGTCTCCATATATGAATAATG
GATTTCATTTTTTTCCAAGATCTAAGAAATGAAAAAACTTTTATCGATCAAAAACTAAAA
AAAATATTGACACTCTATCATTGATAGAGTATAATTAACGGGATCCTCTATCATTGATAG
AGGGATCCCGCCAAGCTTGGGATCCCCAGCTTGTTGATACACTAATGCTTTTATATAGGG
AAAAGGTGGTGAACTACTATGAGTAGTGAAACTGGACCAGTTGCTGTTGACCCCACTCTC
AGAAGAAGAATAGAACCACATGAATTTGAAGTATTCTTCGATCCAAGAGAATTAAGAAAA
GAAACTTGTTTATTATATGAAATTAATTGGGGAGGAAGACATAGTATTTGGAGACATACT
AGTCAAAATACTAATAAACATGTTGAAGTTAATTTTATTGAGAAGTTTACTACTGAAAGA
TATTTCTGTCCAAATACTAGATGTAGTATTACTTGGTTCTTAAGTTGGAGTCCATGTGGA
GAATGTAGTAGAGCTATAACTGAATTCCTGAGTAGATATCCACATGTTACTTTATTTATT
TATATTGCTAGATTATATCATCATGCTGATCCAAGAAATAGACAAGGATTAAGAGATTTA

ATTAGTAGTGGAGTTACTATTCAAATTATGACTGAACAAGAAAGTGGATATTGTTGGAGA
AATTTTGTTAATTATAGTCCAAGTAATGAAGCTCATTGGCCAAGATATCCACATTTATGG
GTTAGATTATATGTTTTAGAATTATATTGTATTATTTTAGGATTACCACCATGTTTAAAT
ATTTTAAGAAGAAAACAACCACAATTAACATTCTTCACTATTGCTTTACAAAGTTGTCAT
TATCAAAGATTACCACCACATATTTTATGGGCTACTGGATTAAAAAGTGGAAGTGAAACT
CCAGGAACTAGTGAAAGTGCTACTCCAGAAAGTGACAAAAAATATTCTATTGGATTAGCT
ATTGGAACTAATTCAGTAGGTTGAGCTGTTATTACAGATGAATATAAGGTACCATCAAAG
AAATTTAAGGTTTTAGGTAATACTGATAGACATTCAATTAAAAAAAATTTAATCGGAGCA
TTACTTTTTGATTCTGGAGAAACAGCAGAAGCTACCAGATTAAAAAGAACGGCTCGTAGA
CGATATACTAGACGTAAAAACAGAATCTGTTATCTTCAAGAAATTTTTAGTAATGAAATG
GCTAAAGTTGATGACTCTTTTTTTCACAGATTGGAAGAGTCTTTCTTAGTAGAAGAAGAT
AAAAAGCATGAAAGACACCCCATCTTTGGTAATATTGTAGATGAAGTCGCCTATCATGAA
AAATATCCTACAATTTATCATTTAAGAAAAAAATTAGTAGATAGCACAGATAAAGCTGAT
TTAAGATTAATTTATTTAGCACTAGCACATATGATCAAATTTAGAGGTCACTTCTTAATT
GAAGGTGATTTAAACCCTGATAATAGTGACGTTGATAAATTATTTATTCAATTAGTACAA
ACGTATAATCAACTTTTCGAAGAAAACCCAATTAATGCTAGTGGGGTTGATGCAAAAGCT
ATCCTTTCGGCTCGTCTTTCAAAATCTAGGAGACTTGAAAACTTAATTGCACAATTACCG
GGAGAGAAAAAAAACGGTTTATTTGGTAACTTAATCGCGTTATCTTTAGGTTTAACCCCG
AATTTTAAGAGTAATTTCGATCTAGCTGAAGATGCTAAACTACAATTATCTAAAGATACT
TATGACGATGACTTAGATAATTTATTAGCTCAGATTGGTGATCAATATGCAGACTTATTT
TTAGCAGCAAAAAACTTAAGCGACGCAATCTTATTGAGTGATATATTGAGAGTTAACACA
GAAATCACTAAAGCACCATTAAGTGCAAGTATGATTAAACGTTATGATGAACACCACCAA
GATTTAACACTATTAAAAGCATTAGTTAGACAACAATTACCTGAAAAGTATAAAGAAATT
TTCTTCGATCAAAGCAAAAATGGTTATGCTGGTTATATTGATGGTGGAGCTTCACAAGAA
GAATTTTATAAGTTCATTAAGCCTATCCTAGAAAAAATGGATGGAACAGAAGAACTATTA
GTCAAGTTAAATCGTGAAGATTTACTACGCAAACAAAGAACTTTTGATAATGGTAGCATT
CCTCATCAAATTCACTTAGGAGAACTACACGCTATCCTAAGAAGACAAGAAGATTTTTAT
CCTTTTTTAAAAGATAATAGAGAAAAAATTGAAAAAATCTTAACATTTAGAATCCCTTAC
TATGTAGGTCCGTTAGCTAGAGGAAATAGTAGATTTGCATGAATGACTCGAAAATCAGAA
GAGACTATCACACCATGAAATTTTGAGGAAGTTGTGGATAAAGGTGCATCTGCGCAATCT
TTTATTGAGCGAATGACTAATTTCGATAAGAACTTACCTAATGAAAAAGTATTACCTAAG
CACTCATTATTATATGAATACTTTACTGTTTATAACGAACTTACTAAAGTAAAATATGTT
ACCGAAGGAATGAGAAAACCAGCGTTCCTAAGTGGAGAACAAAAGAAGGCTATTGTTGAT
TTATTATTTAAGACAAATAGAAAAGTAACTGTAAAACAACTAAAAGAAGATTATTTTAAA
AAAATTGAATGTTTTGATTCAGTCGAAATTTCTGGAGTTGAAGACCGTTTCAACGCAAGT
TTAGGCACTTACCACGATCTACTAAAAATTATTAAAGATAAAGATTTTCTTGATAACGAA
GAAAATGAAGACATTCTAGAAGATATTGTCCTAACTTTAACTTTATTCGAAGACAGAGAA
ATGATTGAAGAAAGATTAAAAACTTACGCTCACTTATTTGATGATAAAGTTATGAAGCAG
TTGAAGCGCCGACGATATACCGGTTGAGGTAGACTCTCAAGAAAGCTAATCAATGGTATT
AGAGACAAACAATCAGGTAAAACAATTTTAGATTTTTTAAAAAGCGACGGATTTGCTAAT
AGAAACTTCATGCAATTGATCCACGATGATTCATTAACTTTTAAAGAAGATATTCAAAAG
GCACAAGTCTCAGGTCAAGGTGATAGCTTACACGAACATATCGCTAATCTAGCTGGTTCA
CCTGCAATCAAAAAGGGAATTTTACAGACAGTGAAAGTTGTTGATGAACTAGTAAAGGTA
ATGGGACGCCACAAACCAGAGAACATCGTGATTGAAATGGCTAGAGAAAACCAAACAACA
CAAAAAGGCCAAAAAAACAGTAGAGAAAGAATGAAGAGAATCGAAGAAGGTATCAAAGAG
TTAGGGTCTCAAATCTTAAAGGAACATCCTGTTGAAAACACTCAATTACAAAATGAAAAA
TTATACCTATATTACTTACAAAATGGTAGAGATATGTATGTTGATCAGGAATTAGATATT
AACCGTTTATCAGATTACGACGTGGATGCTATTGTACCTCAATCATTTCTAAAAGATGAT
TCTATCGACAATAAGGTTTTAACTAGATCCGATAAAAATCGAGGAAAATCTGACAATGTA
CCTAGTGAAGAAGTTGTAAAAAAGATGAAAAATTACTGACGACAACTTTTAAACGCAAAA
TTAATTACACAAAGAAAATTTGATAACCTAACTAAAGCAGAGCGTGGAGGTCTGTCTGAA
CTTGATAAGGCTGGATTTATTAAACGACAACTAGTTGAAACACGTCAAATCACCAAACAT
GTTGCACAAATTTTAGATTCTCGTATGAATACAAAGTACGACGAAAACGATAAATTAATC
AGAGAAGTTAAAGTTATTACTCTAAAATCAAAATTAGTGAGTGATTTTCGCAAAGATTTC
CAATTTTACAAGGTTAGAGAGATCAATAATTATCATCACGCACATGATGCTTATTTAAAT
GCTGTGGTTGGGACTGCTTTAATCAAAAAGTATCCTAAATTAGAAAGCGAATTTGTATAC
GGTGATTATAAGGTTTATGATGTTCGTAAAATGATTGCTAAAAGTGAACAAGAAATTGGA
AAGGCTACTGCTAAATATTTTTTTTACTCAAATATTATGAATTTTTTCAAAACTGAGATC
ACATTAGCAAATGGTGAAATCCGTAAAAGACCTTTAATTGAAACTAACGGGGAAACTGGG
GAAATTGTGTGAGATAAAGGTCGTGATTTTGCAACAGTAAGAAAAGTATTATCAATGCCA
CAAGTTAATATCGTTAAAAAAACAGAGGTGCAAACTGGAGGTTTCTCTAAAGAATCGATC
TTACCTAAAAGAAACAGTGATAAATTAATTGCGAGAAAAAAAGATTGAGATCCAAAAAAA
TATGGTGGTTTTGATTCGCCGACTGTTGCATACTCTGTATTAGTGGTTGCTAAAGTTGAA
AAAGGTAAGAGTAAAAAATTAAAATCTGTTAAAGAATTATTAGGCATTACGATTATGGAG
AGATCATCTTTTGAAAAAAATCCAATCGATTTTTTGGAAGCTAAAGGGTACAAAGAAGTA
AAAAAAGATCTTATAATTAAATTACCTAAATATTCTTTATTTGAACTTGAAAACGGTCGT
AAAAGAATGTTAGCTTCTGCAGGAGAACTACAAAAAGGTAATGAATTAGCTCTACCAAGT
AAATATGTAAATTTTTTATATTTAGCGAGTCACTATGAAAAATTAAAAGGATCACCTGAA
GATAATGAACAAAAACAATTATTTGTTGAACAACACAAACATTATTTAGATGAAATTATT
GAACAAATTTCAGAATTTAGCAAAAGAGTTATACTTGCCGATGCAAATTTAGATAAAGTG
TTAAGTGCGTACAACAAGCACAGAGATAAACCAATCCGCGAACAAGCAGAAAATATTATC
CACTTATTCACTCTTACTAACTTAGGTGCTCCAGCAGCTTTTAAATATTTTGATACAACT
ATCGATAGAAAGAGATATACATCAACAAAGGAAGTTTTAGACGCGACTTTAATTCATCAA
AGTATTACTGGACTTTATGAGACTCGCATCGATCTATCACAATTGGGTGGAGACAGCGGT
GGAAGTACTAATTTAAGTGATATTATTGAGAAAGAAACTGGAAAACAATTAGTTATTCAA
GAAAGTATTTTAATGTTACCAGAAGAAGTTGAAGAAGTTATTGGAAATAAACCAGAAAGT
GATATTTTAGTTCATACTGCTTATGATGAAAGTACTGATGAAAATGTTATGTTATTAACT

AGTGATGCTCCAGAATATAAACCATGGGCTTTAGTTATTCAAGATAGTAATGGAGAAAAT
AAAATTAAAATGTTATAAGATCTAAATTAAAGTTGGTTCATTCAAAGTTACAATTCGTTT
AAAAAGTGAATAATTAATTATTAATTAATTTATTAAAAAACAACCAAAAGGTTGTTTTTT
ATTTTTTAAAGAAGGATGGTCTCCAGACACTTATAAGCCTCTCTACTGCAATTTATCTCA
ACTTTGATATAATTAAAGACATACGAAAGGATTTTAATATGACTGAATATAAACCTACTG
TTAGATTAGCTACTAGAGATGATGTTCCTAGAGCTGTTAGAACTTTAGCTGCTGCTTTTG
CTGATTATCCTGCTACTAGACATACTGTTGATCCTGATAGACATATTGAAAGAGTTACTG
AATTACAAGAATTATTCTTAACTAGAGTTGGTTTAGATATTGGTAAAGTTTGGGTTGCTG
ATGATGGTGCTGCTGTTGCTGTTTGGACTACTCCTGAAAGTGTTGAAGCTGGTGCTGTAT
TTGCTGAAATTGGTCCTCGTATGGCTGAATTAAGTGGTAGTAGATTAGCTGCTCAACAAC
AAATGGAAGGTTTATTAGCTCCACATAGACCTAAAGAACCTGCTTGGTTCTTAGCTACTG
TTGGTGTTAGTCCTGATCATCAAGGTAAAGGTTTAGGTAGTGCTGTTGTTTTACCTGGTG
TTGAAGCTGCTGAAAGAGCTGGTGTTCCTGCTTTCTTAGAAACTAGTGCTCCTAGAAATT
TACCTTTCTATGAAAGATTAGGTTTTACTGTTACTGCTGATGTTGAAGTTCCTGAAGGTC
CTAGAACTTGGTGTATGACTAGAAAACCTGGTGCTTAATGTAATTTAAGTTGTTATATAA
AGATCTGAACTGCAGGTCGACTCTAGAGCTCGAGACTAGTACTAGTTTTTACACAGGAGT
CTGGACTTGACTGAGCTCCAGCTTTTGTTCCCTTTAGTGAGGGTTAATTTCGAGCTTGGC
GTAATCATGGTCATAGCTGTTTCCTGTGTGAAATTGTTATCCGCTCACAATTCCACACAA
CATACGAGCCGGAAGCATAAAGTGTAAAGCCTGGGGTGCCTAATGAGTGAGCTAACTCAC
ATTAATTGCGTTGCGCTCACTGCCCGCTTTCCAGTCGGGAAACCTGTCGTGCCAGCTGCA
TTAATGAATCGGCCAACGCGCGGGGAGAGGCGGTTTGCGTATTGGGCGCTCTTCCGCTTC
CTCGCTCACTGACTCGCTGCGCTCGGTCGTTCGGCTGCGGCGAGCGGTATCAGCTCACTC
AAAGGCGGTAATACGGTTATCCACAGAATCAGGGGATAACGCAGGAAAGAACATGTGAGC
AAAAGGCCAGCAAAAGGCCAGGAACCGTAAAAAGGCCGCGTTGCTGGCGTTTTTCCATAG
GCTCCGCCCCCCTGACGAGCATCACAAAAATCGACGCTCAAGTCAGAGGTGGCGAAACCC
GACAGGACTATAAAGATACCAGGCGTTTCCCCCTGGAAGCTCCCTCGTGCGCTCTCCTGT
TCCGACCCTGCCGCTTACCGGATACCTGTCCGCCTTTCTCCCTTCGGGAAGCGTGGCGCT
TTCTCATAGCTCACGCTGTAGGTATCTCAGTTCGGTGTAGGTCGTTCGCTCCAAGCTGGG
CTGTGTGCACGAACCCCCCGTTCAGCCCGACCGCTGCGCCTTATCCGGTAACTATCGTCT
TGAGTCCAACCCGGTAAGACACGACTTATCGCCACTGGCAGCAGCCACTGGTAACAGGAT
TAGCAGAGCGAGGTATGTAGGCGGTGCTACAGAGTTCTTGAAGTGGTGGCCTAACTACGG
CTACACTAGAAGAACAGTATTTGGTATCTGCGCTCTGCTGAAGCCAGTTACCTTCGGAAA
AAGAGTTGGTAGCTCTTGATCCGGCAAACAAACCACCGCTGGTAGCGGTGGTTTTTTTGT
TTGCAAGCAGCAGATTACGCGCAGAAAAAAAGGATCTCAAGAAGATCCTTTGATCTTTTC
TACGGGGTCTGACGCTCAGTGGAACGAAAACTCACGTTAAGGGATTTTGGTCATGAGATT
ATCAAAAAGGATCTTCACCTAGATCCTTTTAAATTAAAAATGAAGTTTTAAATCAATCTA
AAGTATATATGAGTAAACTTGGTCTGACAGTTACCAATGCTTAATCAGTGAGGCACCTAT
CTCAGCGATCTGTCTATTTCGTTCATCCATAGTTGCCTGACTCCCCGTCGTGTAGATAAC
TACGATACGGGAGGGCTTACCATCTGGCCCCAGTGCTGCAATGATACCGCGAGACCCACG
CTCACCGGCTCCAGATTTATCAGCAATAAACCAGCCAGCCGGAAGGGCCGAGCGCAGAAG
TGGTCCTGCAACTTTATCCGCCTCCATCCAGTCTATTAATTGTTGCCGGGAAGCTAGAGT
AAGTAGTTCGCCAGTTAATAGTTTGCGCAACGTTGTTGCCATTGCTACAGGCATCGTGGT
GTCACGCTCGTCGTTTGGTATGGCTTCATTCAGCTCCGGTTCCCAACGATCAAGGCGAGT
TACATGATCCCCCATGTTGTGCAAAAAAGCGGTTAGCTCCTTCGGTCCTCCGATCGTTGT
CAGAAGTAAGTTGGCCGCAGTGTTATCACTCATGGTTATGGCAGCACTGCATAATTCTCT
TACTGTCATGCCATCCGTAAGATGCTTTTCTGTGACTGGTGAGTACTCAACCAAGTCATT
CTGAGAATAGTGTATGCGGCGACCGAGTTGCTCTTGCCCGGCGTCAATACGGGATAATAC
CGCGCCACATAGCAGAACTTTAAAAGTGCTCATCATTGGAAAACGTTCTTCGGGGCGAAA
ACTCTCAAGGATCTTACCGCTGTTGAGATCCAGTTCGATGTAACCCACTCGTGCACCCAA
CTGATCTTCAGCATCTTTTACTTTCACCAGCGTTTCTGGGTGAGCAAAAACAGGAAGGCA
AAATGCCGCAAAAAAGGGAATAAGGGCGACACGGAAATGTTGAATACTCATACTCTTCCT
TTTTCAATATTATTGAAGCATTTATCAGGGTTATTGTCTCATGAGCGGATACATATTTGA
ATGTATTTAGAAAAATAAACAAATAGGGGTTCCGCGCACATTTCCCCGAAAAGTGCCACC
TAAATTGTAAGCGTTAATATTTTGTTAAAATTCGCGTTAAATTTTTGTTAAATCAGCTCA
TTTTTTAACCAATAGGCCGAAATCGGCAAAATCCCTTATAAATCAAAAGAATAGACCGAG
ATAGGGTTGAGTGTTGTTCCAGTTTGGAACAAGAGTCCACTATTAAAGAACGTGGACTCC
AACGTCAAAGGGCGAAAAACCGTCTATCAGGGCGATGGCCCACTACGTGAACCATCACCC
TAATCAAGTTTTTTGGGGTCGAGGTGCCGTAAAGCACTAAATCGGAACCCTAAAGGGAGC
CCCCGATTTAGAGCTTGACGGGGAAAGCCGGCGAACGTGGCGAGAAAGGAAGGGAAGAAA
GCGAAAGGAGCGGGCGCTAGGGCGCTGGCAAGTGTAGCGGTCACGCTGCGCGTAACCACC
ACACCCGCCGCGCTTAATGCGCCGCTACAGGGCGCGTCCCATTCGCCATTCAGGCTGCGC
AACTGTTGGGAAGGGCGATCGGTGCGGGCCTCTTCGCTATTACGCCAGCTGGCGAAAGGG
GGATGTGCTGCAAGGCGATTAAGTTGGGTAACGCCAGGGTTTTCCCAGTCACGACGTTGT
AAAACGACGGCCAGTGAATTGTAATACGACTCACTATAGGGCGAATTGGGTACCCCATTT
CTACTTATCAAAATTGATGTATTTTCTTGAAGAATAAATCCATTCATCATGTAGGTCCAT
AAGAACGGCTCCAATTAAGCGATTGGCTGATGTTTGATTGGGGAAGATGCGAATAATCTT
TTCTCTTCTGCGTACTTCTTGATTCAGTCGTTCAATTAGATTGGTACTCTTTAGTCGATT
GTGGGAATTTCCTTGTACGGTATATTGAAAGGCGTCTTCGAATCCATCATCCAATGATGC
GCAAGCTTTTGAATATTTTGGTTGATCGATATAATCATGAATCAATCGATTTTTAGCCTC
ACGCGCTAAGTTAATATCTGTGAACTTAAAAATTCCTTTAACAGCTTCTCTGAAAGATTT
TGAATTTTTTTTAGGAATGGTGGTAAAGATATTTCTTAGGAAGTGAACTTGGCATCTTTG
CCAACTTACGTTGGTGAAGGATTTTCTAATGGCAGAGACTAATCCTTTGTGCGCATCAGA
AATAACGAGTTCCGTACCTTGTAAACCGCGTTCTTTTAGGTATTCAAAAAATGTTGTCCA
GGTCTCTTCGCTTTCGCCACTTTGAATCATGAAGCCGATAATTTCACGGTCGCCATCTTT
GGTTATTCCAATCGCTATATGACAGCTTTTTGAGAGTACTCGATTTTCTTCTCGTACTTT
TATATAGAGTACATCGGTCATTAAGTAAGGATAATTTTTTTCTGATAATAAACGATTCTG
CCACTCGTTAACCATAGGTTCTAGCTGTTCTGTTAAGCTAGAAACGAAGGACTTAGAGAC

GGATTTACCACAAAGTTCTTCCACAATTTTTGATACTTTACGAGTTGAAACGCCTGATAC
A

### pTi4.0_SpdCas9_pmcDA1

TGATTCATTAACTTTTAAAGAAGATATTCAAAAGGCACAAGTCTCAGGTCAAGGTGATAG
CTTACACGAACATATCGCTAATCTAGCTGGTTCACCTGCAATCAAAAAGGGAATTTTACA
GACAGTGAAAGTTGTTGATGAACTAGTAAAGGTAATGGGACGCCACAAACCAGAGAACAT
CGTGATTGAAATGGCTAGAGAAAACCAAACAACACAAAAAGGCCAAAAAAACAGTAGAGA
AAGAATGAAGAGAATCGAAGAAGGTATCAAAGAGTTAGGGTCTCAAATCTTAAAGGAACA
TCCTGTTGAAAACACTCAATTACAAAATGAAAAATTATACCTATATTACTTACAAAATGG
TAGAGATATGTATGTTGATCAGGAATTAGATATTAACCGTTTATCAGATTACGACGTGGA
TGCTATTGTACCTCAATCATTTCTAAAAGATGATTCTATCGACAATAAGGTTTTAACTAG
ATCCGATAAAAATCGAGGAAAATCTGACAATGTACCTAGTGAAGAAGTTGTAAAAAAGAT
GAAAAATTACTGACGACAACTTTTAAACGCAAAATTAATTACACAAAGAAAATTTGATAA
CCTAACTAAAGCAGAGCGTGGAGGTCTGTCTGAACTTGATAAGGCTGGATTTATTAAACG
ACAACTAGTTGAAACACGTCAAATCACCAAACATGTTGCACAAATTTTAGATTCTCGTAT
GAATACAAAGTACGACGAAAACGATAAATTAATCAGAGAAGTTAAAGTTATTACTCTAAA
ATCAAAATTAGTGAGTGATTTTCGCAAAGATTTCCAATTTTACAAGGTTAGAGAGATCAA
TAATTATCATCACGCACATGATGCTTATTTAAATGCTGTGGTTGGGACTGCTTTAATCAA
AAAGTATCCTAAATTAGAAAGCGAATTTGTATACGGTGATTATAAGGTTTATGATGTTCG
TAAAATGATTGCTAAAAGTGAACAAGAAATTGGAAAGGCTACTGCTAAATATTTTTTTTA
CTCAAATATTATGAATTTTTTCAAAACTGAGATCACATTAGCAAATGGTGAAATCCGTAA
AAGACCTTTAATTGAAACTAACGGGGAAACTGGGGAAATTGTGTGAGATAAAGGTCGTGA
TTTTGCAACAGTAAGAAAAGTATTATCAATGCCACAAGTTAATATCGTTAAAAAAACAGA
GGTGCAAACTGGAGGTTTCTCTAAAGAATCGATCTTACCTAAAAGAAACAGTGATAAATT
AATTGCGAGAAAAAAAGATTGAGATCCAAAAAAATATGGTGGTTTTGATTCGCCGACTGT
TGCATACTCTGTATTAGTGGTTGCTAAAGTTGAAAAAGGTAAGAGTAAAAAATTAAAATC
TGTTAAAGAATTATTAGGCATTACGATTATGGAGAGATCATCTTTTGAAAAAAATCCAAT
CGATTTTTTGGAAGCTAAAGGGTACAAAGAAGTAAAAAAAGATCTTATAATTAAATTACC
TAAATATTCTTTATTTGAACTTGAAAACGGTCGTAAAAGAATGTTAGCTTCTGCAGGAGA
ACTACAAAAAGGTAATGAATTAGCTCTACCAAGTAAATATGTAAATTTTTTATATTTAGC
GAGTCACTATGAAAAATTAAAAGGATCACCTGAAGATAATGAACAAAAACAATTATTTGT
TGAACAACACAAACATTATTTAGATGAAATTATTGAACAAATTTCAGAATTTAGCAAAAG
AGTTATACTTGCCGATGCAAATTTAGATAAAGTGTTAAGTGCGTACAACAAGCACAGAGA
TAAACCAATCCGCGAACAAGCAGAAAATATTATCCACTTATTCACTCTTACTAACTTAGG
TGCTCCAGCAGCTTTTAAATATTTTGATACAACTATCGATAGAAAGAGATATACATCAAC
AAAGGAAGTTTTAGACGCGACTTTAATTCATCAAAGTATTACTGGACTTTATGAGACTCG
CATCGATCTATCACAATTGGGTGGAGACGGAGGTGGTGGAACTGGTGGTGGTGGTAGTGC
TGAATATGTTAGAGCTTTATTTGATTTTAATGGTAATGATGAAGAAGATTTACCATTTAA
GAAAGGTGATATTTTAAGAATTAGAGATAAACCAGAAGAACAATGGTGGAATGCTGAAGA
TAGTGAAGGTAAAAGAGGTATGATTTTAGTTCCATATGTTGAGAAATATAGTGGTACTGA
TGCTGAATATGTTAGAATTCATGAGAAATTAGATATTTATACTTTTAAGAAACAATTCTT
TAATAATAAGAAGAGTGTTAGTCATCGTTGCTATGTGCTCTTTGAATTAAAGAGAAGAGG
TGAAAGAAGAGCTTGCTTTTGAGGATATGCTGTTAATAAACCACAAAGTGGAACTGAAAG
AGGAATTCATGCTGAAATATTTAGTATTAGAAAAGTTGAAGAATATTTAAGAGATAATCC
AGGACAATTTACTATTAATTGGTATAGTAGTTGGAGTCCATGTGCTGATTGTGCTGAGAA
GATTTTAGAATGGTATAATCAAGAATTAAGAGGAAATGGACATACTTTGAAAATTTGGGC
TTGTAAATTATATTATGAGAAGAATGCTAGAAATCAAATTGGATTATGGAATTTAAGAGA
TAATGGAGTTGGATTAAATGTTATGGTTAGTGAACATTATCAATGTTGTAGAAAGATATT
TATTCAAAGTAGTCATAATCAATTAAATGAAAATAGATGGTTAGAGAAAACTTTAAAGAG
AGCTGAGAAAAGAAGAAGTGAATTAAGTATTATGATTCAAGTTAAAATTTTACATACTAC
TAAAAGTCCAGCTGTTAGTGGAGGAAGTACTAATTTAAGTGATATTATTGAGAAAGAAAC
TGGAAAACAATTAGTTATTCAAGAAAGTATTTTAATGTTACCAGAAGAAGTTGAAGAAGT
TATTGGAAATAAACCAGAAAGTGATATTTTAGTTCATACTGCTTATGATGAAAGTACTGA
TGAAAATGTTATGTTATTAACTAGTGATGCTCCAGAATATAAACCATGGGCTTTAGTTAT
TCAAGATAGTAATGGAGAAAATAAAATTAAAATGTTATAAGATCTAAATTAAAGTTGGTT
CATTCAAAGTTACAATTCGTTTAAAAAGTGAATAATTAATTATTAATTAATTTATTAAAA
AACAACCAAAAGGTTGTTTTTTATTTTTTAAAGAAGGATGGTCTCCAGACACTTATAAGC
CTCTCTACTGCAATTTATCTCAACTTTGATATAATTAAAGACATACGAAAGGATTTTAAT
ATGACTGAATATAAACCTACTGTTAGATTAGCTACTAGAGATGATGTTCCTAGAGCTGTT
AGAACTTTAGCTGCTGCTTTTGCTGATTATCCTGCTACTAGACATACTGTTGATCCTGAT
AGACATATTGAAAGAGTTACTGAATTACAAGAATTATTCTTAACTAGAGTTGGTTTAGAT
ATTGGTAAAGTTTGGGTTGCTGATGATGGTGCTGCTGTTGCTGTTTGGACTACTCCTGAA
AGTGTTGAAGCTGGTGCTGTATTTGCTGAAATTGGTCCTAGAATGGCTGAATTAAGTGGT
AGTAGATTAGCTGCTCAACAACAAATGGAAGGTTTATTAGCTCCACATAGACCTAAAGAA
CCTGCTTGGTTCTTAGCTACTGTTGGTGTTAGTCCTGATCATCAAGGTAAAGGTTTAGGT
AGTGCTGTTGTTTTACCTGGTGTTGAAGCTGCTGAAAGAGCTGGTGTTCCTGCTTTCTTA
GAAACTAGTGCTCCTAGAAATTTACCTTTCTATGAAAGATTAGGTTTTACTGTTACTGCT
GATGTTGAAGTTCCTGAAGGTCCTAGAACTTGGTGTATGACTAGAAAACCTGGTGCTTAA
TGTAATTTAAGTTGTTATATAAAGATCTGAACTGCAGGTCGACTCTAGAGACTAGTACTA
GTTTTTACACAGGAGTCTGGACTTGACTGAGCTCCAGCTTTTGTTCCCTTTAGTGAGGGT
TAATTTCGAGCTTGGCGTAATCATGGTCATAGCTGTTTCCTGTGTGAAATTGTTATCCGC
TCACAATTCCACACAACATACGAGCCGGAAGCATAAAGTGTAAAGCCTGGGGTGCCTAAT
GAGTGAGCTAACTCACATTAATTGCGTTGCGCTCACTGCCCGCTTTCCAGTCGGGAAACC
TGTCGTGCCAGCTGCATTAATGAATCGGCCAACGCGCGGGGAGAGGCGGTTTGCGTATTG
GGCGCTCTTCCGCTTCCTCGCTCACTGACTCGCTGCGCTCGGTCGTTCGGCTGCGGCGAG
CGGTATCAGCTCACTCAAAGGCGGTAATACGGTTATCCACAGAATCAGGGGATAACGCAG

GAAAGAACATGTGAGCAAAAGGCCAGCAAAAGGCCAGGAACCGTAAAAAGGCCGCGTTGC
TGGCGTTTTTCCATAGGCTCCGCCCCCCTGACGAGCATCACAAAAATCGACGCTCAAGTC
AGAGGTGGCGAAACCCGACAGGACTATAAAGATACCAGGCGTTTCCCCCTGGAAGCTCCC
TCGTGCGCTCTCCTGTTCCGACCCTGCCGCTTACCGGATACCTGTCCGCCTTTCTCCCTT
CGGGAAGCGTGGCGCTTTCTCATAGCTCACGCTGTAGGTATCTCAGTTCGGTGTAGGTCG
TTCGCTCCAAGCTGGGCTGTGTGCACGAACCCCCCGTTCAGCCCGACCGCTGCGCCTTAT
CCGGTAACTATCGTCTTGAGTCCAACCCGGTAAGACACGACTTATCGCCACTGGCAGCAG
CCACTGGTAACAGGATTAGCAGAGCGAGGTATGTAGGCGGTGCTACAGAGTTCTTGAAGT
GGTGGCCTAACTACGGCTACACTAGAAGAACAGTATTTGGTATCTGCGCTCTGCTGAAGC
CAGTTACCTTCGGAAAAAGAGTTGGTAGCTCTTGATCCGGCAAACAAACCACCGCTGGTA
GCGGTGGTTTTTTTGTTTGCAAGCAGCAGATTACGCGCAGAAAAAAAGGATCTCAAGAAG
ATCCTTTGATCTTTTCTACGGGGTCTGACGCTCAGTGGAACGAAAACTCACGTTAAGGGA
TTTTGGTCATGAGATTATCAAAAAGGATCTTCACCTAGATCCTTTTAAATTAAAAATGAA
GTTTTAAATCAATCTAAAGTATATATGAGTAAACTTGGTCTGACAGTTACCAATGCTTAA
TCAGTGAGGCACCTATCTCAGCGATCTGTCTATTTCGTTCATCCATAGTTGCCTGACTCC
CCGTCGTGTAGATAACTACGATACGGGAGGGCTTACCATCTGGCCCCAGTGCTGCAATGA
TACCGCGAGACCCACGCTCACCGGCTCCAGATTTATCAGCAATAAACCAGCCAGCCGGAA
GGGCCGAGCGCAGAAGTGGTCCTGCAACTTTATCCGCCTCCATCCAGTCTATTAATTGTT
GCCGGGAAGCTAGAGTAAGTAGTTCGCCAGTTAATAGTTTGCGCAACGTTGTTGCCATTG
CTACAGGCATCGTGGTGTCACGCTCGTCGTTTGGTATGGCTTCATTCAGCTCCGGTTCCC
AACGATCAAGGCGAGTTACATGATCCCCCATGTTGTGCAAAAAAGCGGTTAGCTCCTTCG
GTCCTCCGATCGTTGTCAGAAGTAAGTTGGCCGCAGTGTTATCACTCATGGTTATGGCAG
CACTGCATAATTCTCTTACTGTCATGCCATCCGTAAGATGCTTTTCTGTGACTGGTGAGT
ACTCAACCAAGTCATTCTGAGAATAGTGTATGCGGCGACCGAGTTGCTCTTGCCCGGCGT
CAATACGGGATAATACCGCGCCACATAGCAGAACTTTAAAAGTGCTCATCATTGGAAAAC
GTTCTTCGGGGCGAAAACTCTCAAGGATCTTACCGCTGTTGAGATCCAGTTCGATGTAAC
CCACTCGTGCACCCAACTGATCTTCAGCATCTTTTACTTTCACCAGCGTTTCTGGGTGAG
CAAAAACAGGAAGGCAAAATGCCGCAAAAAAGGGAATAAGGGCGACACGGAAATGTTGAA
TACTCATACTCTTCCTTTTTCAATATTATTGAAGCATTTATCAGGGTTATTGTCTCATGA
GCGGATACATATTTGAATGTATTTAGAAAAATAAACAAATAGGGGTTCCGCGCACATTTC
CCCGAAAAGTGCCACCTAAATTGTAAGCGTTAATATTTTGTTAAAATTCGCGTTAAATTT
TTGTTAAATCAGCTCATTTTTTAACCAATAGGCCGAAATCGGCAAAATCCCTTATAAATC
AAAAGAATAGACCGAGATAGGGTTGAGTGTTGTTCCAGTTTGGAACAAGAGTCCACTATT
AAAGAACGTGGACTCCAACGTCAAAGGGCGAAAAACCGTCTATCAGGGCGATGGCCCACT
ACGTGAACCATCACCCTAATCAAGTTTTTTGGGGTCGAGGTGCCGTAAAGCACTAAATCG
GAACCCTAAAGGGAGCCCCCGATTTAGAGCTTGACGGGGAAAGCCGGCGAACGTGGCGAG
AAAGGAAGGGAAGAAAGCGAAAGGAGCGGGCGCTAGGGCGCTGGCAAGTGTAGCGGTCAC
GCTGCGCGTAACCACCACACCCGCCGCGCTTAATGCGCCGCTACAGGGCGCGTCCCATTC
GCCATTCAGGCTGCGCAACTGTTGGGAAGGGCGATCGGTGCGGGCCTCTTCGCTATTACG
CCAGCTGGCGAAAGGGGGATGTGCTGCAAGGCGATTAAGTTGGGTAACGCCAGGGTTTTC
CCAGTCACGACGTTGTAAAACGACGGCCAGTGAATTGTAATACGACTCACTATAGGGCGA
ATTGGGTACCCCATTTCTACTTATCAAAATTGATGTATTTTCTTGAAGAATAAATCCATT
CATCATGTAGGTCCATAAGAACGGCTCCAATTAAGCGATTGGCTGATGTTTGATTGGGGA
AGATGCGAATAATCTTTTCTCTTCTGCGTACTTCTTGATTCAGTCGTTCAATTAGATTGG
TACTCTTTAGTCGATTGTGGGAATTTCCTTGTACGGTATATTGAAAGGCGTCTTCGAATC
CATCATCCAATGATGCGCAAGCTTTTGAATATTTTGGTTGATCGATATAATCATGAATCA
ATCGATTTTTAGCCTCACGCGCTAAGTTAATATCTGTGAACTTAAAAATTCCTTTAACAG
CTTCTCTGAAAGATTTTGAATTTTTTTTAGGAATGGTGGTAAAGATATTTCTTAGGAAGT
GAACTTGGCATCTTTGCCAACTTACGTTGGTGAAGGATTTTCTAATGGCAGAGACTAATC
CTTTGTGCGCATCAGAAATAACGAGTTCCGTACCTTGTAAACCGCGTTCTTTTAGGTATT
CAAAAAATGTTGTCCAGGTCTCTTCGCTTTCGCCACTTTGAATCATGAAGCCGATAATTT
CACGGTCGCCATCTTTGGTTATTCCAATCGCTATATGACAGCTTTTTGAGAGTACTCGAT
TTTCTTCTCGTACTTTTATATAGAGTACATCGGTCATTAAGTAAGGATAATTTTTTTCTG
ATAATAAACGATTCTGCCACTCGTTAACCATAGGTTCTAGCTGTTCTGTTAAGCTAGAAA
CGAAGGACTTAGAGACGGATTTACCACAAAGTTCTTCCACAATTTTTGATACTTTACGAG
TTGAAACGCCTGATACATACATTTCCAACATTGAAGCCATGAGGGCTTTTTCGTTTCGTT
GATAACGTTCAAACACTGTGGGTGAAAAATGGCCATCACGTGTTCTGGGTACTTTTAATT
CTAGCGTGCCTACACGTGTCGTAAAGCTGCGCTCATAATAGCCATTTCGTTGACTTTGTC
GGTTTTCTGTTCGTTCATATTCTTTTGCTTGAATATATTCTGTTCGTTGATTTTCCATTA
GTTGATTAAATACCGTTGTTAAAATATTTTTAGAAACGTCATCCTTTACAGAATATTCAA
TAATGCTTTGAATCTCTTCGCTTTTCAGTGTAAAATGTACTTGGGTCATGTAAAAGTCCT
CCTGGGTATGTTTTTGTCGTTAAAAACATTGTACCGTAAAAGGACTGTTATATGGCCTTT
TTACTTTTACACGTCGACGGTATCGATAAGCTTGATAAAGTCCGTATAATTGTGTAAAAG
AATTCGCTTGCATGCCTGCAGAATTAAAAGTTAGTGAACAAGAAAACAGTGAAGCACCAG
TTTCTGAACCAAAAGAAGACGAAAAAACAAAAAAAGATTAAGCAATTTATTTGGAAAATC
TTTTTTTGTTTTTTTAAGAAATATTTATTGTTTTTTTTAAAAAATTATTGTACAGTTGCT
ACTATAAGGGAAAGAAAAAAAGAAAGATATAAATTGTATAAAGTAGGGTTAGAAGCAATT
AATAATTATTATTAATGTTATTTTTCTCTTATATATTCAATGTAATTTTAATTACATTTG
CTTTTAATAAAAACACTACTTAATAGAGAAAGGAAATATAAGATCTCATATGTCTAGATT
AGATAAAAGTAAAGTGATTAACAGCGCATTAGAGCTGCTTAATGAGGTCGGAATCGAAGG
TTTAACAACCCGTAAACTCGCCCAGAAGCTAGGTGTAGAGCAGCCTACATTGTATTGGCA
TGTAAAAAATAAGCGGGCTTTGCTCGACGCCTTAGCCATTGAGATGTTAGATAGGCACCA
TACTCACTTTTGCCCTTTAGAAGGGGAAAGCTGGCAAGATTTTTTACGTAATAACGCTAA
AAGTTTTAGATGTGCTTTACTAAGTCATCGCGATGGAGCAAAAGTACATTTAGGTACACG
GCCTACAGAAAAACAGTATGAAACTCTCGAAAATCAATTAGCCTTTTTATGCCAACAAGG
TTTTTCACTAGAGAATGCATTATATGCACTCAGCGCTGTGGGGCATTTTACTTTAGGTTG
CGTATTGGAAGATCAAGAGCATCAAGTCGCTAAAGAAGAAAGGGAAACACCTACTACTGA

TAGTATGCCGCCATTATTACGACAAGCTATCGAATTATTTGATCACCAAGGTGCAGAGCC
AGCCTTCTTATTCGGCCTTGAATTGATCATATGCGGATTAGAAAAACAACTTAAATGTGA
AAGTGGGTCTTAAGGATCTGAACTGCAGGTCGATggatccCCGCCTCCAGCAGAATTAAA
AGTTAGTGAACAAGAAAACAGTGAAGCACCAGTTTCTGAACCAAAAGAAGACGAAAAAAC
AAAAAAAGATTAAGCAATTTATTTGGAAAATCTTTTTTTGTTTTTTTAAGAAATATTTAT
TGTTTTTTTTAAAAAATTATGTACAATTGCTACTATAAGGGAAAGAAAAAAAGAAAGATA
TAAATTGTATAAAGTAGGGTTAGAAGCAATTAATAATTATTATTAATGTTATTTTTCTCT
TATATATTCAATGTAATTTTAATTACATTTGCTTTTAATAAAAACACTACTTAATAGAGA
AAGGAAATATAACCTAGGGTCTCCTCTTAGAGGAGTTAGCTTAGTTTTAGAGCTAGAAAT
AGCAAGTTAAAATAAGGCTAGTCCGTTATCAACTTGAAAAAGTGGCACCGAGTCGGTGCT
TTTTTTACGGATCTAAATTAAAGTTGGTTCATTCAAAGTTACAATTCGTTTAAAAAGTGA
ATAATTAATTATTAATTAATTTATTAAAAAACAACCAAAAGGTTGTTTTTTATTTTTTAA
AGGTCGGTCCGTAATACGACTCACTTAAGGCCTTGACTAGAGGGTACCAATTCCAGAGTC
TCCATATATGAATAATGGATTTCATTTTTTTCCAAGATCTAAGAAATGAAAAAACTTTTA
TCGATCAAAAACTAAAAAAAATATTGACACTCTATCATTGATAGAGTATAATTAACGGGA
TCCTCTATCATTGATAGAGGGATCCCGCCAAGCTTGGGATCCCCAGCTTGTTGATACACT
AATGCTTTTATATAGGGAAAAGGTGGTGAACTACTATGGACAAAAAATATTCTATTGGAT
TAGCTATTGGAACTAATTCAGTAGGTTGAGCTGTTATTACAGATGAATATAAGGTACCAT
CAAAGAAATTTAAGGTTTTAGGTAATACTGATAGACATTCAATTAAAAAAAATTTAATCG
GAGCATTACTTTTTGATTCTGGAGAAACAGCAGAAGCTACCAGATTAAAAAGAACGGCTC
GTAGACGATATACTAGACGTAAAAACAGAATCTGTTATCTTCAAGAAATTTTTAGTAATG
AAATGGCTAAAGTTGATGACTCTTTTTTTCACAGATTGGAAGAGTCTTTCTTAGTAGAAG
AAGATAAAAAGCATGAAAGACACCCCATCTTTGGTAATATTGTAGATGAAGTCGCCTATC
ATGAAAAATATCCTACAATTTATCATTTAAGAAAAAAATTAGTAGATAGCACAGATAAAG
CTGATTTAAGATTAATTTATTTAGCACTAGCACATATGATCAAATTTAGAGGTCACTTCT
TAATTGAAGGTGATTTAAACCCTGATAATAGTGACGTTGATAAATTATTTATTCAATTAG
TACAAACGTATAATCAACTTTTCGAAGAAAACCCAATTAATGCTAGTGGGGTTGATGCAA
AAGCTATCCTTTCGGCTCGTCTTTCAAAATCTAGGAGACTTGAAAACTTAATTGCACAAT
TACCGGGAGAGAAAAAAAACGGTTTATTTGGTAACTTAATCGCGTTATCTTTAGGTTTAA
CCCCGAATTTTAAGAGTAATTTCGATCTAGCTGAAGATGCTAAACTACAATTATCTAAAG
ATACTTATGACGATGACTTAGATAATTTATTAGCTCAGATTGGTGATCAATATGCAGACT
TATTTTTAGCAGCAAAAAACTTAAGCGACGCAATCTTATTGAGTGATATATTGAGAGTTA
ACACAGAAATCACTAAAGCACCATTAAGTGCAAGTATGATTAAACGTTATGATGAACACC
ACCAAGATTTAACACTATTAAAAGCATTAGTTAGACAACAATTACCTGAAAAGTATAAAG
AAATTTTCTTCGATCAAAGCAAAAATGGTTATGCTGGTTATATTGATGGTGGAGCTTCAC
AAGAAGAATTTTATAAGTTCATTAAGCCTATCCTAGAAAAAATGGATGGAACAGAAGAAC
TATTAGTCAAGTTAAATCGTGAAGATTTACTACGCAAACAAAGAACTTTTGATAATGGTA
GCATTCCTCATCAAATTCACTTAGGAGAACTACACGCTATCCTAAGAAGACAAGAAGATT
TTTATCCTTTTTTAAAAGATAATAGAGAAAAAATTGAAAAAATCTTAACATTTAGAATCC
CTTACTATGTAGGTCCGTTAGCTAGAGGAAATAGTAGATTTGCATGAATGACTCGAAAAT
CAGAAGAGACTATCACACCATGAAATTTTGAGGAAGTTGTGGATAAAGGTGCATCTGCGC
AATCTTTTATTGAGCGAATGACTAATTTCGATAAGAACTTACCTAATGAAAAAGTATTAC
CTAAGCACTCATTATTATATGAATACTTTACTGTTTATAACGAACTTACTAAAGTAAAAT
ATGTTACCGAAGGAATGAGAAAACCAGCGTTCCTAAGTGGAGAACAAAAGAAGGCTATTG
TTGATTTATTATTTAAGACAAATAGAAAAGTAACTGTAAAACAACTAAAAGAAGATTATT
TTAAAAAAATTGAATGTTTTGATTCAGTCGAAATTTCTGGAGTTGAAGACCGTTTCAACG
CAAGTTTAGGCACTTACCACGATCTACTAAAAATTATTAAAGATAAAGATTTTCTTGATA
ACGAAGAAAATGAAGACATTCTAGAAGATATTGTCCTAACTTTAACTTTATTCGAAGACA
GAGAAATGATTGAAGAAAGATTAAAAACTTACGCTCACTTATTTGATGATAAAGTTATGA
AGCAGTTGAAGCGCCGACGATATACCGGTTGAGGTAGACTCTCAAGAAAGCTAATCAATG
GTATTAGAGACAAACAATCAGGTAAAACAATTTTAGATTTTTTAAAAAGCGACGGATTTG
CTAATAGAAACTTCATGCAATTGATCCACGA

### pMT85_SpdCas9_pmcDA1

ATGCTATTGTACCTCAATCATTTCTAAAAGATGATTCTATCGACAATAAGGTTTTAACTA
GATCCGATAAAAATCGAGGAAAATCTGACAATGTACCTAGTGAAGAAGTTGTAAAAAAGA
TGAAAAATTACTGACGACAACTTTTAAACGCAAAATTAATTACACAAAGAAAATTTGATA
ACCTAACTAAAGCAGAGCGTGGAGGTCTGTCTGAACTTGATAAGGCTGGATTTATTAAAC
GACAACTAGTTGAAACACGTCAAATCACCAAACATGTTGCACAAATTTTAGATTCTCGTA
TGAATACAAAGTACGACGAAAACGATAAATTAATCAGAGAAGTTAAAGTTATTACTCTAA
AATCAAAATTAGTGAGTGATTTTCGCAAAGATTTCCAATTTTACAAGGTTAGAGAGATCA
ATAATTATCATCACGCACATGATGCTTATTTAAATGCTGTGGTTGGGACTGCTTTAATCA
AAAAGTATCCTAAATTAGAAAGCGAATTTGTATACGGTGATTATAAGGTTTATGATGTTC
GTAAAATGATTGCTAAAAGTGAACAAGAAATTGGAAAGGCTACTGCTAAATATTTTTTTT
ACTCAAATATTATGAATTTTTTCAAAACTGAGATCACATTAGCAAATGGTGAAATCCGTA
AAAGACCTTTAATTGAAACTAACGGGGAAACTGGGGAAATTGTGTGAGATAAAGGTCGTG
ATTTTGCAACAGTAAGAAAAGTATTATCAATGCCACAAGTTAATATCGTTAAAAAAACAG
AGGTGCAAACTGGAGGTTTCTCTAAAGAATCGATCTTACCTAAAAGAAACAGTGATAAAT
TAATTGCGAGAAAAAAAGATTGAGATCCAAAAAAATATGGTGGTTTTGATTCGCCGACTG
TTGCATACTCTGTATTAGTGGTTGCTAAAGTTGAAAAAGGTAAGAGTAAAAAATTAAAAT
CTGTTAAAGAATTATTAGGCATTACGATTATGGAGAGATCATCTTTTGAAAAAAATCCAA
TCGATTTTTTGGAAGCTAAAGGGTACAAAGAAGTAAAAAAAGATCTTATAATTAAATTAC
CTAAATATTCTTTATTTGAACTTGAAAACGGTCGTAAAAGAATGTTAGCTTCTGCAGGAG
AACTACAAAAAGGTAATGAATTAGCTCTACCAAGTAAATATGTAAATTTTTTATATTTAG
CGAGTCACTATGAAAAATTAAAAGGATCACCTGAAGATAATGAACAAAAACAATTATTTG
TTGAACAACACAAACATTATTTAGATGAAATTATTGAACAAATTTCAGAATTTAGCAAAA
GAGTTATACTTGCCGATGCAAATTTAGATAAAGTGTTAAGTGCGTACAACAAGCACAGAG

ATAAACCAATCCGCGAACAAGCAGAAAATATTATCCACTTATTCACTCTTACTAACTTAG
GTGCTCCAGCAGCTTTTAAATATTTTGATACAACTATCGATAGAAAGAGATATACATCAA
CAAAGGAAGTTTTAGACGCGACTTTAATTCATCAAAGTATTACTGGACTTTATGAGACTC
GCATCGATCTATCACAATTGGGTGGAGACGGAGGTGGTGGAACTGGTGGTGGTGGTAGTG
CTGAATATGTTAGAGCTTTATTTGATTTTAATGGTAATGATGAAGAAGATTTACCATTTA
AGAAAGGTGATATTTTAAGAATTAGAGATAAACCAGAAGAACAATGGTGGAATGCTGAAG
ATAGTGAAGGTAAAAGAGGTATGATTTTAGTTCCATATGTTGAGAAATATAGTGGTACTG
ATGCTGAATATGTTAGAATTCATGAGAAATTAGATATTTATACTTTTAAGAAACAATTCT
TTAATAATAAGAAGAGTGTTAGTCATCGTTGCTATGTGCTCTTTGAATTAAAGAGAAGAG
GTGAAAGAAGAGCTTGCTTTTGAGGATATGCTGTTAATAAACCACAAAGTGGAACTGAAA
GAGGAATTCATGCTGAAATATTTAGTATTAGAAAAGTTGAAGAATATTTAAGAGATAATC
CAGGACAATTTACTATTAATTGGTATAGTAGTTGGAGTCCATGTGCTGATTGTGCTGAGA
AGATTTTAGAATGGTATAATCAAGAATTAAGAGGAAATGGACATACTTTGAAAATTTGGG
CTTGTAAATTATATTATGAGAAGAATGCTAGAAATCAAATTGGATTATGGAATTTAAGAG
ATAATGGAGTTGGATTAAATGTTATGGTTAGTGAACATTATCAATGTTGTAGAAAGATAT
TTATTCAAAGTAGTCATAATCAATTAAATGAAAATAGATGGTTAGAGAAAACTTTAAAGA
GAGCTGAGAAAAGAAGAAGTGAATTAAGTATTATGATTCAAGTTAAAATTTTACATACTA
CTAAAAGTCCAGCTGTTAGTGGAGGAAGTACTAATTTAAGTGATATTATTGAGAAAGAAA
CTGGAAAACAATTAGTTATTCAAGAAAGTATTTTAATGTTACCAGAAGAAGTTGAAGAAG
TTATTGGAAATAAACCAGAAAGTGATATTTTAGTTCATACTGCTTATGATGAAAGTACTG
ATGAAAATGTTATGTTATTAACTAGTGATGCTCCAGAATATAAACCATGGGCTTTAGTTA
TTCAAGATAGTAATGGAGAAAATAAAATTAAAATGTTATAAGATCTAAATTAAAGTTGGT
TCATTCAAAGTTACAATTCGTTTAAAAAGTGAATAATTAATTATTAATTAATTTATTAAA
AAACAACCAAAAGGTTGTTTTTTATTTTTTAAAGAAGGATGGTCTCCAGACACGTTGCCT
GGTTTCCGGCACCAGAAGCGGTGCCGGAAAGCTGGCTGGAGTGCGATCTTCCTGAGGCCG
ATACTGTCGTCGTCCCCTCAAACTGGCAGATGCACGGTTACGATGCGCCCATCTACACCA
ACGTGACCTATCCCATTACGGTCAATCCGCCGTTTGTTCCCACGGAGAATCCGACGGGTT
GTTACTGCGCTCACATTTATATGYTGACTGAAAGCTGGCTACAGGAAGGCCAGACGCGAA
TTATTTTTGATGGCTAGAAGCTTTAGGATGAATGGATTTATTCTTCAAGAAAATACATCA
ATTTTGATAAGTAGAAATGGTAAAAACATTGTATAGCATTTTACACAGGAGTCTGGACTT
GACTGAGTTTATGGAAGAAGTTTTAATTGATGATAATATGGTTTTTGATATTGATAATTT
AAAAGGATTTCTTAATGATACCAGTTCAATTTGGGTTTATAGCTAAAGAAAATAATAAAA
TTATAGGATTTGCATATTGCTATACACTTTTAAGACCTGATGGAAAAACAATGTTTTATT
TACACTCAATAGGAATGTTACCTAACTATCAAGACAAAGGTTATGGTTCAAAATTATTAT
CTTTTATTAAGGAATATTCTAAAGAGATTGGTTGTTCTGAAATGTTTTTAATAACTGATA
AAGGTAATCCTAGAGCTTGCCATGTATATGAAAAATTAGGTGGTAAAAATGATTATAAAG
ATGAAATAGTATATGTATATGATTATGAAAAAGGTGATAAATAAATGAATATAGTTGAAA
ATGAAATATGTATAAGAACTTTAATAGATGATGATTTTCCTTTGATGTTAAAATGGTTAA
CTGATGAAAGAGTATTAGAATTTTATGGTGGTAGAGATAAAAAATATACATTAGAATCAT
TAAAAAAACATTATACAGAGCCTTGGGAAGATGAAGTTTTTAGAGTAATTATTGAATATA
ACAATGTTCCTATTGGATATGGACAAATATATAAAATGTATGATGAGTTATATACTGATT
ATCATTATCCAAAAACTGATGAGATAGTCTATGGTATGGATCAATTTATAGGAGAGCCAA
ATTATTGGAGTAAAGGAATTGGTACAAGATATATTAAATTGATTTTTGAATTTTTGAAAA
AAGAAAGAAATGCTAATGCAGTTATTTTAGACCCTCATAAAAATAATCCAAGAGCAATAA
GGGCATACCAAAAATCTGGTTTTAGAATTATTGAAGATTTGCCAGAACATGAATTACACG
AGGGCAAAAAAGAAGATTGTTATTTAATGGAATATAGATATGATGATAATGCCACAAATG
TTAAGGCAATGAAATATTTAATTGAGCATTACTTTGATAATTTCAAAGTAGATAGTATTG
AAATAATCGGTAGTGGTTATGATAGTGTGGCATATTTAGTTAATAATGAATACATTTTTA
AAACAAAATTTAGTACTAATAAGAAAAAAGGTTATGCAAAAGAAAAAGCAATATATAATT
TTTTAAATACAAATTTAGAAACTAATGTAAAAATTCCTAATATTGAATATTCGTATATTA
GTGATGAATTATCTATACTAGGTTATAAAGAAATTAAAGGAACTTTTTTAACACCAGAAA
TTTATTCTACTATGTCAGAAGAAGAACAAAATTTGTTAAAACGAGATATTGCCAGTTTTT
TAAGACAAATGCACGGTTTAGATTATACAGATATTAGTGAATGTACTATTGATAATAAAC
AAAATGTATTAGAAGAGTATATATTGTTGCGTGAAACTATTTATAATGATTTAACTGATA
TAGAAAAAGATTATATAGAAAGTTTTATGGAAAGACTAAATGCAACAACAGTTTTTGAGG
GTAAAAAGTGTTTATGCCATAATGATTTTAGTTGTAATCATCTATTGTTAGATGGCAATA
ATAGATTAACTGGAATAATTGATTTTGGAGATTCTGGAATTATAGATGAATATTGTGATT
TTATATACTTACTTGAAGATAGTGAAGAAGAAATAGGAACAAATTTTGGAGAAGATATAT
TAAGAATGTATGGAAATATAGATATTGAGAAAGCAAAAGAATATCAAGATATAGTTGAAG
AATATTATCCTATTGAAACTATTGTTTATGGAATTAAAAATATTAAACAGGAATTTATCG
AAAATGGTAGAAAAGAAATTTATAAAAGGACTTATAAAGATTRATAAAGATCTACGAAGG
CATGACCAAAATCCCTTAACGTGAGTTTTCGTTCCACTGAGCGTCAGACCCCGTAGAAAA
GATCAAAGGATCTTCTTGAGATCCTTTTTTTCTGCGCGTAATCTGCTGCTTGCAAACAAA
AAAACCACCGCTACCAGCGGTGGTTTGTTTGCCGGATCAAGAGCTACCAACTCTTTTTCC
GAAGGTAACTGGCTTCAGCAGAGCGCAGATACCAAATACTGTTCTTCTAGTGTAGCCGTA
GTTAGGCCACCACTTCAAGAACTCTGTAGCACCGCCTACATACCTCGCTCTGCTAATCCT
GTTACCAGTGGCTGCTGCCAGTGGCGATAAGTCGTGTCTTACCGGGTTGGACTCAAGACG
ATAGTTACCGGATAAGGCGCAGCGGTCGGGCTGAACGGGGGGTTCGTGCACACAGCCCAG
CTTGGAGCGAACGACCTACACCGAACTGAGATACCTACAGCGTGAGCTATGAGAAAGCGC
CACGCTTCCCGAAGGGAGAAAGGCGGACAGGTATCCGGTAAGCGGCAGGGTCGGAACAGG
AGAGCGCACGAGGGAGCTTCCAGGGGGAAACGCCTGGTATCTTTATAGTCCTGTCGGGTT
TCGCCACCTCTGACTTGAGCGTCGATTTTTGTGATGCTCGTCAGGGGGGCGGAGCCTATG
GAAAAACGCCAGCAACGCGGCCTTTTTACGGTTCCTGGCCTTTTGCTGGCCTTTTGCTCA
CATGTTCTTTCCTGCGTTATCCCCTGATTCTGTGGATAACCGTATTACCGCCTTTGAGTG
AGCTGATACCGCTCGCCGCAGCCGAACGACCGAGCGCAGCGAGTCAGTGAGCGAGGAAGC
GGAAGAGCGCCCAATACGCAAACCGCCTCTCCCCGCGCGTTGGCCGATTCATTAATGCAC
GCTAGCGGATCTCATAAAAATGTATCCTAAATCAAATATCGGACAAGCAGTGTCTGTTAT

AACAAAAAATCGATTTAATAGACACATTAACAGCACTGTTTTTATGTGTGCGATAATTTA
TAATATTTCGGACGGTTGCGGTACCCTTTTACACAATTATACGGACTTTATCCTGCAGGG
GCCCAATTGTGTAAAAGTAAAAAGGCCATATAACAGTCCTTTTACGGTACAATGTTTTTA
ACGACAAAAACATACCCAGGAGGACTTTTACATGACCCAAGTACATTTTACACTGAAAAG
CGAAGAGATTCAAAGCATTATTGAATATTCTGTAAAGGATGACGTTTCTAAAAATATTTT
AACAACGGTATTTAATCAACTAATGGAAAATCAACGAACAGAATATATTCAAGCAAAAGA
ATATGAACGAACAGAAAACCGACAAAGTCAACGAAATGGCTATTATGAGCGCAGCTTTAC
GACACGTGTAGGCACGCTAGAATTAAAAGTACCCAGAACACGTGATGGCCATTTTTCACC
CACAGTGTTTGAACGTTATCAACGAAACGAAAAAGCCCTCATGGCTTCAATGTTGGAAAT
GTATGTATCAGGCGTTTCAACTCGTAAAGTATCAAAAATTGTGGAAGAACTTTGTGGTAA
ATCCGTCTCTAAGTCCTTCGTTTCTAGCTTAACAGAACAGCTAGAACCTATGGTTAACGA
GTGACAGAATCGTTTATTATCAGAAAAAAATTATCCTTACTTAATGACCGATGTACTCTA
TATAAAAGTACGAGAAGAAAATCGAGTACTCTCAAAAAGCTGTCATATAGCGATTGGAAT
AACCAAAGATGGCGACCGTGAAATTATCGGCTTCATGATTCAAAGTGGCGAAAGCGAAGA
GACCTGGACAACATTTTTTGAATACCTAAAAGAACGCGGTTTACAAGGTACGGAACTCGT
TATTTCTGATGCGCACAAAGGATTAGTCTCTGCCATTAGAAAATCCTTCACCAACGTAAG
TTGGCAAAGATGCCAAGTTCACTTCCTAAGAAATATCTTTACCACCATTCCTAAAAAAAA
TTCAAAATCTTTCAGAGAAGCTGTTAAAGGAATTTTTAAGTTCACAGATATTAACTTAGC
GCGTGAGGCTAAAAATCGATTGATTCATGATTATATCGATCAACCAAAATATTCAAAAGC
TTGCGCATCATTGGATGATGGATTCGAAGACGCCTTTCAATATACCGTACAAGGAAATTC
CCACAATCGACTAAAGAGTACCAATCTAATTGAACGACTGAATCAAGAAGTACGCAGAAG
AGAAAAGATTATTCGCATCTTCCCCAATCAAACATCAGCCAATCGCTTAATTGGAGCCGT
TCTTATGGACCTACATGATGAATGGATTTATTCTTCAAGAAAATACATCAATTTTGATAA
GTAGAAATGGTAAAAACATTGTATAGCATTTTACACAGGAGTCTGGACTTGACTCACTTC
CTTTATTATTTTTCATTTTTTTGACCTCGAGGGGGGGCCCACCATACAGCTGACGATAAA
GTCCGTATAATTGTGTAAAAACCCATAGCTTTGGACACACACTAGTGGATCTCATAAAAA
TGTATCCTAAATCAAATATCGGACAAGCAGTGTCTGTTATAACAAAAAATCGATTTAATA
GACACATTAACAGCACTGTTTTTATGTGTGCGATAATTTATAATATTTCGGACGGTTGCG
GATCCACCCGCAATTACTGTGAGTTAGCTCACTCATTAGGCACCCCAGGCTTTACACTTT
ATACTTCCGGCTCGTATATTGTGTGGAATTGTGAGCGGATAACAATTTCACACAGGAAAC
AGCTATGACCTTGATTACGGAATTCACGGCCGGGGGGGCCACCCCACCAATTGACGCGGC
CGCAACTCTAGAGGATTCATCGGCCGTCGAATTCGCTTGCATGCCTGCAGAATTAAAAGT
TAGTGAACAAGAAAACAGTGAAGCACCAGTTTCTGAACCAAAAGAAGACGAAAAAACAAA
AAAAGATTAAGCAATTTATTTGGAAAATCTTTTTTTGTTTTTTTAAGAAATATTTATTGT
TTTTTTTAAAAAATTATTGTACAGTTGCTACTATAAGGGAAAGAAAAAAAGAAAGATATA
AATTGTATAAAGTAGGGTTAGAAGCAATTAATAATTATTATTAATGTTATTTTTCTCTTA
TATATTCAATGTAATTTTAATTACATTTGCTTTTAATAAAAACACTACTTAATAGAGAAA
GGAAATATAAGATCTCATATGTCTAGATTAGATAAAAGTAAAGTGATTAACAGCGCATTA
GAGCTGCTTAATGAGGTCGGAATCGAAGGTTTAACAACCCGTAAACTCGCCCAGAAGCTA
GGTGTAGAGCAGCCTACATTGTATTGGCATGTAAAAAATAAGCGGGCTTTGCTCGACGCC
TTAGCCATTGAGATGTTAGATAGGCACCATACTCACTTTTGCCCTTTAGAAGGGGAAAGC
TGGCAAGATTTTTTACGTAATAACGCTAAAAGTTTTAGATGTGCTTTACTAAGTCATCGC
GATGGAGCAAAAGTACATTTAGGTACACGGCCTACAGAAAAACAGTATGAAACTCTCGAA
AATCAATTAGCCTTTTTATGCCAACAAGGTTTTTCACTAGAGAATGCATTATATGCACTC
AGCGCTGTGGGGCATTTTACTTTAGGTTGCGTATTGGAAGATCAAGAGCATCAAGTCGCT
AAAGAAGAAAGGGAAACACCTACTACTGATAGTATGCCGCCATTATTACGACAAGCTATC
GAATTATTTGATCACCAAGGTGCAGAGCCAGCCTTCTTATTCGGCCTTGAATTGATCATA
TGCGGATTAGAAAAACAACTTAAATGTGAAAGTGGGTCTTAAGGATCTGAACTGCAGGTC
GATggatccCCGCCTCCAGCAGAATTAAAAGTTAGTGAACAAGAAAACAGTGAAGCACCA
GTTTCTGAACCAAAAGAAGACGAAAAAACAAAAAAAGATTAAGCAATTTATTTGGAAAAT
CTTTTTTTGTTTTTTTAAGAAATATTTATTGTTTTTTTTAAAAAATTATGTACAATTGCT
ACTATAAGGGAAAGAAAAAAAGAAAGATATAAATTGTATAAAGTAGGGTTAGAAGCAATT
AATAATTATTATTAATGTTATTTTTCTCTTATATATTCAATGTAATTTTAATTACATTTG
CTTTTAATAAAAACACTACTTAATAGAGAAAGGAAATATAACCTAGGGTCTCCTCTTAGA
GGAGTTAGCTTAGTTTTAGAGCTAGAAATAGCAAGTTAAAATAAGGCTAGTCCGTTATCA
ACTTGAAAAAGTGGCACCGAGTCGGTGCTTTTTTTACGGATCTAAATTAAAGTTGGTTCA
TTCAAAGTTACAATTCGTTTAAAAAGTGAATAATTAATTATTAATTAATTTATTAAAAAA
CAACCAAAAGGTTGTTTTTTATTTTTTAAAGGTCGGTCCGTAATACGACTCACTTAAGGC
CTTGACTAGAGGGTACCAATTCCAGAGTCTCCATATATGAATAATGGATTTCATTTTTTT
CCAAGATCTAAGAAATGAAAAAACTTTTATCGATCAAAAACTAAAAAAAATATTGACACT
CTATCATTGATAGAGTATAATTAACGGGATCCTCTATCATTGATAGAGGGATCCCGCCAA
GCTTGGGATCCCCAGCTTGTTGATACACTAATGCTTTTATATAGGGAAAAGGTGGTGAAC
TACTATGGACAAAAAATATTCTATTGGATTAGCTATTGGAACTAATTCAGTAGGTTGAGC
TGTTATTACAGATGAATATAAGGTACCATCAAAGAAATTTAAGGTTTTAGGTAATACTGA
TAGACATTCAATTAAAAAAAATTTAATCGGAGCATTACTTTTTGATTCTGGAGAAACAGC
AGAAGCTACCAGATTAAAAAGAACGGCTCGTAGACGATATACTAGACGTAAAAACAGAAT
CTGTTATCTTCAAGAAATTTTTAGTAATGAAATGGCTAAAGTTGATGACTCTTTTTTTCA
CAGATTGGAAGAGTCTTTCTTAGTAGAAGAAGATAAAAAGCATGAAAGACACCCCATCTT
TGGTAATATTGTAGATGAAGTCGCCTATCATGAAAAATATCCTACAATTTATCATTTAAG
AAAAAAATTAGTAGATAGCACAGATAAAGCTGATTTAAGATTAATTTATTTAGCACTAGC
ACATATGATCAAATTTAGAGGTCACTTCTTAATTGAAGGTGATTTAAACCCTGATAATAG
TGACGTTGATAAATTATTTATTCAATTAGTACAAACGTATAATCAACTTTTCGAAGAAAA
CCCAATTAATGCTAGTGGGGTTGATGCAAAAGCTATCCTTTCGGCTCGTCTTTCAAAATC
TAGGAGACTTGAAAACTTAATTGCACAATTACCGGGAGAGAAAAAAAACGGTTTATTTGG
TAACTTAATCGCGTTATCTTTAGGTTTAACCCCGAATTTTAAGAGTAATTTCGATCTAGC
TGAAGATGCTAAACTACAATTATCTAAAGATACTTATGACGATGACTTAGATAATTTATT
AGCTCAGATTGGTGATCAATATGCAGACTTATTTTTAGCAGCAAAAACTTAAGCGACGC

AATCTTATTGAGTGATATATTGAGAGTTAACACAGAAATCACTAAAGCACCATTAAGTGC
AAGTATGATTAAACGTTATGATGAACACCACCAAGATTTAACACTATTAAAAGCATTAGT
TAGACAACAATTACCTGAAAAGTATAAAGAAATTTTCTTCGATCAAAGCAAAAAATGGTTA
TGCTGGTTATATTGATGGTGGAGCTTCACAAGAAGAATTTTATAAGTTCATTAAGCCTAT
CCTAGAAAAAATGGATGGAACAGAAGAACTATTAGTCAAGTTAAATCGTGAAGATTTACT
ACGCAAACAAAGAACTTTTGATAATGGTAGCATTCCTCATCAAATTCACTTAGGAGAACT
ACACGCTATCCTAAGAAGACAAGAAGATTTTTATCCTTTTTTAAAAGATAATAGAGAAAA
AATTGAAAAAATCTTAACATTTAGAATCCCTTACTATGTAGGTCCGTTAGCTAGAGGAAA
TAGTAGATTTGCATGAATGACTCGAAAATCAGAAGAGACTATCACACCATGAAATTTTGA
GGAAGTTGTGGATAAAGGTGCATCTGCGCAATCTTTTATTGAGCGAATGACTAATTTCGA
TAAGAACTTACCTAATGAAAAAGTATTACCTAAGCACTCATTATTATATGAATACTTTAC
TGTTTATAACGAACTTACTAAAGTAAAATATGTTACCGAAGGAATGAGAAAACCAGCGTT
CCTAAGTGGAGAACAAAAGAAGGCTATTGTTGATTTATTATTTAAGACAAATAGAAAAGT
AACTGTAAAACAACTAAAAGAAGATTATTTTAAAAAAATTGAATGTTTTGATTCAGTCGA
AATTTCTGGAGTTGAAGACCGTTTCAACGCAAGTTTAGGCACTTACCACGATCTACTAAA
AATTATTAAAGATAAAGATTTTCTTGATAACGAAGAAAATGAAGACATTCTAGAAGATAT
TGTCCTAACTTTAACTTTATTCGAAGACAGAGAAATGATTGAAGAAAGATTAAAAACTTA
CGCTCACTTATTTGATGATAAAGTTATGAAGCAGTTGAAGCGCCGACGATATACCGGTTG
AGGTAGACTCTCAAGAAAGCTAATCAATGGTATTAGAGACAAACAATCAGGTAAAACAAT
TTTAGATTTTTTAAAAAGCGACGGATTTGCTAATAGAAACTTCATGCAATTGATCCACGA
TGATTCATTAACTTTTAAAGAAGATATTCAAAAGGCACAAGTCTCAGGTCAAGGTGATAG
CTTACACGAACATATCGCTAATCTAGCTGGTTCACCTGCAATCAAAAAGGGAATTTTACA
GACAGTGAAAGTTGTTGATGAACTAGTAAAGGTAATGGGACGCCACAAACCAGAGAACAT
CGTGATTGAAATGGCTAGAGAAAACCAAACAACACAAAAAGGCCAAAAAAACAGTAGAGA
AAGAATGAAGAGAATCGAAGAAGGTATCAAAGAGTTAGGGTCTCAAATCTTAAAGGAACA
TCCTGTTGAAAACACTCAATTACAAAATGAAAAATTATACCTATATTACTTACAAAATGG
TAGAGATATGTATGTTGATCAGGAATTAGATATTAACCGTTTATCAGATTACGACGTGG

### pMYCO_SpdCas9_pmcDA1

CATTAACTTTTAAAGAAGATATTCAAAAGGCACAAGTCTCAGGTCAAGGTGATAGCTTAC
ACGAACATATCGCTAATCTAGCTGGTTCACCTGCAATCAAAAAGGGAATTTTACAGACAG
TGAAAGTTGTTGATGAACTAGTAAAGGTAATGGGACGCCACAAACCAGAGAACATCGTGA
TTGAAATGGCTAGAGAAAACCAAACAACACAAAAAGGCCAAAAAAACAGTAGAGAAAGAA
TGAAGAGAATCGAAGAAGGTATCAAAGAGTTAGGGTCTCAAATCTTAAAGGAACATCCTG
TTGAAAACACTCAATTACAAAATGAAAAATTATACCTATATTACTTACAAAATGGTAGAG
ATATGTATGTTGATCAGGAATTAGATATTAACCGTTTATCAGATTACGACGTGGATGCTA
TTGTACCTCAATCATTTCTAAAAGATGATTCTATCGACAATAAGGTTTTAACTAGATCCG
ATAAAAATCGAGGAAAATCTGACAATGTACCTAGTGAAGAAGTTGTAAAAAAGATGAAAA
ATTACTGACGACAACTTTTAAACGCAAAATTAATTACACAAAGAAAATTTGATAACCTAA
CTAAAGCAGAGCGTGGAGGTCTGTCTGAACTTGATAAGGCTGGATTTATTAAACGACAAC
TAGTTGAAACACGTCAAATCACCAAACATGTTGCACAAATTTTAGATTCTCGTATGAATA
CAAAGTACGACGAAAACGATAAATTAATCAGAGAAGTTAAAGTTATTACTCTAAAATCAA
AATTAGTGAGTGATTTTCGCAAAGATTTCCAATTTTACAAGGTTAGAGAGATCAATAATT
ATCATCACGCACATGATGCTTATTTAAATGCTGTGGTTGGGACTGCTTTAATCAAAAAGT
ATCCTAAATTAGAAAGCGAATTTGTATACGGTGATTATAAGGTTTATGATGTTCGTAAAA
TGATTGCTAAAAGTGAACAAGAAATTGGAAAGGCTACTGCTAAATATTTTTTTTACTCAA
ATATTATGAATTTTTTCAAAACTGAGATCACATTAGCAAATGGTGAAATCCGTAAAAGAC
CTTTAATTGAAACTAACGGGGAAACTGGGGAAATTGTGTGAGATAAAGGTCGTGATTTTG
CAACAGTAAGAAAAGTATTATCAATGCCACAAGTTAATATCGTTAAAAAAACAGAGGTGC
AAACTGGAGGTTTCTCTAAAGAATCGATCTTACCTAAAAGAAACAGTGATAAATTAATTG
CGAGAAAAAAAGATTGAGATCCAAAAAAATATGGTGGTTTTGATTCGCCGACTGTTGCAT
ACTCTGTATTAGTGGTTGCTAAAGTTGAAAAAGGTAAGAGTAAAAAATTAAAATCTGTTA
AAGAATTATTAGGCATTACGATTATGGAGAGATCATCTTTTGAAAAAAATCCAATCGATT
TTTTGGAAGCTAAAGGGTACAAAGAAGTAAAAAAAGATCTTATAATTAAATTACCTAAAT
ATTCTTTATTTGAACTTGAAAACGGTCGTAAAAGAATGTTAGCTTCTGCAGGAGAACTAC
AAAAAGGTAATGAATTAGCTCTACCAAGTAAATATGTAAATTTTTTATATTTAGCGAGTC
ACTATGAAAAATTAAAAGGATCACCTGAAGATAATGAACAAAAACAATTATTTGTTGAAC
AACACAAACATTATTTAGATGAAATTATTGAACAAATTTCAGAATTTAGCAAAAGAGTTA
TACTTGCCGATGCAAATTTAGATAAAGTGTTAAGTGCGTACAACAAGCACAGAGATAAAC
CAATCCGCGAACAAGCAGAAAATATTATCCACTTATTCACTCTTACTAACTTAGGTGCTC
CAGCAGCTTTTAAATATTTTGATACAACTATCGATAGAAAGAGATATACATCAACAAAGG
AAGTTTTAGACGCGACTTTAATTCATCAAAGTATTACTGGACTTTATGAGACTCGCATCG
ATCTATCACAATTGGGTGGAGACGGAGGTGGTGGAACTGGTGGTGGTGGTAGTGCTGAAT
ATGTTAGAGCTTTATTTGATTTTAATGGTAATGATGAAGAAGATTTACCATTTAAGAAAG
GTGATATTTTAAGAATTAGAGATAAACCAGAAGAACAATGGTGGAATGCTGAAGATAGTG
AAGGTAAAAGAGGTATGATTTTAGTTCCATATGTTGAGAAATATAGTGGTACTGATGCTG
AATATGTTAGAATTCATGAGAAATTAGATATTTATACTTTTAAGAAACAATTCTTTAATA
ATAAGAAGAGTGTTAGTCATCGTTGCTATGTGCTCTTTGAATTAAAGAGAAGAGGTGAAA
GAAGAGCTTGCTTTTGAGGATATGCTGTTAATAAACCACAAAGTGGAACTGAAAGAGGAA
TTCATGCTGAAATATTTAGTATTAGAAAAGTTGAAGAATATTTAAGAGATAATCCAGGAC
AATTTACTATTAATTGGTATAGTAGTTGGAGTCCATGTGCTGATTGTGCTGAGAAGATTT
TAGAATGGTATAATCAAGAATTAAGAGGAAATGGACATACTTTGAAAATTTGGGCTTGTA
AATTATATTATGAGAAGAATGCTAGAAATCAAATTGGATTATGGAATTTAAGAGATAATG
GAGTTGGATTAAATGTTATGGTTAGTGAACATTATCAATGTTGTAGAAAGATATTTATTC
AAAGTAGTCATAATCAATTAAATGAAAATAGATGGTTAGAGAAAACTTTAAAGAGAGCTG
AGAAAAGAAGAAGTGAATTAAGTATTATGATTCAAGTTAAAATTTTACATACTACTAAAA
GTCCAGCTGTTAGTGGAGGAAGTACTAATTTAAGTGATATTATTGAGAAAGAAACTGGAA

AACAATTAGTTATTCAAGAAAGTATTTTAATGTTACCAGAAGAAGTTGAAGAAGTTATTG
GAAATAAACCAGAAAGTGATATTTTAGTTCATACTGCTTATGATGAAAGTACTGATGAAA
ATGTTATGTTATTAACTAGTGATGCTCCAGAATATAAACCATGGGCTTTAGTTATTCAAG
ATAGTAATGGAGAAAATAAAATTAAAATGTTATAAGATCTAAATTAAAGTTGGTTCATTC
AAAGTTACAATTCGTTTAAAAAGTGAATAATTAATTATTAATTAATTTATTAAAAAACAA
CCAAAAGGTTGTTTTTTATTTTTTAAAGAAGGATGGTCTCCAGACACTTATAAGCCTCTC
TACTGCAATTTATCTCAACTTTGATATAATTAAAGACATACGAAAGGATTTTAATATGAC
TGAATATAAACCTACTGTTAGATTAGCTACTAGAGATGATGTTCCTAGAGCTGTTAGAAC
TTTAGCTGCTGCTTTTGCTGATTATCCTGCTACTAGACATACTGTTGATCCTGATAGACA
TATTGAAAGAGTTACTGAATTACAAGAATTATTCTTAACTAGAGTTGGTTTAGATATTGG
TAAAGTTTGGGTTGCTGATGATGGTGCTGCTGTTGCTGTTTGGACTACTCCTGAAAGTGT
TGAAGCTGGTGCTGTATTTGCTGAAATTGGTCCTAGAATGGCTGAATTAAGTGGTAGTAG
ATTAGCTGCTCAACAACAAATGGAAGGTTTATTAGCTCCACATAGACCTAAAGAACCTGC
TTGGTTCTTAGCTACTGTTGGTGTTAGTCCTGATCATCAAGGTAAAGGTTTAGGTAGTGC
TGTTGTTTTACCTGGTGTTGAAGCTGCTGAAAGAGCTGGTGTTCCTGCTTTCTTAGAAAC
TAGTGCTCCTAGAAATTTACCTTTCTATGAAAGATTAGGTTTTACTGTTACTGCTGATGT
TGAAGTTCCTGAAGGTCCTAGAACTTGGTGTATGACTAGAAAACCTGGTGCTTAATGCAG
GTCGACTCTAGAGGATCCTCAATTACTTTAGCTGCTTTTGATAAATTATCTAATAAAACT
ATTCTATTTATTGAAAAATTCATAATTACTCCTTTTTTAAAACAAATTGTTTTTACTTTA
TTTTAGAAAATAATTGGGTTTGTTATATATTACTATTTCTAATACTTATTTCTAAATTAT
TAATATTATGAGGTTAATTTGTGGATAACTGTTAATAAGTTAGGTTTAAATAGCTATTTT
TATTATTTTGTTAAAATTTTGTTCTTCAAAATATCAACAGTCTTTTTTAATTGCTTATCT
TTTTTTAACATTGTTTCTATTTTTCTTTCAGCATTAATAACTGTTGTGTGATCTCTACCA
CCAAATTCTTCTCCAATTTGAGCTAAAGTGTGATTTAAAATCTCTTTTGTTAAAAACATT
GCTATATGTCTTGCTGTTACAATTGACTTACTTCTAGCCTTTCCATCAATAGCATTAACT
GAAATACCATATTTTTCACTAACAACTTCTTTAATTTTTTTAACATTTAAAATACCTAAT
TTTGAAGTAGGTATATCTCTAAATAGATCAGAAATTATTTCTATAGTAATAATTTTTTCT
TCTGGATTTTGTTGAGATCAAAAGTTTAATCTTGAAACACTTCCTTTAATTTTTCTAACA
TCATCTGAATAATAATTAGAAATAAAATTAATTGCTTCACTAGTTACTTCTGATTTAATA
TTTTGATTTTTAATTTCTTTTTTAATGATAGCTGTTGCTGTTTTATTATCTAGTTTTTGA
ATAGCAATACTTAATCCCATATTAAATCTAGTAATTAATCTATTATCAAAACCATTTAAT
AATTCAGGAGATTTATCACTTGAAAAAAACAATTGTTTATCATTTTCTATAAAGTTATTA
AAAATAGTAAAAAATATTTCATTAGTTTTTTCTTTATAACTTAAAAATTGAACATCATCA
ATAATTAATACATCATTTTGACATACTTCATTTTTAAATTGCTCAATCTCTTTATGAGTT
TTTTGTAATATATCAACTGCTTTTCTTGCAAACTCATCACCACTCATATAACTAACTTTT
AGATCAGAAAAATTAGATTCAATATAGTTTTTTGCAGCTTTTAGTAAATGAGTTTTTCCC
ATTCCAGATTCACCATAAATAAACAATGGATTATAAGAAATTCCAGGGTTTTTGCTTGCT
GTTTGAACTGCTATAAAAGCTTGTTCATTACTTGCACCAATTACAAAATTTTCAAATGTG
TTTTCATTAATTTTTTTAACTTTTTTAGTAATGATATCAGAATGATCTTTATTAATTAGT
TCATCTTTTTCTAGTTGTTTTTTATATTCTTGTTCGTATGTAAAACTAATATTTACAGGT
TCTTTTAAAATATTTTTTATCTCATTTTCAATAGTTTGACGAAACTGTTTTATAGCTAAC
AAACCAAATTGTGATTTAACAACAACAATATAATCAGAAATCCCCTTTTTATGAATATTT
ATTGTCTTTATATAGTCGTTATACACGGATTCATCAATATTTTTATTAGCCATTAAACTT
AGTTTAAGTTCTTTTAAAATATCGTTTACGTTCATATTTGTCTCCAAAAATATTATAGTT
AAACACTTGTTAAAAATGTGGAAAACGTGGAAAAAATCCTTATAACATAGATATATTAAC
AAATTAACAAATTCTAAAATCTTATTTTAATGGTGTTATTTTATAGTTATCAACTTATCA
ACAATTAACTAACATTTTTGTTAGTAATTTGTAGTTTTATCTACATTTGATTGAGAAGCT
AAAATTTAAGTTTATAAGTGTATTTTTAAAGCAAAATATTATATAATTAGGTGTGAATTA
TTTATTTTTTTGTAAAGGAGGTAGTGGTATGAAAAGAACTTGACAACCATCAAAACTAAA
ACATGCAGGTGTTCATGGATTTAGAGCAAGAATGGCTAGGATCCCCGGGTACCGAGCTCG
AATTCGCCCTATAGTGAGTCGTATTACAATTCACTGGCCGTCGTTTTACAACGTCGTGAC
TGGGAAAACCCTGGCGTTACCCAACTTAATCGCCTTGCAGCACATCCCCCCTTCGCCAGC
TGGCGTAATAGCGAAGAGGCCCGCACCGATCGCCCTTCCCAACAGTTGCGTAGCCTGAAT
GGCGAATGGCGCGACGCGCCCTGTAGCGGCGCATTAAGCGCGGCGGGTGTGGTGGTTACG
CGCAGCGTGACCGCTACACTTGCCAGCGCCCTAGCGCCCGCTCCTTTCGCTTTCTTCCCT
TCCTTTCTCGCCACGTTCGCCGGCTTTCCCCGTCAAGCTCTAAATCGGGGGCTCCCTTTA
GGGTTCCGATTTAGTGCTTTACGGCACCTCGACCCCAAAAAACTTGATTAGGGTGATGGT
TCACGTAGTGGGCCATCGCCCTGATAGACGGTTTTTCGCCCTTTGACGTTGGAGTCCACG
TTCTTTAATAGTGGACTCTTGTTCCAAACTGGAACAACACTCAACCCTATCTCGGTCTAT
TCTTTTGATTTATAAGGGATTTTGCCGATTTCGGCCTATTGGTTAAAAAATGAGCTGATT
TAACAAAAATTTAACGCGAATTTTAACAAAATATTAACGTTTACAATTTCCTGATGCGGT
ATTTTCTCCTTACGCATCTGTGCGGTATTTCACACCGCATATGGTGCACTCTCAGTACAA
TCTGCTCTGATGCCGCATAGTTAAGCCAGCCCCGACACCCGCCAACACCCGCTGACGCGC
CCTGACGGGCTTGTCTGCTCCCGGCATCCGCTTACAGACAAGCTGTGACCGTCTCCGGGA
GCTGCATGTGTCAGAGGTTTTCACCGTCATCACCGAAACGCGCGAGACGAAAGGGCCTCG
TGATACGCCTATTTTTATAGGTTAATGTCATGATAATAATGGTTTCTTAGACGTCAGGTG
GCACTTTTCGGGGAAATGTGCGCGGAACCCCTATTTGTTTATTTTTCTAAATACATTCAA
ATATGTATCCGCTCATGAGACAATAACCCTGATAAATGCTTCAATAATATTGAAAAAGGA
AGAGTATGAGTATTCAACATTTCCGTGTCGCCCTTATTCCCTTTTTTGCGGCATTTTGCC
TTCCTGTTTTTGCTCACCCAGAAACGCTGGTGAAAGTAAAAGATGCTGAAGATCAGTTGG
GTGCACGAGTGGGTTACATCGAACTGGATCTCAACAGCGGTAAGATCCTTGAGAGTTTTC
GCCCCGAAGAACGTTTTCCAATGATGAGCACTTTTAAAGTTCTGCTATGTGGCGCGGTAT
TATCCCGTATTGACGCCGGGCAAGAGCAACTCGGTCGCCGCATACACTATTCTCAGAATG
ACTTGGTTGAGTACTCACCAGTCACAGAAAAGCATCTTACGGATGGCATGACAGTAAGAG
AATTATGCAGTGCTGCCATAACCATGAGTGATAACACTGCGGCCAACTTACTTCTGACAA
CGATCGGAGGACCGAAGGAGCTAACCGCTTTTTTGCACAACATGGGGGATCATGTAACTC
GCCTTGATCGTTGGGAACCGGAGCTGAATGAAGCCATACCAAACGACGAGCGTGACACCA

CGATGCCTGTAGCAATGGCAACAACGTTGCGCAAACTATTAACTGGCGAACTACTTACTC
TAGCTTCCCGGCAACAATTAATAGACTGGATGGAGGCGGATAAAGTTGCAGGACCACTTC
TGCGCTCGGCCCTTCCGGCTGGCTGGTTTATTGCTGATAAATCTGGAGCCGGTGAGCGTG
GGTCTCGCGGTATCATTGCAGCACTGGGGCCAGATGGTAAGCCCTCCCGTATCGTAGTTA
TCTACACGACGGGGAGTCAGGCAACTATGGATGAACGAAATAGACAGATCGCTGAGATAG
GTGCCTCACTGATTAAGCATTGGTAACTGTCAGACCAAGTTTACTCATATATACTTTAGA
TTGATTTAAAACTTCATTTTTAATTTAAAAGGATCTAGGTGAAGATCCTTTTTGATAATC
TCATGACCAAAATCCCTTAACGTGAGTTTTCGTTCCACTGAGCGTCAGACCCCGTAGAAA
AGATCAAAGGATCTTCTTGAGATCCTTTTTTTCTGCGCGTAATCTGCTGCTTGCAAACAA
AAAAACCACCGCTACCAGCGGTGGTTTGTTTGCCGGATCAAGAGCTACCAACTCTTTTTC
CGAAGGTAACTGGCTTCAGCAGAGCGCAGATACCAAATACTGTCCTTCTAGTGTAGCCGT
AGTTAGGCCACCACTTCAAGAACTCTGTAGCACCGCCTACATACCTCGCTCTGCTAATCC
TGTTACCAGTGGCTGCTGCCAGTGGCGATAAGTCGTGTCTTACCGGGTTGGACTCAAGAC
GATAGTTACCGGATAAGGCGCAGCGGTCGGGCTGAACGGGGGGTTCGTGCACACAGCCCA
GCTTGGAGCGAACGACCTACACCGAACTGAGATACCTACAGCGTGAGCATTGAGAAAGCG
CCACGCTTCCCGAAGGGAGAAAGGCGGACAGGTATCCGGTAAGCGGCAGGGTCGGAACAG
GAGAGCGCACGAGGGAGCTTCCAGGGGGAAACGCCTGGTATCTTTATAGTCCTGTCGGGT
TTCGCCACCTCTGACTTGAGCGTCGATTTTTGTGATGCTCGTCAGGGGGGCGGAGCCTAT
GGAAAAACGCCAGCAACGCGGCCTTTTTACGGTTCCTGGCCTTTTGCTGGCCTTTTGCTC
ACATGTTCTTTCCTGCGTTATCCCCTGATTCTGTGGATAACCGTATTACCGCCTTTGAGT
GAGCTGATACCGCTCGCCGCAGCCGAACGACCGAGCGCAGCGAGTCAGTGAGCGAGGAAT
TCGCTTGCATGCCTGCAGAATTAAAAGTTAGTGAACAAGAAAACAGTGAAGCACCAGTTT
CTGAACCAAAAGAAGACGAAAAAACAAAAAAAGATTAAGCAATTTATTTGGAAAATCTTT
TTTTGTTTTTTTAAGAAATATTTATTGTTTTTTTTAAAAAATTATTGTACAGTTGCTACT
ATAAGGGAAAGAAAAAAAGAAAGATATAAATTGTATAAAGTAGGGTTAGAAGCAATTAAT
AATTATTATTAATGTTATTTTTCTCTTATATATTCAATGTAATTTTAATTACATTTGCTT
TTAATAAAAACACTACTTAATAGAGAAAGGAAATATAAGATCTCATATGTCTAGATTAGA
TAAAAGTAAAGTGATTAACAGCGCATTAGAGCTGCTTAATGAGGTCGGAATCGAAGGTTT
AACAACCCGTAAACTCGCCCAGAAGCTAGGTGTAGAGCAGCCTACATTGTATTGGCATGT
AAAAAATAAGCGGGCTTTGCTCGACGCCTTAGCCATTGAGATGTTAGATAGGCACCATAC
TCACTTTTGCCCTTTAGAAGGGGAAAGCTGGCAAGATTTTTTACGTAATAACGCTAAAAG
TTTTAGATGTGCTTTACTAAGTCATCGCGATGGAGCAAAAGTACATTTAGGTACACGGCC
TACAGAAAAACAGTATGAAACTCTCGAAAATCAATTAGCCTTTTTATGCCAACAAGGTTT
TTCACTAGAGAATGCATTATATGCACTCAGCGCTGTGGGGCATTTTACTTTAGGTTGCGT
ATTGGAAGATCAAGAGCATCAAGTCGCTAAAGAAGAAAGGGAAACACCTACTACTGATAG
TATGCCGCCATTATTACGACAAGCTATCGAATTATTTGATCACCAAGGTGCAGAGCCAGC
CTTCTTATTCGGCCTTGAATTGATCATATGCGGATTAGAAAAACAACTTAAATGTGAAAG
TGGGTCTTAAGGATCTGAACTGCAGGTCGATggatccCCGCCTCCAGCAGAATTAAAAGT
TAGTGAACAAGAAAACAGTGAAGCACCAGTTTCTGAACCAAAAGAAGACGAAAAAACAAA
AAAAGATTAAGCAATTTATTTGGAAAATCTTTTTTTGTTTTTTTAAGAAATATTTATTGT
TTTTTTTAAAAAATTATGTACAATTGCTACTATAAGGGAAAGAAAAAAAGAAAGATATAA
ATTGTATAAAGTAGGGTTAGAAGCAATTAATAATTATTATTAATGTTATTTTTCTCTTAT
ATATTCAATGTAATTTTAATTACATTTGCTTTTAATAAAAACACTACTTAATAGAGAAAG
GAAATATAACCTAGGGTCTCCTCTTAGAGGAGTTAGCTTAGTTTTAGAGCTAGAAATAGC
AAGTTAAAATAAGGCTAGTCCGTTATCAACTTGAAAAAGTGGCACCGAGTCGGTGCTTTT
TTTACGGATCTAAATTAAAGTTGGTTCATTCAAAGTTACAATTCGTTTAAAAAGTGAATA
ATTAATTATTAATTAATTTATTAAAAAACAACCAAAAGGTTGTTTTTTATTTTTTAAAGG
TCGGTCCGTAATACGACTCACTTAAGGCCTTGACTAGAGGGTACCAATTCCAGAGTCTCC
ATATATGAATAATGGATTTCATTTTTTTCCAAGATCTAAGAAATGAAAAAACTTTTATCG
ATCAAAAACTAAAAAAAATATTGACACTCTATCATTGATAGAGTATAATTAACGGGATCC
TCTATCATTGATAGAGGGATCCCGCCAAGCTTGGGATCCCCAGCTTGTTGATACACTAAT
GCTTTTATATAGGGAAAAGGTGGTGAACTACTATGGACAAAAAATATTCTATTGGATTAG
CTATTGGAACTAATTCAGTAGGTTGAGCTGTTATTACAGATGAATATAAGGTACCATCAA
AGAAATTTAAGGTTTTAGGTAATACTGATAGACATTCAATTAAAAAAAATTTAATCGGAG
CATTACTTTTTGATTCTGGAGAAACAGCAGAAGCTACCAGATTAAAAAGAACGGCTCGTA
GACGATATACTAGACGTAAAAACAGAATCTGTTATCTTCAAGAAATTTTTAGTAATGAAA
TGGCTAAAGTTGATGACTCTTTTTTTCACAGATTGGAAGAGTCTTTCTTAGTAGAAGAAG
ATAAAAAGCATGAAAGACACCCCATCTTTGGTAATATTGTAGATGAAGTCGCCTATCATG
AAAAATATCCTACAATTTATCATTTAAGAAAAAAATTAGTAGATAGCACAGATAAAGCTG
ATTTAAGATTAATTTATTTAGCACTAGCACATATGATCAAATTTAGAGGTCACTTCTTAA
TTGAAGGTGATTTAAACCCTGATAATAGTGACGTTGATAAATTATTTATTCAATTAGTAC
AAACGTATAATCAACTTTTCGAAGAAAACCCAATTAATGCTAGTGGGGTTGATGCAAAAG
CTATCCTTTCGGCTCGTCTTTCAAAATCTAGGAGACTTGAAAACTTAATTGCACAATTAC
CGGGAGAGAAAAAAAACGGTTTATTTGGTAACTTAATCGCGTTATCTTTAGGTTTAACCC
CGAATTTTAAGAGTAATTTCGATCTAGCTGAAGATGCTAAACTACAATTATCTAAAGATA
CTTATGACGATGACTTAGATAATTTATTAGCTCAGATTGGTGATCAATATGCAGACTTAT
TTTTAGCAGCAAAAAACTTAAGCGACGCAATCTTATTGAGTGATATATTGAGAGTTAACA
CAGAAATCACTAAAGCACCATTAAGTGCAAGTATGATTAAACGTTATGATGAACACCACC
AAGATTTAACACTATTAAAAGCATTAGTTAGACAACAATTACCTGAAAAGTATAAAGAAA
TTTTCTTCGATCAAAGCAAAAATGGTTATGCTGGTTATATTGATGGTGGAGCTTCACAAG
AAGAATTTTATAAGTTCATTAAGCCTATCCTAGAAAAAATGGATGGAACAGAAGAACTAT
TAGTCAAGTTAAATCGTGAAGATTTACTACGCAAACAAAGAACTTTTGATAATGGTAGCA
TTCCTCATCAAATTCACTTAGGAGAACTACACGCTATCCTAAGAAGACAAGAAGATTTTT
ATCCTTTTTTAAAAGATAATAGAGAAAAAATTGAAAAAATCTTAACATTTAGAATCCCTT
ACTATGTAGGTCCGTTAGCTAGAGGAAATAGTAGATTTGCATGAATGACTCGAAAATCAG
AAGAGACTATCACACCATGAAATTTTGAGGAAGTTGTGGATAAAGGTGCATCTGCGCAAT
CTTTTATTGAGCGAATGACTAATTTCGATAAGAACTTACCTAATGAAAAAGTATTACCTA

AGCACTCATTATTATATGAATACTTTACTGTTTATAACGAACTTACTAAAGTAAAATATG
TTACCGAAGGAATGAGAAAACCAGCGTTCCTAAGTGGAGAACAAAAGAAGGCTATTGTTG
ATTTATTATTTAAGACAAATAGAAAAGTAACTGTAAAACAACTAAAAGAAGATTATTTTA
AAAAAATTGAATGTTTTGATTCAGTCGAAATTTCTGGAGTTGAAGACCGTTTCAACGCAA
GTTTAGGCACTTACCACGATCTACTAAAAATTATTAAAGATAAAGATTTTCTTGATAACG
AAGAAAATGAAGACATTCTAGAAGATATTGTCCTAACTTTAACTTTATTCGAAGACAGAG
AAATGATTGAAGAAAGATTAAAAACTTACGCTCACTTATTTGATGATAAAGTTATGAAGC
AGTTGAAGCGCCGACGATATACCGGTTGAGGTAGACTCTCAAGAAAGCTAATCAATGGTA
TTAGAGACAAACAATCAGGTAAAACAATTTTAGATTTTTTAAAAAGCGACGGATTTGCTA
ATAGAAACTTCATGCAATTGATCCACGATGATT
